# Probing Critical Injury Thresholds for Maladaptive Epithelial Injury and Repair Processes with Photoresponsive Bioinspired Synthetic Basement Membrane

**DOI:** 10.64898/2025.12.24.695213

**Authors:** Lina Pradhan, Bryan P. Sutherland, Samantha L. Swedzinski, Kartik Bomb, Qi Zhang, Samantha E. Cassel, Catherine A. Fromen, April M. Kloxin

## Abstract

Microinjuries to the lung epithelium are hypothesized to initiate maladaptive processes that lead to fibrosis. Human *in vitro* models remain a great need for studying this injury-initiation process for mechanistic understanding and therapeutic development. We established a photoresponsive synthetic extracellular matrix (ECM) inspired by the basement membrane that enables triggered ‘injuries’ of defined size and frequency for probing cellular responses. The synthetic matrix integrated a photolabile *bis*-coumarin linker for light-triggered ‘injury’ and relevant integrin-binding peptides for cell function. Bio-orthogonal chemistry was used to create hydrogel-based ECMs with tunable elasticity in transwells, which are traditionally used for epithelial cell culture. Integrin-binding peptide combinations synergistically promoted model epithelial cell layer formation with increased E-cadherin expression and barrier function. An accessible photomasking approach was established for selectively photodegrading the synthetic matrix with cytocompatible visible light and achieving different injury depths and widths. Following a ‘critical’ injury size, cell responses recapitulated key features of dysregulated re-epithelialization with decreased E-cadherin, proliferation, and barrier function and increased apoptosis. This work provides a new materials-based tool for probing injury and repair processes with tunable control of both the ECM and injury to it with opportunities for future mechanistic and therapeutic insights to address maladaptive wound healing processes.

## **1.** Introduction

Fibrosis affects almost every tissue in the body, including pulmonary, dermal, ocular, and cardiac tissues, and is the pathological outcome of misregulated wound healing or chronic inflammation^[1]^. Fibrotic diseases are defined by the accumulation of excess fibrous connective tissue, causing scarring and organ malfunction^[2]^ and accounting for 17% of deaths worldwide^[3]^. For example, lung fibrosis particularly idiopathic pulmonary fibrosis (IPF) is incurable and often fatal^[4–7]^. IPF amongst other fibrotic diseases is thought to be initiated by repeated micro-injuries to the epithelium and accelerated by the persistence of activated fibroblasts, leading to accumulation of extracellular matrix (ECM) proteins, tissue stiffening, and uncontrolled wound healing with an eventual decline in lung function^[7–10]^. However, the human disease is often caught in advanced stages, and its exact origins and mechanism accordingly are not fully understood or captured by existing model systems, impeding the development of effective therapeutics^[11–12]^. Such imperfect wound healing is increasingly recognized as a precursor to a range of chronic diseases, yet little often is known about the initiation of the underlying maladaptive processes^[13]^. Relevant human culture model systems for studying the initiation and early development of human fibrotic diseases remain a great need for probing mechanism and toward the development of more effective therapies^[14–16]^.

While many preclinical animal models have provided valuable insights and helped to identify therapeutic targets^[17–19]^, these models often lack specificity regarding epithelial injury location, affected cell types, and the ability to study repeated injury, particularly in the context of internal epithelial tissues such as found in the lung^[20–21]^. Toward addressing this gap, several *in vitro* models have been developed for studying epithelial injury and repair processes. Most studies have employed the gold-standard ‘scratch’ assay in two-dimensional (2D) culture on tissue culture plastic: epithelial cells are cultured on a plastic plate or a transwell membrane in submerged conditions, and upon introducing a scratch to the cell layer, the effects of cell-microenvironment interactions on cell migration and proliferation during wound healing are probed^[22–24]^. A scratch can be introduced by scraping a tool (e.g., pipette tip) across the plastic culture surface, removing a barrier between seeded cells, or selectively dislodging cells from the surface using a laser or microjet, each providing different levels of uniformity and sizes of ‘injury’^[25–28]^. With the traditional scratch model, linear wound closure dynamics have been observed within 12 h for a variety of cell types, and drugs for influencing wound healing velocity evaluated (e.g., microtubule inhibitors that impact cell motility)^[29]^. To further probe cell-cell interactions, stromal cells have been integrated: for example, model lung alveolar epithelial cells (A549) have been cultured in a transwell insert while fibroblast cells are cultured in the bottom chamber, demonstrating that fibroblast cells from healthy versus IPF tissue origins stimulate a differential epithelial repair responses after scratch injury. While such systems model aspects of cell wound healing responses, the biophysical and biochemical properties of the tissue microenvironment, particularly the ECM, typically are not represented^[11]^. Further, while existing scratch methods afford different levels of spatiotemporal precision, their approach to injury is non-specific in nature and does not capture the underlying damage that occurs to tissues and the ECM upon injury.

Engineered materials have been integrated within dynamic model systems to provide different levels of microenvironment property control and probe cellular responses to stimuli in more physiologically relevant *in vitro* cultures^[30]^. For example, poly(dimethylsiloxane) (PDMS)-based systems have been created that enable real-time probing of cell responses to dynamic stiffening in harvested collagen I ECMs modulated with magnetomechanical cantilevers in three-dimensional (3D) culture^[31–32]^. Such approaches have been used to create membranous lung-mimicking microtissues that integrate epithelial cells and have shown utility for evaluation of anti-fibrotic therapies^[33]^. Further, silk-collagen type I matrices have been integrated within bioreactors for co-culturing lung cells to reproduce pulmonary fibrotic tissue pathology and shown feasibility for probing epithelial injury response to small molecules (e.g., bleomycin) and modulation of cellular responses with anti-fibrotic drugs^[34]^. Use of light-based chemistries within engineered systems provides further opportunities for probing dynamics in both space and time. For example, polymer-peptide synthetic ECMs have been stiffened *in vitro* using photopolymerization during 3D culture of lung epithelial cells^[35]^, and *ex vivo* mouse and human lung tissues have been stiffened using visible-light-mediated photochemistry to mimic early fibrotic lesions and epithelial cell differentiation and remodeling processes^[14]^. Photolabile chemistries that respond to cytocompatible doses of long wavelength UV and visible light, such as coumarin and nitrobenzyl groups^[36–37]^, also have been used as linkers within synthetic ECMs for matrix degradation upon the application of light and have shown utility for the development of organoid models of the lung and gut in 3D culture^[38–40]^. Overall, these works demonstrate the utility of dynamic engineered materials systems in studies of cell responses to fibrotic processes and evaluation of therapeutic targets and strategies. The precise control that synthetic ECMs and light-based chemistries afford provides opportunities for now probing the earliest initiation of such processes upon triggered epithelial injury.

Human *in vitro* model systems remain a need for studying epithelial cell responses to wound healing events, from relevant basement membrane interactions that support epithelium formation and maintain phenotypes under healthy conditions to controlled injury to enable probing of wound repair and maladaptive processes that lead to fibrotic disease. To address this need, we established a well-defined photoresponsive synthetic ECM inspired by the basement membrane that allows precise light-triggered ‘injuries’ of defined size and frequency to an epithelial monolayer, using model lung epithelial cells (A549 and Calu-3) as prototypical examples. This hydrogel-based synthetic ECM was constructed using bio-orthogonal chemistry with a photolabile poly(ethylene glycol) (PEG)-*bis*-coumarin-azide (PEG-2-CmPN_3_) crosslinked with PEG-*tetra*-bicyclononyne (PEG-4-BCN), imparting visible-light degradability while enabling control of the initial stiffness of the matrix with crosslink density. Further, this chemistry allowed facile integration of integrin-binding peptide combinations (laminin inspired AG73 and fibronectin inspired PHSRN-RGDSP (PHSRN)) that synergistically promoted epithelial layer formation with enhanced expression of an adherens junction protein E-cadherin and epithelial barrier function. The tunable biophysical and biochemical properties of this synthetic ECM, combined with formation in a transwell insert and an accessible photomasking strategy, allowed selective photodegradation of the basement membrane mimic to achieve different injury depths and widths to a model epithelial cell layer. Using this platform, we observed that, when injury size surpassed a critical threshold, epithelial cells exhibited hallmark features of dysregulated re-epithelialization, including decreased E-cadherin expression, reduced proliferation, impaired barrier function, and increased apoptosis. Repeated microinjuries further suppressed growth and recovery, resulting in global epithelial impairment and failure to repair wounds. Together, these studies demonstrate that the size and repetition of epithelial injury are key determinants of repair outcomes and establish a new materials-based tool for probing mechanisms of maladaptive wound healing and fibrosis initiation with future opportunities for use in therapeutic discovery.

## 2. Results

### 2.1. Synthetic basement membrane for photo-injury model

We first aimed to formulate a photodegradable bioinspired synthetic ECM relevant for studying lung epithelial cell response to injuries. The hydrogel-based synthetic ECM was formed by reacting PEG-4-BCN (exo) crosslinker with photodegradable CmPN_3_ *bis*-linker via strain promoted azide-alkyne cycloaddition (SPAAC) (Figure 1A), allowing the formation of a hydrogel-based synthetic ECM with tailored biophysical and biochemical properties while maintaining the integrity of the photolabile linker (Figure S1A). The BCN (exo) and CmPN_3_ functional groups were selected from amongst other SPAAC reactive handles to give sufficient working time prior to gelation for sample preparation (∼ 12.5 minutes) while achieving complete gelation within hours (Figure S1B). For promoting cell attachment and function, we integrated 1 mM of each bioinspired peptide within the hydrogel precursor solution: here, laminin inspired K(N_3_)-GWG-RKRLQVQLSIRT (AG73, binding domain within laminin 111 for heparan sulfate and syndecan) and fibronectin inspired K(N_3_)-GWG-PHSRNG_10_RGDSPG (PHSRN, binding domain within FnIII9’10 for α5β1)^[41–44]^, where each peptide has shown relevance in the adhesion and phenotype of other types of epithelial cells^[45]^. We hypothesized that together these peptides would be synergistic in promoting model lung epithelial cell adhesion and function for epithelial barrier formation given the importance of these cues in lung development and regeneration^[46–48]^. The weight percentage of the total PEG monomers crosslinked to form the synthetic ECM was then varied to achieve moduli inspired by healthy to diseased lung tissue, as measured with rheometry. We identified equilibrium swollen synthetic ECM formulations inspired by healthy (Young’s modulus (E) ∼2 kPa) and fibrotic lung tissue elasticity (∼10 kPa) (Figure 1B, S1).

**Figure 1.**
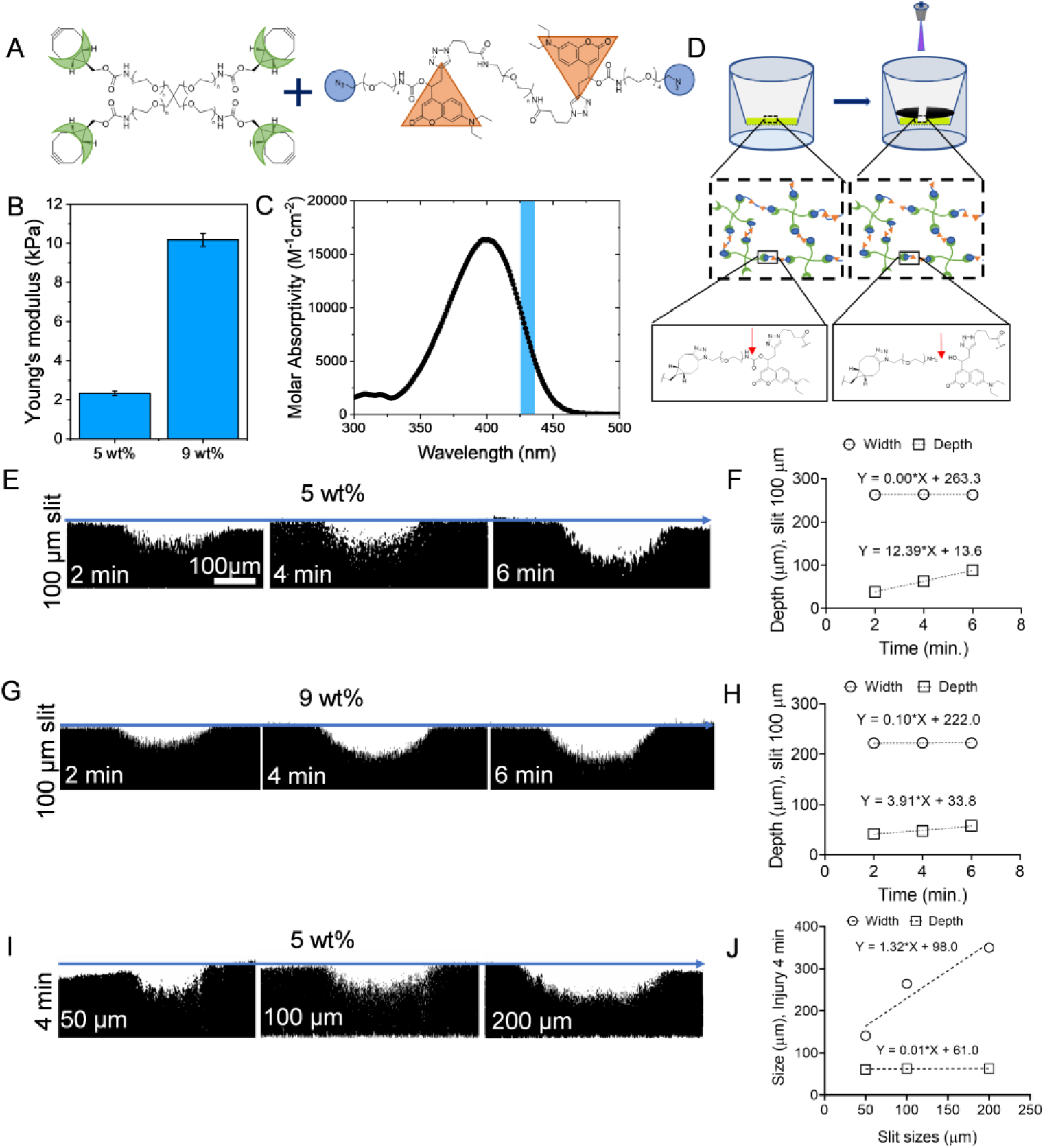
Design of synthetic basement membrane for photo-injury (PI) model system. A) Hydrogel-based synthetic ECM was formed by strain-promoted azide-alkyne cycloaddition (SPAAC) between four-arm PEG-*tetra*-bicyclononyne (PEG-4-BCN) and linear PEG-*bis*-coumarin-azide (PEG-2-CmPN_3_). B) Young’s modulus (kPa) of synthetic ECM inspired by healthy (∼5 wt%) (‘soft’) and diseased (∼9 wt%) (‘stiff’) lung tissues. C) Absorbance Spectra of CmPN_3_ and irradiation wavelength (highlighted area centered at 430 nm light). D) Approach to create injury to synthetic ECM via degradation of CmPN_3_ upon irradiation with 430 nm light. E) Representative images (binary orthogonal x-y projections from confocal z-stacks) of selective degradation of the surface using photomasking with different irradiation times and 100 µm slit on 5 wt% hydrogel. F) Quantitative analysis from (E). G) Different irradiation times with 100 µm slit on 9 wt% hydrogel. H) Quantitative analysis from (G). I) Different photomask ‘slit’ widths on 5 wt% hydrogels. J) Quantitative analysis of images (I). Data shown represent mean ± SD (n=3), where the error bars are smaller than the data points, and equations shown represent linear fits of these data.

The CmPN_3_ *bis*-linker included in these hydrogels then was used to selectively degrade the synthetic ECM with visible light to mimic injury. Importantly, the absorbance spectra of the crosslinker, CmPN_3_ (Figure 1C), spanned the UV to visible range such that the hydrogel readily degraded with low cytocompatible doses of light of wavelengths in the range of ∼350– 475 nm. Tuning in this range can be used to modulate the rate and mode of degradation: rapid degradation is achieved by irradiation with a highly-absorbed wavelength (e.g., 430 nm) along with surface erosion for thick gels (e.g., 0.5 mm), whereas slower degradation of the bulk is achieved by irradiation with a less-absorbed wavelength (e.g., 455 nm) (Figure S2). Accordingly, to simulate injuries, we selected a light source centered at 430 nm to rapidly erode regions of interest on the surface of the synthetic ECM, where upon irradiation the coumarin group degrades to a coumarin methyl ester cleaving the linker within the network (Figure 1D).

Selective degradation of these hydrogels was achieved using a simple, accessible photomasking approach. A photomask patterned with features of interest was held close to the surface of the synthetic ECM with custom 3D-printed, modular apparatus. The system consists of multiple interlocking components: their design enables *i)* sterile handling, *ii)* use with transwells, and *iii)* close proximity of the photomask to surface without disrupting the attached cell layer during culture while blocking light from transmission to neighboring samples in adjacent wells (Figure S3, Video S1). Here, the photomask had a clear line surrounded by a black background, producing a ‘slit’ through which light was transmitted (Figure S4) and varied in size to change the width of the injury that was created upon photodegradation of the synthetic ECM. Uniform layers of the synthetic ECM labeled with a fluorophore (azido-AF647) were formed within transwell inserts and imaged with confocal microscopy before and after irradiation (6.75 mW*cm^-2^ at 430 nm) (Figure S5). Orthogonal images of the confocal z-stacks enabled visualization and quantification of the degraded area on either a ‘soft’ (5 wt%) or a ‘stiff’ (9 wt%) matrix (Figure 1E-J). Within the conditions probed, the width of the ‘slit’ on the photomask does not affect the depth of the injury (Figure 1J). While the weight percent of the hydrogel does not impact the initial depth of the injury (similar magnitude with 2 min irradiation), differences in injury depth are observed with increased irradiation times (Figure 1F, H), owing to differences in photolabile group concentration and crosslink density between these compositions^[49]^. Irradiation time was found to be an effective way to control the depth of degradation, and a linear correlation between irradiation time and injury depth was observed, supporting the surface erosion mechanism at this irradiation wavelength and sample thickness (erosion rate of ∼ 12 μm/min or 4 μm/min for 5 or 9 wt% hydrogel). Overall, design parameters and irradiation times were established for using low intensity visible light to create an ‘injury’ of different depths or widths to the synthetic ECM.

### 2.2. Development of epithelium on photoresponsive synthetic ECM

Next, we evaluated the effects of biochemical cues for developing a model ‘healthy’ epithelium on the photoresponsive synthetic ECM (Figure 2A): fibronectin inspired PHSRN and laminin inspired AG73 individually or combined. Initial attachment of A549 cells after 24 h was quantified using brightfield images (Figure S6A). Cell coverage of the surface over time was assessed based on staining cells at different culture times with a live/dead membrane integrity assay and quantifying the percentage of the area that was Calcein positive with confocal microscopy (Figure 2B, Figure S7A). A synergistic effect was observed with an extensive coverage of the surfaces by A549s in the PHSRN+AG73 condition, compared to the PHSRN or AG73 conditions (Figure 2C), while cell viability over time was high and similar amongst conditions (Figure S7B). Further, the combination of PHSRN+AG73 peptides led to a significant enhancement in initial cell attachment, compared to PHSRN or AG73 alone (Figure S6B). Notably, the PHSRN+AG73 condition had >90% surface coverage with the formation of a monolayer on day 10 in culture, whereas surface coverage remained <80% on PHSRN or AG73 conditions. To probe this effect more broadly, another model human lung airway epithelial cell line (Calu-3) was cultured on the different synthetic ECMs. Similar trends were observed for Calu-3 cells where a combination of peptides (PHSRN+AG73) showed significant monolayer surface coverage over time and initial cell attachment, compared to the single peptide conditions (Figure S8, S9). Overall, these observations of model lung epithelial cell attachment and monolayer formation supported the relevance of the designed synthetic ECM inspired by basement membrane for promoting desired cellular functions and potential for further studies of cell phenotype and response to injury. To characterize cell phenotype within the formed monolayer, additional samples were stained for *i)* the epithelial marker E-cadherin, an adherens junction protein expressed during epithelial cell growth that is integral in cell adhesion and maintains epithelial phenotype, and *ii)* the mesenchymal marker Vimentin, expressed in the cell cytoskeletal and a marker of mesenchymal phenotype^[50–51]^, and imaged with confocal microscopy. Notably, a significant number of E-cadherin positive cells were observed on PHSRN+AG73 (Figure 2D, E, Figure S10), compared to PHSRN or AG73 alone, while E-cadherin protein expression increased over time in all conditions. Correspondingly, a significantly lower number of Vimentin positive cells was observed on PHSRN+AG73 over time, compared to PHSRN or AG73 alone (Figure S11), indicating maintenance of epithelial phenotype. Additionally, Calu-3s exhibited a statistically higher number of E-cadherin positive cells on PHSRN+AG73 or PHSRN, compared to AG73, with limited expression of Vimentin in any of the conditions (Figure S12, S13), suggesting epithelial phenotypes with the increased importance of PHSRN relative to AG73. These exciting observations of robust E-cadherin expression on the designed synthetic ECM further supported maintenance of epithelial properties in addition to monolayer formation and the potential for a relevant *in vitro* system for further wound healing studies.

**Figure 2.**
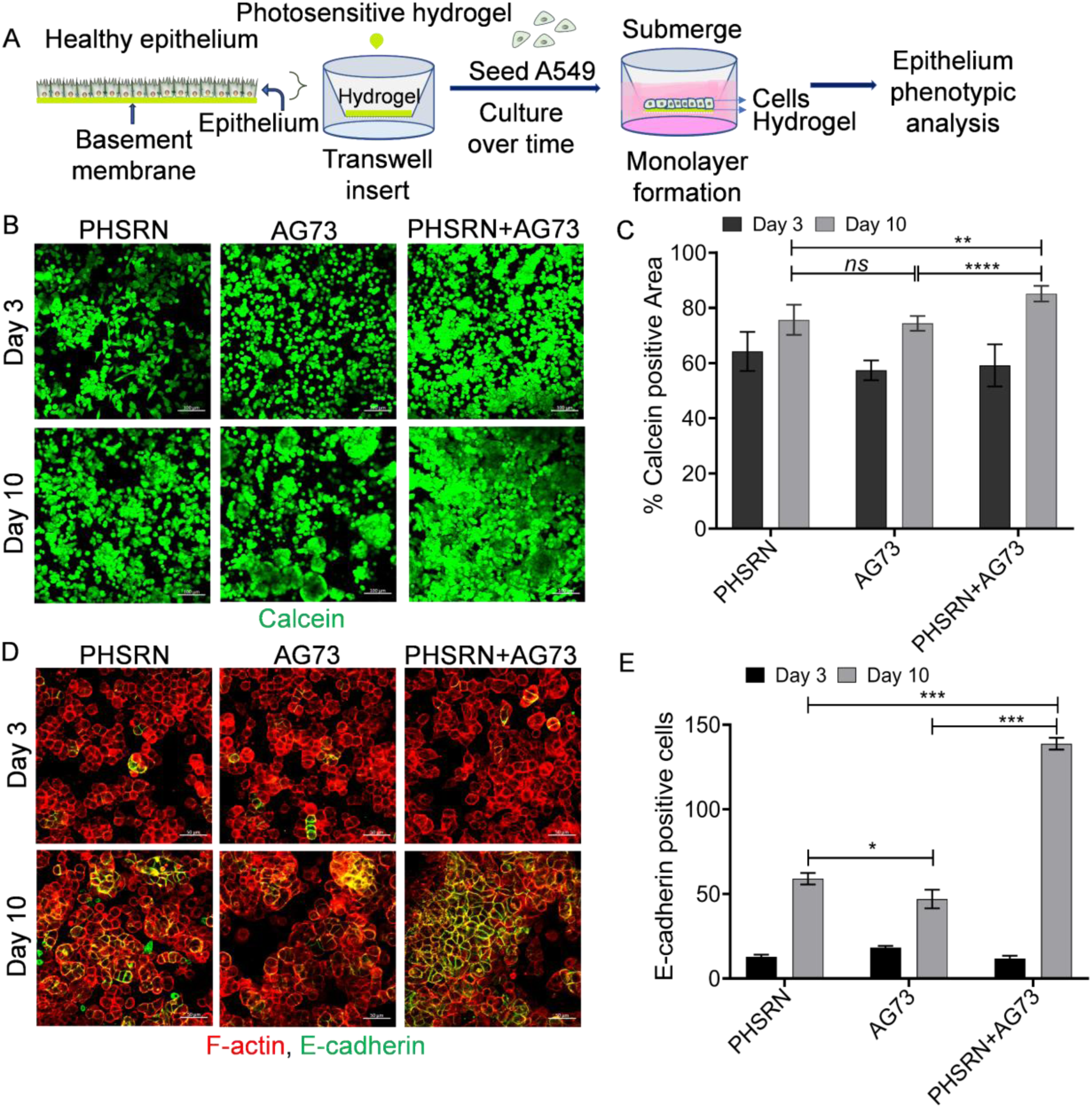
Effect of ECM compositions on model lung epithelial cells on ‘soft’ hydrogels inspired by healthy tissue. A) Schematic approach for development of a ‘healthy’ epithelium on synthetic photosensitive hydrogel with integrated receptor-binding peptides and phenotypic analysis. B) Representative images for assessment of epithelial cell monolayer formation with A549 cells on different synthetic ECM compositions using Calcein (dye for marking live cells), where increased % Calcein positive area was used to assess monolayer formation over time on PHSRN, AG73, or PHSRN+AG73 condition (confocal z-projection [image frame size of 1024 × 1024, z-stack <10 µm with ∼2 µm slice interval], green (Live cells) stained with Calcein, and red (dead cells, shown Figure S7) stained with ethidium homodimer-1 (Ethi-1), Scale bar = 100 µm). C) Quantitative analysis of Calcein positive area of (B). Significant differences between conditions at timepoints of interest were assessed by Student’s two-sided t-test (*p < 0.05, **p < 0.01; ***p < 0.001). D) Representative images of immunofluorescence demonstrate functional monolayer formation of A549 (confocal z-projection, red stained with (F-actin), green (E-cadherin), Scale bar = 50 µm). E) Quantitative analysis of E-cadherin positive cells of (D). Significant differences assessed by Student’s two-sided t-test, where differences shown for comparison between conditions (*p < 0.05, **p < 0.01; ***p < 0.001). Data shown represent mean ± SD (n=3).

### 2.3. *In vitro* injury model for real-time monitoring of epithelial cell response

To investigate monolayer formation and then response to injury in real-time, we focused on the PHSRN+AG73 synthetic ECM and deployed a stable reporter A549 cell line, which was engineered to express a red fluorescent protein constitutively and conditionally express a green fluorescent protein when alpha smooth muscle actin (αSMA) was upregulated^[52]^. Cells were seeded on the synthetic ECM within the transwell insert and monitored over time by confocal microscopy (Figure 3A, B). Greater than 90% surface coverage was observed on day 10 in culture with limited expression of αSMA, consistent with our earlier observations with end point assays and supporting an epithelial-like phenotype (Figure 3C, D, Figure S14).

**Figure 3.**
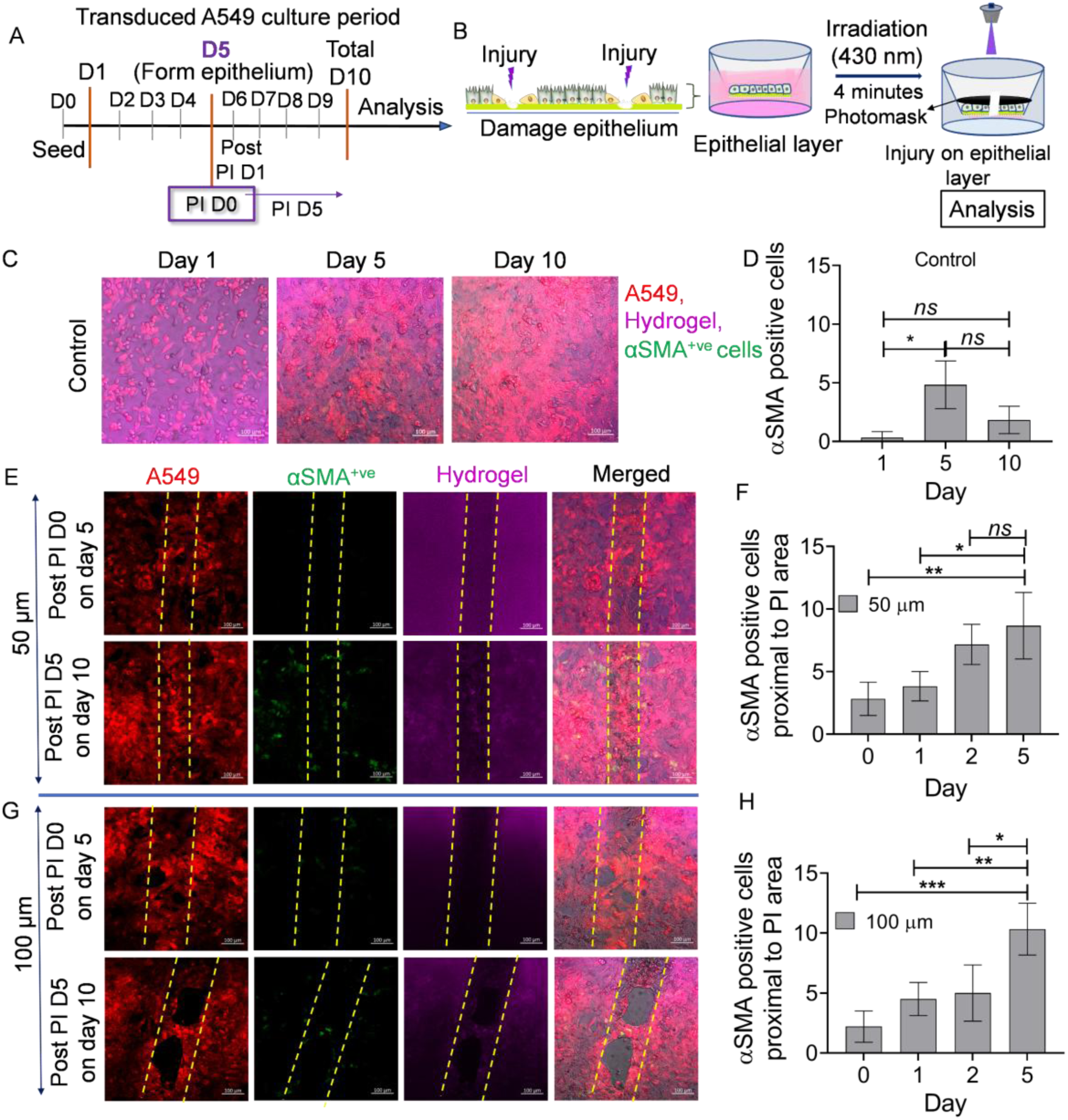
Real-time analysis of epithelial monolayer formation and response to injury over time. A) Representation of culture period prior to and after photo-injury (PI) on model epithelium. Here, A549 cells were used that had been transduced for constitutive expression of a red fluorescent protein, allowing visualization of cells, and conditional expression of a green fluorescent protein when alpha smooth muscle actin (αSMA) expression was upregulated, allowing assessment of cells exhibiting a mesenchymal-like wound healing phenotype. B) Approach for light-triggered ‘injury’ to induce damage to synthetic matrix underlying the epithelial monolayer. C) Representative images of epithelium formation with A549 cells on the PHSRN+AG73 synthetic matrix over time (confocal z-projection, red channel (A549) shows all transduced cells in the growth area, green channel (αSMA^+ve^) shows cells expressing αSMA, and far-red channel (hydrogel) shows the hydrogel-based synthetic ECM, Scale bar = 100 µm). D) Quantitative analysis of (C). E) Representative images of real-time epithelial cell recovery after PI (50 µm slit) (confocal z-projection, red channel (A549 fluorescent reporter), green channel (αSMA^+ve^ fluorescent reporter), far red channel (hydrogel) where injury area is visible with loss of intensity and boundaries of the injury are noted with dashed lines, Scale bar = 100 µm). F) Quantitative analysis of (E). G) Representative images of epithelial cell recovery after PI (100 µm slit) (confocal z-projection, Scale bar = 100 µm). H) Quantitative analysis of (G). Significant differences over time assessed by Student’s two-sided t-test, where differences shown for comparison between time points (*p < 0.05, **p < 0.01; ***p < 0.001). Data are shown represent mean ± SD (n=3).

Next, we developed an approach for controlled epithelium injury utilizing the photodegradability of the synthetic ECM and assessed the effects of epithelial cell response. Here, ‘injury’ to the ECM was triggered upon irradiation with low cytocompatible doses of 430 nm light. Epithelial injuries of different sizes (widths) were induced with different photomask ‘slit’ widths while using the same dose of light (6.75 mW*cm^-2^ at 430 nm for 4 minutes) for creating similar ‘injury’ depths amongst all conditions. As described above, a photomask with 3D printed holder was used to avoid disturbing the cell monolayer (Figure S3, Video S1). Differential responses of cells for small versus large injury sizes (50 µm versus 100 or 200 µm ‘slit’ condition) were observed in real-time brightfield imaging overnight after injury (Video S2, Figure S15). To further probe and understand mechanism, we examined healing *in situ* 5 days after injury using the reporter cell line.

A significant upregulation of expression of αSMA was observed over time proximal (within the same field of view) to the photo-injury (PI) for injury widths of 50 and 100 µm (Figure 3E, H, Figure S16A, S17A). Further, the expression of αSMA remained significantly higher over time in regions of the monolayer that were far from the injury area (distal, approximately 3 fields of view from the injury) (day 5 post PI compared to day 0 post PI) (Figure S16B, C, 17B, C). Interestingly, with the larger injury width created by a 100 µm size of ‘slit’ photomask, cells did not fill the gap to ‘close’ the injury over the experimental time course (over 5 days), whereas cells filled the gap to ‘close’ the injury with the smaller width created by a 50 µm ‘slit’ photomask. This differential healing was correlated with a significant number of cells upregulating αSMA expression distal to the larger injury compared to the smaller injury (Figure S17D), suggesting mesenchymal phenotype enhancement with increased injury width. Note, no effects were observed for irradiation alone, when light was applied to a cell monolayer on a non-degradable synthetic ECM (Figure S18), or with degradation products alone, where a photodegradable synthetic ECM was eroded with light and degradation products incubated with cells in 2D plate culture (Figure S19). Overall, these findings indicated that many A549 cells maintained an epithelial phenotype with a minimum expression of αSMA during a healthy epithelium formation, whereas many A549 cells exhibited a mesenchymal phenotype with a significant number of αSMA expressing cells upon epithelium injury. These observations support the global phenotypic loss of epithelial-like cells during injury that may influence maladaptive healing processes and contribute to disease processes depending on injury size.

To benchmark responses against those observed with a traditional ‘scratch’ approach, we cultured reporter A549 cells on the synthetic ECM with PHSRN+AG73 and introduced a scratch to the epithelial layer on day 5 in culture using a 20-µL sterile plastic pipette tip (Figure S20).

Injury closure over another 5 days (total of 10 days in culture) then was examined. At 5 days after the scratch, we observed the original injury area was covered by epithelial cells, with more cells expressing αSMA in the scratch area for day 5 compared to day 0 post scratch (Figure S20A, B). Additionally, an increased in number of cells expressing αSMA was observed distal from the scratch area over time (Figure S20C, D). These similarities between cellular responses on the synthetic ECM to an injury, whether by physical scratching of the epithelial monolayer or light-triggered degradation of the ECM that underlies the monolayer, support the relevance of the photo-injury model system for studies of acute injury and wound healing responses. Importantly, the photo-triggered injury non-invasively allows control of injury size, where differential healing responses to ‘critically-sized’ injuries can be observed, while the synthetic ECM provides control of the initial biophysical and biochemical properties of the matrix.

### 2.4. Epithelial injury leads to global loss of phenotypic marker expression

To further understand epithelial cell response to injury, we cultured non-transduced A549 cells on the synthetic ECM and characterized phenotype by immunofluorescence (Figure 4A). Different injury widths were created on day 5 in culture using 50, 100, or 200 µm slits, while producing similar depths of injury with 4 minutes of irradiation (6.75 mW*cm^-2^ at 430 nm) for all the conditions. On day 10, A549 cells on samples with and without (control) photo-injury were stained for phenotypic markers. Here, we examined *i)* epithelial marker E-cadherin, whose loss is not only associated with but causal for a range of lung diseases, and *ii)* mesenchymal marker Vimentin, which is required for wound repair after lung injury owing in part to its important function in cell migration and proliferation^[53–56]^.

**Figure 4.**
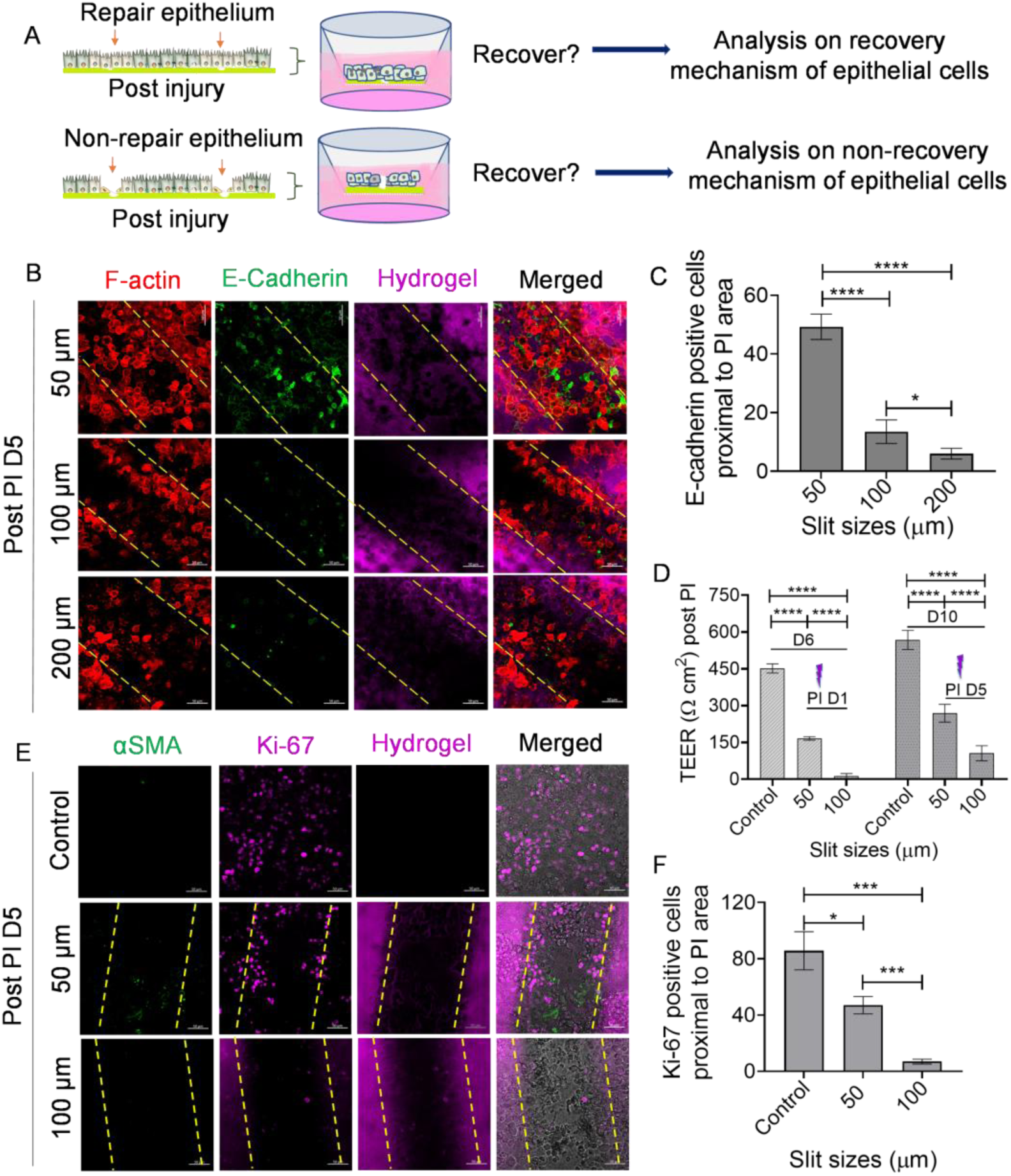
Large injury that does not heal leads to loss of E-cadherin and Ki-67 protein expression. A) Schematic representation for repair and non-repair of epithelium on synthetic photosensitive ECM and phenotypic analysis, where cellular responses are probed at 5 days post photo-injury (PI D5). B) Representative images of non-reporter A549 showing different responses after creating of different sizes of PI in comparison to controls (confocal z-projection, Day 5 after injury, F-actin (red), E-cadherin (green), hydrogel (far-red) for visualization of ‘injury’ area observed with loss of intensity, boundaries of ‘injury’ noted with dashed lines, Scale bar = 50 µm). All channels and images of control samples shown in Figure S22. C) Quantitative analysis of cells positive for E-cadherin protein in (B). D) Transepithelial electric resistance (TEER) before and after PI for samples in (B). E) Representative images of immunofluorescent staining of proliferative (Ki-67 positive, magenta) and mesenchymal (αSMA, green) markers; synthetic ECM labeled with fluorophore AF647 for visualization of ‘injury’ area observed with loss of intensity and boundaries of ‘injury’ noted with dashed lines (confocal z-projection, Scale bar = 50 µm). All channels and images of control samples shown in Figure S23. F) Quantitative analysis of cells positive for Ki-67 in (E). Statistical differences between conditions at timepoints of interest were determined Student’s two-sided t-test (*p < 0.05, **p < 0.01; ***p < 0.001). Data shown represent mean ± SD (n=3).

Distinct localization of E-cadherin positive cells was observed proximal to a small injury, created a 50 µm slit (Figure 4B). On the other hand, few E-cadherin positive cells were observed proximal to larger injuries, created using 100 and 200 µm slits (Figure 4C), suggesting loss of cell-cell communication when a larger ‘wound’ is created in the epithelium. Additionally, the number of Vimentin-positive cells was significantly increased with a small injury (using 50 µm slit) relative to larger injuries (using 100 and 200 µm slit) (Figure S21A, B), suggesting a potential mechanism that underlies the successful closure of small versus large injuries.

To further probe injury effects on the epithelium, transepithelial electrical resistance (TEER) measurements of the epithelial layer were conducted using a commercial system with the chopstick electrodes method (EVOM2, World Precision Instruments)^[57]^. We evaluated TEER values on the monolayer of A549 over time, which exhibited epithelial resistance >500 Ohm cm^2^, suggesting healthy epithelium formation on photoresponsive synthetic ECM. In contrast, TEER values were significantly decreased to <200 Ohm cm^2^ on the damaged epithelium, created by photo-injury on day 6 (PI D1). Notably, the 100 µm slit condition had a TEER value <20 Ohm cm^2^, which is similar to a bare transwell membrane and suggests complete loss of epithelial barrier integrity. We continued the culture after injury for 5 days (PI D5; day 10 in culture), followed by the measurement of epithelial resistance, where TEER values slightly increased in both 50 and 100 µm slit injury conditions compared to PI D1. Importantly, a significantly higher TEER value was detected on the injured epithelium with 50 µm slit on PI D5 compared to either *i)* 100 µm slit on PI D5 or *ii)* 50 µm slit on PI D1, suggesting progression in healing of the model epithelium (Figure 4D).

Overall, these observations of loss of E-cadherin and differential expression of vimentin with the PI model system are consistent with *in vitro* and *in vivo* observations of lung injury and disease, supporting model system validity. Further, the photoinjury system allows probing of the effects of injury size on phenotype and function. A smaller injury that ‘heals’ has a mixture of E-cadherin and Vimentin positive cells and maintenance of barrier integrity. In contrast, a larger injury that does not ‘heal’ in the experiment time frame has a loss of E-cadherin positive cells, few Vimentin positive cells, and loss of barrier function.

To investigate phenotype within the broader cell population after injury, the expression of E-cadherin protein was analyzed distal to the site of injury. A significant decrease in the number of E-cadherin positive cells was detected distal to the injury site for all injury sizes probed (50, 100, or 200 µm slit) 5 days after PI compared to control. Additionally, a significantly lower number of E-cadherin positive cells was noted with the largest injury (200 µm slit condition) compared to the two smaller injuries (50 or 100 µm slit conditions) (Figure S22A, B). Taken together, these results suggested that injury of any size affects both the population at the wounded area and in the broader epithelial population.

To further probe the wound healing process upon injury, additional samples were immunostained for cell proliferation (Ki-67) and other mesenchymal markers (αSMA). Here, we focused on 50 and 100 µm slit conditions given the differential responses observed between the resulting injury widths. A large number of Ki-67-positive and αSMA-negative cells were analyzed on control samples, suggesting a proliferative stage of A549 cells without mesenchymal phenotypes on the ‘healthy’ synthetic ECM condition; these observations with immunostaining are consistent with the observations *in situ* using the reporter cell line as noted above. After injury (day 5 post PI), the number of Ki-67-positive cells decreased proximal to the injury area with both the 50 and 100 µm conditions, where a significant decrease in Ki-67-positive cells was noted in 100 µm slit condition compared to 50 µm slit condition (Figure 4E, F). Additionally, a few αSMA-positive cells were observed proximal to the 50 µm injury area but not the 100 µm injury area (Figure S23). These results suggested that A549 cells can proliferate and undergo the epithelial-to-mesenchymal transition associated with cell migration for healing when a small injury occurs on the epithelium.

To investigate epithelial cell apoptosis and necrosis after injury, cells were stained for an apoptotic marker (Apo-15), and necrotic dead cells were detected with ethidium homodimer-1 (Ethi-1), both before and after injury. By day 5 after injury, apoptotic and necrotic cells were found proximal to the injury, both within the injury and the area that surrounds it (Figure 5A). Notably, a significant increase in both Ethi-1 (necrotic) or Apo-15 (apoptotic) populations was observed at the injury area compared to control; additionally, a significantly increased number of Apo-15 and Ethi-1 positive cells were observed at the injury created with 100 µm slit size compared to the small injury created with the 50 µm slit size (Figure 5B). Further, a higher population of Ethi-1-positive cells was observed relative to Apo-15 positive cells, suggesting that most of the cells entered necrosis phase rather than apoptosis after injury. These observations within the model system are consistent with observations in injured and diseased lung tissue^[58–59]^.

**Figure 5.**
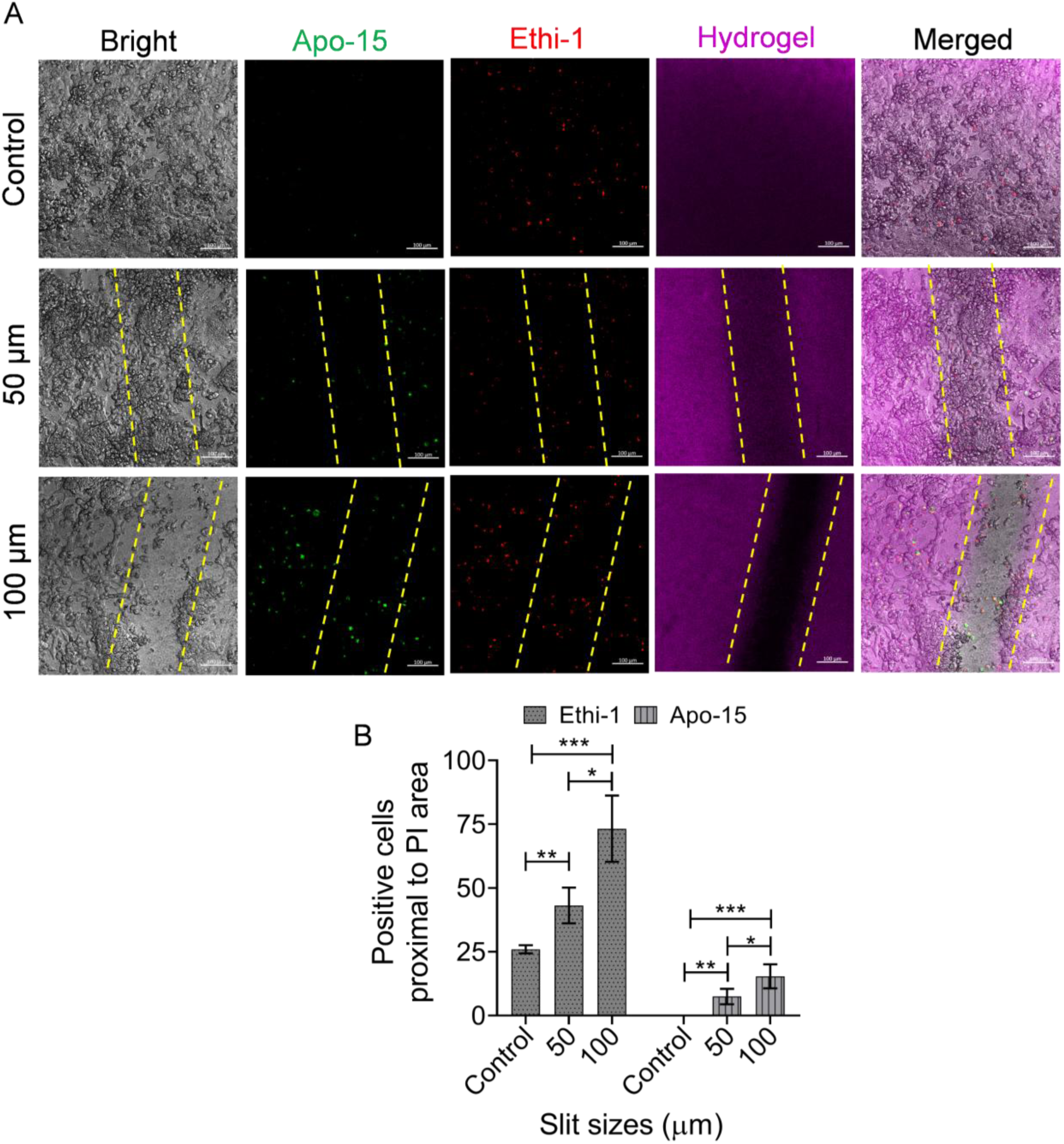
Injury and impaired healing lead to cell death processes. A) Representative images of A549 cells on day 10 (post PI D5) (confocal z-projection, Apo-15 (apoptotic cell death, green), ethidium homodimer-1 (Ethi-1, necrotic dead cells, red), hydrogel labeled with AF647 and injury border noted with dashed lines, Scale bar = 100 µm). B) Quantitative analysis of (A). Statistical differences determined by Student’s two-sided t-test, where differences are shown for comparison between conditions (*p < 0.05, **p < 0.01; ***p < 0.001). Data shown represent mean ± SD (n=3).

### 2.5. Multiple injuries lead to increased apoptosis relative to single injury

Further, enabled by the photoresponsive synthetic ECM, we introduced multiple injuries to the same location or different locations on the epithelium. Briefly, this was achieved using the same method for creating a single injury and, after the first injury, rotating the photomask ninety degrees to create multiple injuries in the shape of a plus sign (+) upon irradiation. The center point of the plus sign represents multiple injuries at the same location (two overlapping injuries), whereas the segments of the plus sign represent multiple injuries at different locations (four adjacent injuries). Increased cell death (apoptosis or necrosis) was observed proximal and distal to multiple injuries even with a smaller size (50 µm) of injury (Figure 6A, C) as compared to a single injury (Figure 5), more than a two-fold change. Additionally, the number of Ethi-1-positive cells significantly increased relative to Apo-15 positive cells with multiple injuries using 50 and 100 µm slit both proximal and distal to the site of injury (Figure 6B, D) relative to a single injury (Figure 5), suggesting most of the cells entered in necrotic rather than apoptotic phase after multiple injuries of either small or large size. Further, while closing of the injury was observed for a small single injury (Figure 5A), closure was not observed for small multiple injuries, whether overlapping or adjacent multiple injuries (Figure 6A), given the significant level of cell death. Note, since many cells die upon multiple injuries, necrotic cells are washed away during sample processing, so the absolute values of necrotic cells may not fully reflect the number of dead cells upon multiple injuries, particularly with overlapping injuries. Overall, these results suggest that epithelial cells can repair and recover from small single injuries but not from a critical number of injuries or size of injury, where cell death ensues.

**Figure 6.**
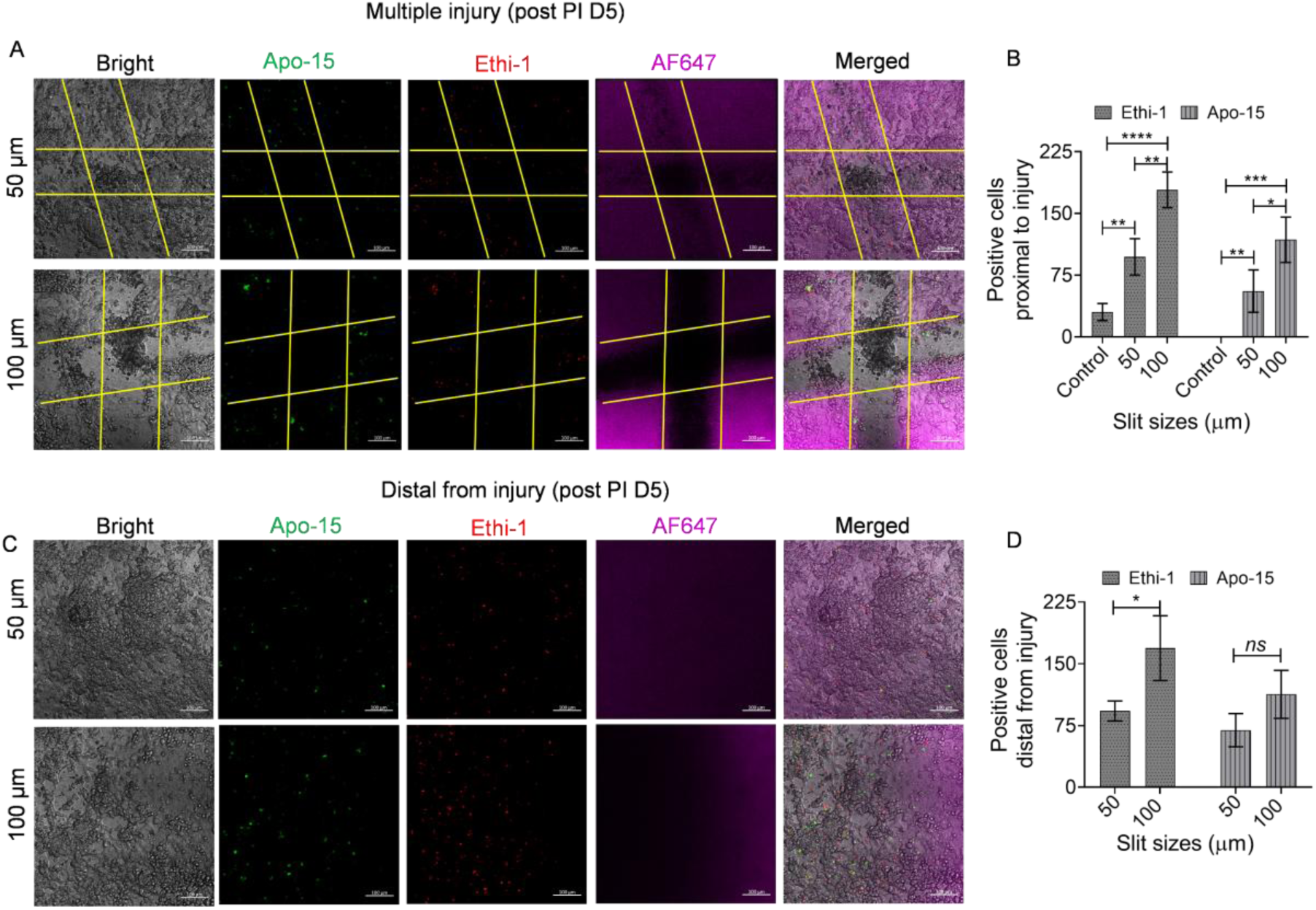
Effects of multiple injuries. Multiple injuries were introduced and led to apoptotic cells both proximal and distal to the injury site 5 days after photo-injury (PI D5). A) Representative images of multiple PI (confocal z-projection, Apo-15 (apoptotic cell death), ethidium homodimer-1 (Ethi-1, necrotic dead cells), hydrogel labeled with AF647, injury boundaries noted with dashed lines, Scale bar = 100 µm). B) Quantitative analysis of (A). C) Representative images showing dead and apoptotic populations distal to the PI area day 5 after multiple PI. D) Quantitative analysis of (C). Significant differences assessed by Student’s two-sided t-test, where differences shown for comparison between PI sizes (*p < 0.05, **p < 0.01; ***p < 0.001). Data shown represent mean ± SD (n=3). Scale bar = 100 µm.

## 3. Discussion

Fibrotic diseases are a significant and growing human health challenge and are attributed to over fifteen percent of deaths worldwide^[3, 13]^. In particular, IPF is one of the leading causes of death in developed countries, and effective medical therapies remain a need^[60–61]^. The origins of IPF amongst other fibrotic diseases are thought to be repeated micro-injuries to the epithelium that are hypothesized to initiate a maladaptive wound healing cascade and lead to fibrotic tissue and disease rather than functional healing^[62]^. However, such diseases are often caught in advanced stages, and exact origins and mechanism are not well understood or captured by existing model systems, impeding the development of effective therapeutics^[33–34, 63]^. Human model systems that enable studies of the initiation of these injury and repair processes for epithelial tissues are needed for studying this complex process and have the potential to provide insights for fibrotic diseases such as IPF and more broadly wound healing, repair, and disease processes in epithelial tissues. In this work, we established a photoresponsive synthetic ECM inspired by the basement membrane for the creation of a model epithelium and used it to probe the mechanisms of epithelial response directly after injuries of different sizes and frequencies.

The photoresponsive synthetic ECM provides spatiotemporal control, preciseness, and reproducibility from the molecular level up to enable control of matrix degradation and injury creation when and where light is applied and with tunability based light intensity and wavelength. Using visible light as a trigger can be advantageous over UV light in that it may pose less threat of biological toxicity^[64–65]^. Photoresponsive material systems have been used to form, soften, pattern and stiffen hydrogel platforms in a number of biological context^[66]^. With respect to the epithelium, synthetic ECMs have been used to model intestinal monolayers, from selective softening or degradation to influence monolayer geometry^[38, 67]^ to deployment of 3D printing for capturing crypt-villus topography^[68]^. In this work, we leverage the tunability of bio-orthoganol azide-cycoloctyne chemistry^[69]^ and the visible light degradation properties of coumarin^[70]^ to fabricate photoresponsive hydrogels with: *i)* sufficiently long gelation working time for sample formation, *ii)* adjustable initial moduli for replication of different stages of healthy to fibrotic tissues, and *iii)* tunable photodegradation based on the wavelength and intensity selected. We then applied these photoresponsive synthetic ECMs for studying epithelial and repair processes using model lung epithelial cells as a prototypical example, as well as given the great need for model systems of relevance for probing the initiation of fibrotic lung diseases.

For inducing and studying responses to injuries in cell culture, scratch assays have traditionally been used^[71–72]^. These methods broadly use mechanical force or similar to create a ‘scratch’ or void on the surface where cells are cultured and then monitor cells filling in this void as a measure of healing. Variations of this assay include: *i)* scratching the surface on which cells are being cultured with a sharp instrument (e.g., pipette tip), removing cells in that region and creating a ‘scratch’^[11, 22, 73–74]^; *ii)* using a laser or microjet to ablate regions of attached cells^[27–28]^; or *iii)* covering a region of the culture surface prior to cell seeding so that a region without cells is revealed upon barrier removal^[75]^. Each of these approaches provides different levels of uniformity and sizes of injury. While these approaches differ in their spatial and temporal control, they all rely on a destructive mechanical injury that lacks molecular specificity. In contrast, the approach we established uses molecular engineering and light to selectively damage the underlying matrix and probe the resulting cell responses.

Here, for probing epithelial cell response to injury, a model epithelial monolayer was formed on a photoresponsive hydrogel with model A549 lung epithelial cells, where including biomimetic peptides PHSRN and AG73 together was found to be synergistic for promoting epithelial layer growth and phenotypic properties. The utilization of the A549 cell line has some limitations for mimicking lung tissue because these immortalized human alveolar cells spread and proliferate rapidly on stiff substrates and collagen-based hydrogels and show mixed phenotypic populations (epithelial and mesenchymal properties)^[11, 34]^. However, we have found that A549 cells expressed adherens junction protein (E-cadherin) associated with epithelial cells with a significant decrease in the number of cells expressing mesenchymal properties (Vimentin) when forming an epithelium on PHSRN+AG73 synthetic ECM. Further, using model airway epithelial cells, Calu-3 cells similarly exhibited an epithelial phenotype during monolayer formation, without a mesenchymal subpopulation, further supporting the synergistic effect of dual biomimetic peptides to develop a healthy model epithelium with relevant phenotypic maintenance. This bioinspired synthetic ECM with tunable and photoresponsive properties provides a new tool for controlling cell cultures, including mechanistic studies of injury, repair, and disease processes toward developing more effective treatment strategies.

The causes of lung injuries and maladaptive wound healing *in vivo* include particulates and pathogens. During these events, epithelial cells can lose cell-cell communication because of high levels of pro-inflammatory factors and resulting degradation of the ECM and lung tissue, triggering cell death mechanisms like apoptosis or necrosis^[11, 20, 62]^. To probe human cell responses to such injuries at their initiation and in real-time, we created injuries of different sizes and frequency by light-triggered degradation the synthetic ECM, and model epithelial cells were observed to alter phenotypes and lose barrier integrity. A low or non-recovery stage for the epithelial cells after critically-sized or multiple injuries was associated with a comprehensive loss of E-cadherin, reduced expression of proliferation and mesenchymal markers, and ultimately the induction of cell death processes. This observation both validates the photo-injury model system relative to known *in vivo* mechanisms and supports the importance of cell-cell interactions in re-epithelization after injury. Cells can recover by maintaining epithelial phenotype in a small wound size; however, cells unable to recover or repair large or multiple wounds are subject to a maladaptive cycle of phenotypic loss, reduced proliferation, and apoptosis or necrosis. Overall, using this PI system that allows the creation of injuries of different sizes and frequency, our findings suggest that epithelial cells not only do not recover from a ‘critical’ injury size but also from repeated injuries, where injury type (size, number) plays a role in healthy versus maladaptive epithelial wound healing processes. This photoresponsive *in vitro* model provides a new tool for understanding mechanisms of epithelial cell injury and repair processes and evaluating approaches for promoting functional rather than maladaptive healing that sets the stage for disease.

## 4. Conclusion

A photoresponsive hydrogel-based synthetic ECM was created that enables controlled induction of epithelial microinjuries, providing a versatile platform for mechanistic studies of epithelium injury and repair processes. This well-defined, bioinspired synthetic matrix supports the formation of a model epithelium with enhanced E-cadherin expression and barrier function, demonstrating its effectiveness as a biomimetic basement-membrane-like environment. Using a photolabile *bis*-coumarin linker, the synthetic matrix undergoes tunable, cytocompatible degradation upon exposure to visible light (e.g., 430 nm), allowing precise modulation of injury size, depth, and frequency. Importantly, injuries beyond a defined critical size elicited epithelial responses characteristic of dysregulated re-epithelialization, including reduced E-cadherin, proliferation, and barrier function, alongside increased apoptosis. This engineered, dynamic ECM provides a powerful tool for recapitulating early events in maladaptive wound-healing cascades and offers new opportunities for dissecting epithelial-ECM interactions during injury initiation. This tunable, responsive platform will prove useful in future mechanistic studies for the discovery of therapeutic targets and the development of more effective therapeutic strategies with the potential for halting not only the progression, but also addressing the initiating events of fibrotic diseases.

## 5. Experimental Section and Methods

### 5.1. PEG-tetra-bicyclononyne (PEG-4-BCN (exo)) synthesis

PEG-4-BCN (exo) was synthesized according to the following method (Figure S24). First, (1R, 8S, 9R, 4Z)-bicyclo [6.1.0]non-4-ene-9-yl-methanol was synthesized using previously published methods^[76]^. Briefly, rhodium (II) dimer tetraacetate (122 mg, 0.276 mM, Combi-Blocks) was added to a flame-dried and nitrogen-purged 100 mL round bottom flask. (Z,Z)-1,5-cyclooctadiene (TCI) was then added and the flask was covered in aluminum foil. Ethyl diazoacetate (Sigma-Aldrich) was added dropwise over one hour. The reaction was then allowed to proceed for an additional hour. The reaction was then concentrated and purified with flash column chromatography (CombiFlash RF200, Teledyne ISCO), protected from light, using a 0-10% ethyl acetate (Fisher Scientific) gradient in hexanes (Fisher Scientific) to produce a clear oil of the mixed isomer product (15.12 g, 96%).

To a nitrogen-purged, flame-dried 500 mL round bottom flask was added potassium tert-butoxide (26 g, KOtBu, Sigma-Aldrich) anhydrous diethyl ether (125 mL, Fisher Scientific) followed by water (1.75 mL) and cyclooctene ester (15 g). This formed a thick tan solution which was mixed vigorously overnight at room temperature. The following day, a 10% sodium hydroxide solution (80 mL, Fisher Scientific) and DI water (20 mL) were added to the reaction vessel, and the aqueous phase was extracted. The organic phase was washed with an additional 40 mL of 10% sodium hydroxide solution, and the aqueous phase was extracted. The aqueous phases were combined and added to a 500 mL round bottom flask and placed in an ice bath to cool for 15 min. Concentrated hydrochloric acid (Fisher Scientific) was then added slowly to adjust pH to <2. A white/yellow solid precipitated and was collected by filtration and was washed with 0.1 N hydrochloric acid (20 mL). The product was then dried in a vacuum overnight (10.68 g, 83%).

To a flame dried, nitrogen purged round bottom flask was added 2 N LiAlH_4_ in THF (52.7 mL, Sigma-Aldrich) and the solution was placed on ice for 15 min. In a separate flame dried, argon purged, 10 mL scintillation vial was added cyclooctene-acid (5 g) and dissolved in anhydrous THF (15 mL, Thermo Scientific). The cyclooctene-acid solution was then added dropwise to the LiAlH_4_ solution over the course of 1 h and left to react at room temperature for 2 additional hours. The reaction was then placed on another ice bath and chilled for 15 min and small drops of water were added until gas evolution was no longer occurring. The whole contents were filtered to remove the alumina and lithium salts. The filtered solid was then washed with excess diethyl ether. The solvent was then collected and washed with water (200 mL). The aqueous phase was washed an additional time with diethyl ether (100 mL). The organic phases were combined and dried over sodium sulfate (Fisher Chemical) followed by filtration and concentration to produce the product as a pale oil (7.52 g, 76.7%).

The synthesis was continued using additional previously published methods^[77]^. The cyclooctene-acid (7.5 g) was added to a flame dried nitrogen purged round bottom flask followed by anhydrous DCM (300 mL, Sigma-Aldrich). This solution was cooled in an ice bath for 15 min. In a separate flame dried and nitrogen purged round bottom flask was added bromine (2.57 mL, Sigma-Aldrich) in DCM (75 mL). This solution was then added slowly to the cyclooctene-acid solution, which turned yellow. Once the yellow color persisted, the solution was left to mix for 5 min followed by quenching with 100 mL of sodium thiosulfate (Fisher Chemical). The organic phase was then extracted twice with 150 mL DCM and then dried over Mg_2_SO_4_ (Research Products International Corp.), filtered, and concentrated to dryness via rotary evaporation producing a yellow oil. The dibromide intermediate was transferred to a purged round bottom flask; 300 mL of anhydrous THF was added; the solution was then cooled in an ice bath for 15 min; KOtBu was then added dropwise; and the reaction mixture was left to stir on the ice bath for 5 min before attaching a reflux condenser and transferring the reaction to an oil bath. The reaction was refluxed at 75 °C for 2.5 h. Then the reaction was taken out of the oil bath and allowed to stir at room temperature for 30 min. After reaching room temperature, the reaction was quenched with the slow addition of saturated ammonium chloride (300 mL, Sigma-Aldrich). The organic phase was then extracted, and the solution was dried over MgSO_4_, filtered, and then concentrated to produce an orange oil which was purified by flash chromatography using a 0-20% ethyl acetate in hexanes gradient. Once dried the product produced a yellow oil (3.6 g, 48.6%).

(1R,8S,9r)-bicyclo[6.1.0]non-4-yn-9-ylmethyl (4-nitrophenyl) carbonate was then synthesized from (1R,8S,9r)-bicyclo[6.1.0]non-4-yn-9-ylmethyl using a previously reported protocol^[78]^. The starting product (3.6 g) was added to a flame-dried and nitrogen-purged round bottom flask, followed by anhydrous DCM (200 mL) and pyridine (5.31 mL, Sigma). Nitrophenyl chloroformate (5.34 g, TCI) was then added and the reaction was allowed to proceed for 30 min before it was quenched with saturated ammonium chloride. The organic phase was then extracted, and the aqueous phase was washed three additional times, each with 100 mL of DCM. These organic phases were then combined, dried over magnesium sulfate, filtered, and concentrated. The product was then purified with flash chromatography using a 0-25% gradient of ethyl acetate in hexanes. Upon drying, the product, (BCN (exo)), was produced as a colorless oil that solidified to a white solid over time (7.1 g, 94%).

Last, 4-arm PEG amine was functionalized with BCN (exo) using previously established methods with slight modifications^[69]^. BCN (exo) and 4-arm PEG(10k)-NH_2_ (JenKem Technology) were added to a flame-dried and nitrogen-purged 20 mL vial, followed by anhydrous DMF (5 mL, Thermo Scientific Chemicals) and N,N-diisopropylethylamine (412 mg, DIPEA, Sigma-Aldrich).

Once DIPEA was added the solution turned yellow. The reaction was allowed to proceed for 24 h in the dark. This solution was then precipitated in cold diethyl ether (40 mL) and then cooled in a freezer for 15 min, centrifuged down, and the solvent was decanted. The PEG pellet was washed two additional times using fresh, cold diethyl ether. The pellet was then dried with nitrogen, dissolved in minimal DI water (10 mL), filtered using a 0.45 µm filter (PES, 25mm, Avantor), and dialyzed for 36-48 h with 3.5 kDa MWCO snakeskin dialysis tubing (Thermo Scientific). The product was then collected, frozen, and lyophilized to produce the product (2.02 g, 85% functionalized, Figure S25).

### 5.2. PEG-bis-coumarin-azide (PEG-2-CmPN_3_) synthesis

PEG-*bis*-coumarin-azide was synthesized using a previously published multi-step synthetic route, where the CmPN_3_ synthetic procedure provides a water-stable product and scalable synthesis^[79]^ (Figure S26). Briefly, 7-diethylamino-4-methylcoumarin (Sigma-Aldrich) was reacted with dimethylformamide-dimethyl acetal (Chem-Impex International) in DMF under reflux. The resulting dimethylamino coumarin intermediate, a yellow solid, was then triturated with acetone (Fisher Scientific), filtered, and dried (10.62 g, 76%). Sodium periodate (Acros Organics) was then reacted with this intermediate in 1:1 THF:DI water at room temperature for 2 h. 7-Diethylamino-4-formyl-coumarin, a red oil, was extracted, filtered, and concentrated (9.02 g, 97%).

This intermediate was added to propargyl bromide (Sigma-Aldrich), zinc (Aldrich), and THF at 0 °C with trimethyl chlorosilane (Sigma-Aldrich) as a catalyst, resulting in the coumarinyl alcohol derivative (6.88 g, 71%). Ethyl 4-azidobutanoate was synthesized from ethyl-4-bromobutyrate (TCI) and sodium azide (TCI) in DMSO (Fisher Scientific) (26.8 g, 94%). The coumarinyl alcohol, ethyl-4-azidobutyrate, and CuSO_4_ (Sigma-Aldrich) were added to 9:1 methanol:DI water (Fisher Chemical) at room temperature followed by (+)-sodium-L-ascorbate (Sigma) and overnight mixing. The solution was then concentrated, extracted, washed, and purified by column chromatography producing an oil (8.64 g, 91%).

Ten grams of the product was then dissolved in dry DCM and pyridine with 4-dimethylaminopyridine (DMAP, Alfa Aesar) as a catalysis, cooled to 0 °C, and reacted with 4-nitrophenyl chloroformate (TCI) for 4 h. The organic phase was extracted, combined, dried, filtered, concentrated, and then purified by flash chromatography resulting in an orange oil (11.6 g, quant.). Azido-PEG_4_-amine (BroadPharm) was then substituted for the nitrophenyl group by dissolving the coumarin product in anhydrous DCM and slowly adding azido-PEG_4_-amine, anhydrous pyridine, and DMAP at 0 °C to room temperature overnight. The azido-PEG_4_-coumarin intermediate was then extracted, dried, filtered, concentrated, purified, and dried, producing an orange oil (13.35 g, quant.). This was then dissolved in 2:1 THF:water and reacted with LiOH overnight. The product was then extracted, dried, filtered, and concentrated before being purified via flash chromatography, which produced an orange/ yellow oil (2.33 g, 67%). Finally, the photolabile molecule was coupled onto PEG (3.4kDa) *bis*-amine (JenKem Technology) using HATU (Fisher Chemical) and DIPEA in anhydrous DMF reacting overnight. This was then washed with ether, dried, and then dialyzed for 48 h to produce the functionalized PEG product (PEG-2-CmPN_3_) (1.83 g, 81% yield, 90% functionalized, Figure S27).

### 5.3. Peptide synthesis

Solid Phase Peptide synthesis (SPPS) was used to synthesize the peptides used in these studies employing methods previously described^[80]^. Briefly, PHSRN (K(N_3_)-GWG-PHSRNG_10_RGDSPG) and AG73 (K(N_3_)-GWG-RKRLQVQLSIRT) were synthesized. Each peptide was designed with the following functionality: *i)* lysine-azide (K(N_3_)) to enable SPAAC for incorporation of the peptide into the hydrogel network, *ii)* GWG to enable the use of UV-Vis to confirm concentration of solubilized peptide, and *iii)* biologically active sequence. Each of these segments is denoted by a dash within the overall sequence. In brief, Liberty Blue Automated Microwave Peptide Synthesizer (CEM, Matthews, NC) was used to synthesize peptides on rink amide 4-methylbenzhydrylamine (MBHA) resin (Novabiochem). Fmoc chemistry and amino acids (Chempep) were used with triple deprotection and triple coupling steps. Coupling for all amino acids was done with microwave assisted heating at 75 °C for 8 min per coupling, except arginine (R) where coupling was done at 25 °C for 25 min followed by 2 min of microwave assistance heating at 75 °C. Following synthesis, the peptide was then cleaved from resin using TFA (95 % v/v) (Thermo Fisher), TIPS (2.5% v/v) (Chem Impex International Inc.), DI water (2.5% v/v), DTT (2.5% w/v) (Chem Impex International Inc.), and phenol (2.5% w/v) (Sigma-Aldrich) stirring at 390 rpm for 4 h. After 4 h, the cleavage solution was collected, and the peptide precipitated from the cleavage solution in cold ethyl ether (Fisher Scientific). The peptide was washed five times with fresh cold ethyl ether before drying overnight. It was then dissolved in 95:5 DI H_2_O:acetonitrile (Thermo Fisher), and high-performance liquid chromatography (HPLC; XBridge BEH C18 OBD 5 μm column; Waters, Milford, MA) was then used to purify the peptide. A linear water-ACN gradient (water:ACN 95:5 to 65:35; 1.17% change in water per min) was used. Peptide identity was then confirmed by molar mass via UPLC-MS (Figure S28). Purified peptides were then lyophilized and stored at -80 °C until use for hydrogel formation.

### 5.4. Cell sources and maintenance

A549 and Calu-3 cell lines (ATCC) were expanded and maintained in Dulbecco’s Modified Eagle’s Medium/Hams F-12 50/50 Mix with L-glutamine and 15 mM HEPES (DMEM-F12 50/50, Corning Cellgro) supplemented with 10% v/v fetal bovine serum (FBS, Invitrogen), 1% penicillin-streptomycin (PS), and 0.2% v/v fungizone. For real-time monitoring of cell phenotype, a reporter A549 cell line was used, where DsRed-Express2 is constitutively expressed and ZsGreen is conditionally expressed when αSMA is expressed as previously described^[52]^. Media was replaced every 2-3 days, and cells were trypsinized (trypsin/EDTA, Fisher Scientific) at ∼80-90% confluency during routine cell expansion and subculture. A549 and Calu-3 cells at passages 4-10 were used in experiments.

### 5.5. Model lung epithelial cell monolayer formation on photoresponsive hydrogel

Photoresponsive hydrogels (100 µL) were prepared with PEG-4-BCN, PEG-2-CmPN_3_, and selected integrin-binding peptides (PHSRN, AG73, or PHSRN+AG73) in 12-well transwell insert (Corning #3460) and sterilely incubated overnight at 37 °C for full gelation. Briefly, stock solutions of PEG-4-BCN, PEG-2-CmPN_3_, and PHSRN were prepared in sterile PBS and of AG73 in sterile DI water, where peptide concentrations were verified with UV-Vis spectroscopy (Nanodrop 2000). A precursor solution was then prepared, cast onto the transwell surface in a dropwise fashion, and allowed to polymerize overnight (5 or 9 wt% PEG with 2 mM total integrin-binding peptides (1:1 azide:BCN) with a total volume ∼ 400 µL for preparation of ∼ 4 samples at a time with 100 µL per sample). All the hydrogels then were hydrated with sterile DI water overnight, washed with sterile 500 µL of PBS for 2-6 h, and washed with 500 µL of DMEM-F12 before seeding with cells, producing hydrogels with thickness of approximately 400 µm within the transwell inserts as characterized with confocal (see sections below). A549 cells at ∼ 80-90% confluency were trypsinized from the tissue culture flask, pelleted by centrifugation, resuspended in media, and seeded onto the hydrogels in 12-well transwell inserts at 200,000 cells per insert. Cells were cultured submerged in DMEM-F12 growth media over 10 days. DMEM-F12 media was changed every 2-3 days over time in culture.

### 5.6. Cell viability and coverage for monolayer formation using calcein area measurement

Viability of A549 and Calu-3 cells on the hydrogels was performed using a LIVE/DEAD membrane integrity assay (Thermo Fisher Scientific), where Calcein stain was used to label live cells and percentage of Calcein area used to assess monolayer formation. Briefly, media was removed from the insert (n=3) and bottom chamber of 12-well plates. Cells were incubated with 2 μM Calcein acetoxymethyl and 4 μM ethidium homodimer-1 per milliliter of solution in PBS for 18 min at 37 °C (5% CO_2_) after washing twice with 300 µL of PBS for 5 min. Before imaging, cells were washed two times (each 5 min) in 300 μL of PBS and imaged using a confocal microscope (Zeiss LSM 800, 10X objective and image frame size of 1024 × 1024, z-projection of z-stack <10 µm with ∼2 µm slice interval, three images per hydrogel sample). The live (green) and dead (red) cells were counted using Volocity software, and the percentage of the viable cells was calculated using the number of green positive cells/total number of cells × 100%. Further, monolayer formation was assessed based on Calcein (green) positive area using Fiji ImageJ software.

### 5.7. Probing cell response to photoinjury

A549 reporter cells were cultured on the hydrogel for monolayer formation and then monitoring of responses *in situ* and at specific time points of interest. An injury was created on the epithelial monolayer at day 5 in culture using a photolithography-based method, and the time and injury size was controlled utilizing photolithography within 3D printed apparatus designed for this experiment (Figure S3). Briefly, the 3D printed parts were fabricated with a M1 Carbon printer using UMA90 resin. To assemble the apparatus for sample irradiation, a transwell culture insert was placed in a sterile 3D printed black bottom chamber. Then, a photomask with negative feature ‘slit’ of defined width (50, 100, or 200 μm) was placed on the media within the insert, where the photomask was cut from a sheet of photomasks (Advance Reproductions Corporation) with a 10-mm disposable biopsy punch (Robbins Instruments). Excess media was then removed from the insert using a pipetteman, lowering the photomask toward the culture surface and leaving ∼100 µL of media to prevent damage to the epithelial layer. A 3D printed black cone was then placed on the slit photomask, avoiding pressing the photomask on to the monolayer while holding it in place. Next, a 3D printed small black ring was placed inside the 3D printed black cone to further protect any areas outside of the photomask footprint from irradiation. The photo-injury was performed with 6.75 mW*cm^-2^ (430 nm typically for 4 min, where 2-6 min were probed; 455 nm also probed; Mightex WheeLED Wavelength-Switchable LED Sources (VIS-NIR) with BLS-1000-2 driver) while maintaining a sterile condition. To allow imaging of the injury, hydrogels were conjugated during their formation with Alexa Fluor™ 647 Azide (Thermo Fisher Scientific), integrating 3 µM into the precursor solution. Injury and the cell response at the injury area was assessed using confocal microscopy (Zeiss LSM 800, 10X objective and image frame size of 1024 × 1024, z-projection of z-stack <10 µm with ∼2 µm slice interval). Samples were washed with sterile PBS, then fresh media was added, and samples were transferred to the incubator. The cell response was monitored *in situ* after photo-injury every 24 h over 5 days using confocal microscopy (Zeiss LSM 800 confocal microscope equipped with an incubation chamber (37 °C, 5% CO2, with humidity control); for supplementary videos, brightfield images were captured every 30 minutes over the course of 12 hours. Samples were fixed for further analysis at the end of day 10 (total culture time), as described in the immunostaining section.

### 5.8. Cellular barrier integrity measurement of epithelial monolayer

Trans-Epithelial Electrical Resistance (TEER) is a known technique to measure the integrity of tight junction dynamics *in vitro* cell culture models of epithelial monolayers. Briefly, the A549 monolayer was cultured on the hydrogel (PHSRN+AG73) in the insert over five days as described above. The media was removed from both apical and basolateral compartments, and 500 µL sterile PBS was added. For TEER measurements, we used an Epithelial Voltohmmeter (EVOM2) system with a ‘chopstick’ electrode pair, where one electrode was placed in the apical and the other in basolateral compartment. The measurement range for EVOM2 system is 1-10,000 Ω. All measurements were conducted in triplicate (n=3) at specificied time points to assess barrier integrity. The calculation for resistance involved measuring the blank resistance (R_blank_) of the semipermeable membrane only with PBS (without cells) and measuring the total resistance across the cell layer (R_total_). The cell-specific resistance (R_cell_) in units of Ω can be obtained using the following equation:

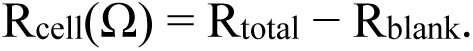

Resistance is inversely proportional to the effective surface area of the semipermeable membrane (M_area_), which is reported in cm^2^ units:

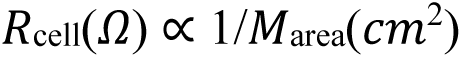

Accordingly, TEER values are reported (TEER_reported_) in units of Ω.cm^2^ and calculated according to the following equation:

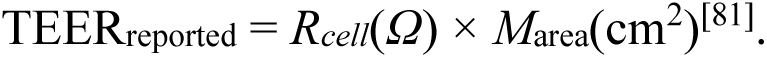

### 5.9. Immunofluorescence staining

Media was removed from the insert, and the sample was gently washed twice with PBS at 37 °C. Cells were fixed with 4% PFA for 30 min at room temperature and washed twice with PBS for 5 min. Cells were permeabilized (1% (w/v) BSA + 0.1% (v/v) Triton X-100 in PBS) and blocked (3% BSA (w/v) + 0.05% (v/v) Triton X-100 in PBS). After blocking, cells were incubated with primary antibody (1:500 dilution of mouse anti-human E-cadherin, Abcam, or mouse anti-human Alexa Fluor 488 conjugated Vimentin, Abcam) in blocking solution overnight at 4 °C. The next day, samples were washed thrice with permeabilization solution (each 10 min) and incubated overnight with secondary antibody Alexa Fluor 488 goat anti-mouse (1:200 dilution; Thermo Fisher Scientific) and F-actin red (40 μL per milliliter; Sigma-Aldrich) in blocking solution at 4 °C. Next day, samples were rinsed thrice (each 15 min) with PBS at room temperature and imaged (n = 3) on a confocal microscope (Zeiss LSM 800, 20X objective and image frame size of 1024 × 1024, z-projection of z-stack <10 µm with ∼2 µm slice interval). The quantification analysis of E-cadherin and Vimentin-positive cells was performed using the Fiji ImageJ software.

For Ki-67 and αSMA staining, cells were incubated with primary antibody (1:300 dilution of rabbit anti-human Ki-67 and 1:500 dilution of mouse anti-human αSMA, Abcam) in blocking solution overnight at 4 °C, followed by the steps as described above. The secondary antibody, Alexa Fluor 647 goat anti-rabbit with 1:300 dilution, was used for Ki-67 staining (Thermo Fisher), and Alexa Fluor 488 goat anti-mouse with 1:500 dilution was used for αSMA staining (Thermo Fisher).

### 5.10. Apoptotic cells staining

Cells were stained with Apotracker-green (Apo-15, excitation/emission maxima 500/520 nm, Biolegend) and ethidium homodimer-1 (Excitation/emission maxima 528/617 nm, Thermo Fisher) for detecting apoptotic and necrotic (dead) cells, respectively. Briefly, reconstituted Apo-15 solution was used at a 1:10 dilution in FACS buffer (2% FBS in PBS). Then, 50 μL mL^-1^ of the diluted apo-15 reagent and 2 μL mL^-1^ ethidium homodimer-1 reagent were mixed in FACS buffer. 300 μL of staining solution was added to each sample at selected time points and incubated at room temperature for 18-20 minutes. Samples were washed at least 2 times with buffer and imaged with confocal microscopy as described above.

## Statistical Analysis

All conditions were processed with three or more replicates, as noted within individual figure captions. All the experiments were repeated more than two times (biological replication) to reproduce results. All data were analyzed using GraphPad Prism 8. Student’s t-tests (unpaired, two-tailed) were performed to compare the means of two normally distributed groups. *p < 0.05, **p < 0.01, ***p < 0.001, and ****p < 0.0001.

## Acknowledgments

This work was supported by National Institutes of Health (NIH) Director’s New Innovator Award with grant number DP2HL152424 (Kloxin), as well as partial support by the National Science Foundation (NSF) through the University of Delaware Materials Research Science and Engineering Center (DMR-2011824). Instrumentation and facilities were supported in part by an Institutional Development Award from the National Institute of General Medical Sciences (NIGMS) (P30GM110758) from the NIH and the NSF through the University of Delaware Materials Research Science and Engineering Center (DMR-2011824). The content is solely the responsibility of the authors and does not necessarily represent the official views of the NIH or NSF. K.B. acknowledges partial support by a Collins Fellowship, Q.Z. acknowledges partial support by the Undergraduate Research Program and Dare to BE FIRST REU (NSF-1460757), and S.S. acknowledges support from the UD Doctoral Fellowship from the Graduate College. The authors thank Prof. Kelvin Lee and his lab for access to TEER and helpful discussions of its use, as well as William Trout for support of its maintenance. L.P., B.S., K.B., S.S., S.C., C.F., and A.M.K. acknowledge the submission of a patent application related to the work (application no. PCT/US2024/054209; filing date 1 Nov. 2024).

## Author contributions

**Lina Pradhan:** conceptualization, data curation, formal analysis, investigation, methodology, validation, visualization, writing – original draft preparation, writing – review and editing. **Bryan**

**P. Sutherland:** conceptualization, data curation, formal analysis, investigation, methodology, visualization, writing – original draft preparation. **Samantha L. Swedzinski:** formal analysis, investigation, methodology, validation, visualization, writing – original draft preparation, writing – review and editing. **Kartik Bomb:** conceptualization, data curation, formal analysis, investigation, methodology. **Qi Zhang:** formal analysis, investigation, methodology, validation.

**Samantha E. Cassel:** conceptualization, investigation, methodology. **Catherine A. Fromen:** conceptualization, resources, supervision. **April M. Kloxin:** conceptualization, funding acquisition, project administration, resources, supervision, writing – review and editing.

## Data availability

The data that support the findings of this study are available from the corresponding author upon request.

## Table of Contents (TOC)

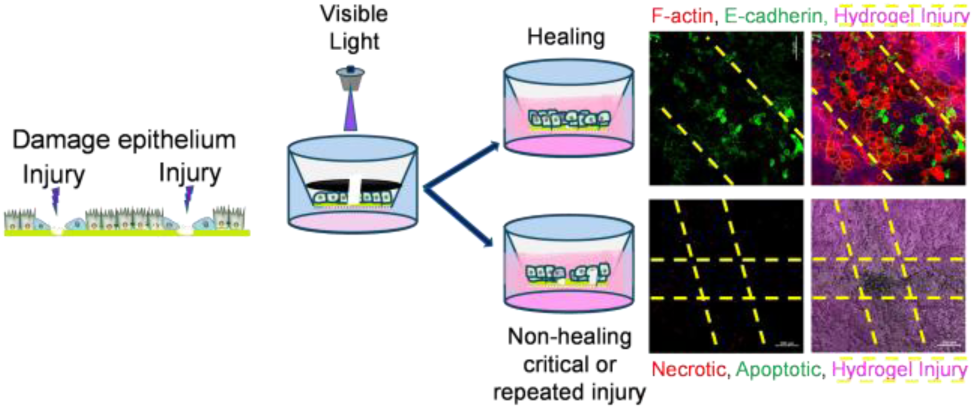

We established a photoresponsive synthetic ECM to model lung epithelial microinjuries. Small injuries allowed functional repair. However, injuries surpassing a critical size or repeated injuries led to dysregulated healing. This failure involved decreased barrier function, loss of epithelial function, and increased cell death, indicating a mechanism for the initiation of maladaptive fibrotic processes.

