## Supplementary figures for "Probing Critical Injury Thresholds for Maladaptive Epithelial Injury and Repair Processes with Photoresponsive Bioinspired Synthetic Basement Membrane"

S. L. Swedzinski, A. M. Kloxin

Department of Materials Science and Engineering, University of Delaware, 127 The Green,

Newark, DE 19716, USA

S.L Swedzinski and K. Bomb Equal contribution

**Data availability**: The data that support the findings of this study are available from the corresponding author upon request.

**Supplementary figures**

**Figure S1.** Rheometric characterization of hydrogel formation and degradation.

**Figure S2.** Photodegradation can be performed with different wavelengths of light *in situ*.

**Figure S3.** Photolithography setup for irradiation of samples in cell culture.

**Figure S4.** Design a 3D printed parts for irradiation of epithelial layer in transwell insert.

**Figure S5.** Imaging approach for characterization of injury to synthetic ECM upon irradiation. **Figure S6.** Monitoring of attachment and monolayer formation of A549 cells in brightfield. **Figure S7**. Assessment of model epithelial monolayer formation with A549.

**Figure S8.** Assessment of model epithelial monolayer formation with Calu-3.

**Figure S9.** Monitoring of attachment and monolayer formation of Calu-3 cells in brightfield.

**Figure S10.** Immunofluorescence staining to assess A549 epithelial monolayer formation.

**Figure S11.** Immunofluorescence staining to assess A549 expression of mesenchymal marker. **Figure S12.** Immunofluorescence staining to assess Calu-3 epithelial monolayer formation.

**Figure S13.** Immunofluorescence staining to assess Calu-3 expression of mesenchymal marker.

**Figure S14.** Epithelium formation *in situ* using reporter A549 cells.

**Figure S15.** Representative images of recovery after photo-injury (PI) using A549 cells.

**Figure S16.** Response to injury (50 µm) using reporter A549 cells.

**Figure S17.** Response to injury (100 µm) using reporter A549 cells with comparison to 50 µm.

**Figure S18**. Irradiation of cells on a non-degradable synthetic ECM.

**Figure S19**. Exposure of cells to photodegradation products.

**Figure S20.** Response to mechanical scratch on synthetic ECM using reporter A549 cells.

**Figure S21**. Immunostaining for mesenchymal marker proximal to photo-injuries of different widths.

**Figure S22**. Immunostaining for epithelial marker distal to photo-injuries of different widths.

**Figure S23**. Immunostaining for phenotypic and proliferation markers proximal and distal to photo-injuries of different injury widths.

**Figure S24.** Synthetic scheme for PEG-4-BCN (exo).

**Figure S25.** NMR of PEG-4-BCN (exo).

**Figure S26.** Synthetic scheme for PEG-2-CmPN_3_.

**Figure S27.** NMR of PEG-2-CmPN_3_.

**Figure S28.** Representative mass spectrometry of bioactive peptides.

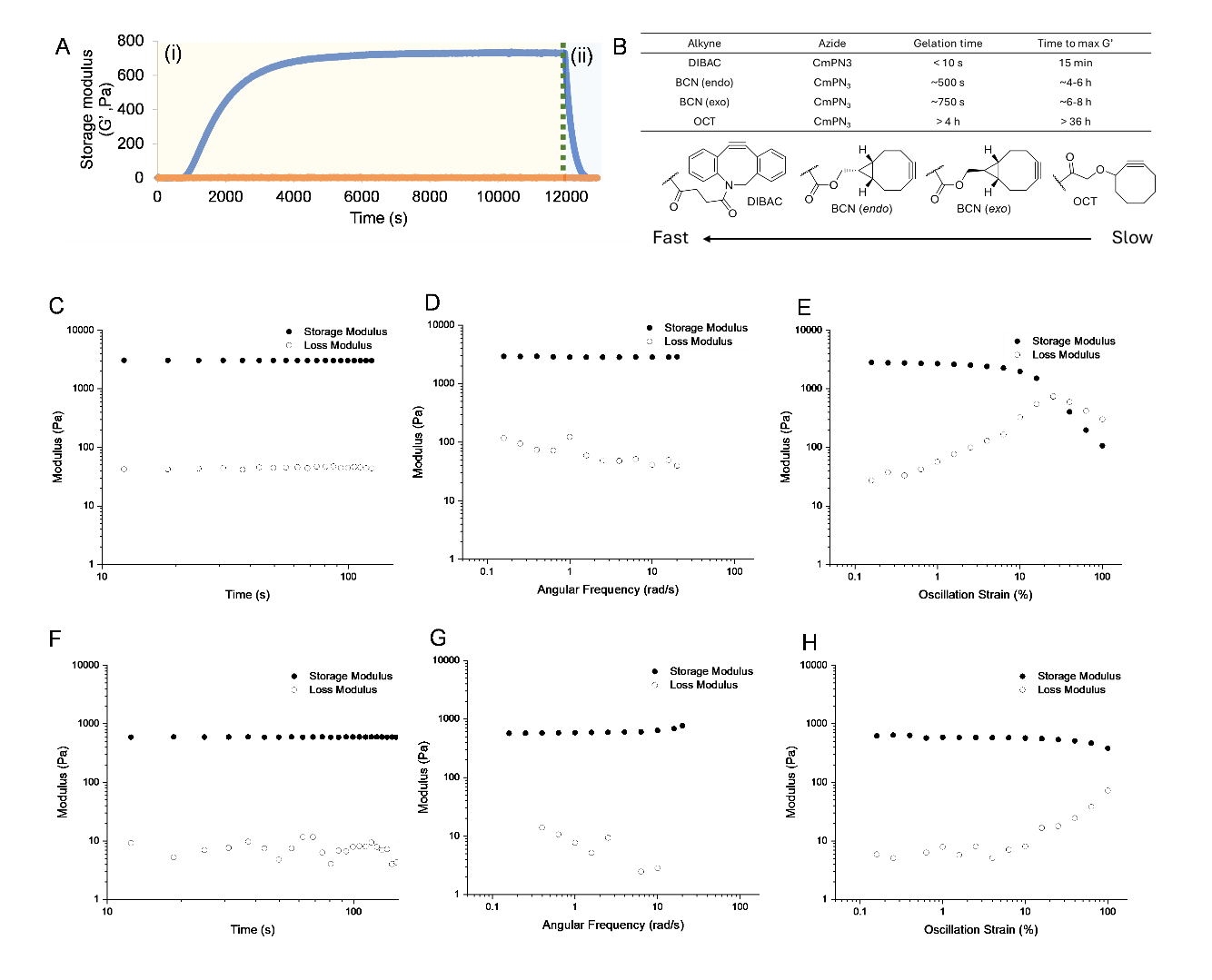

**Figure S1.** Rheometric characterization of hydrogel formation and degradation. A) Hydrogel precursor solution was polymerized *in situ* on a rheometer to assess formation (yellow region) and then monitor degradation (blue region) upon the application of light (dashed green line). B) Different PEG-*tetra*-cyclooctynes (e.g., dibenzocyclooctyne (DIBAC), bicyclononyne (BCN), Cyclooctyne (OCT)) and PEG-*bis*-azide (e.g., coumarin-PEG_2_-azide (PEG-2-CmPN_3_)) were evaluated in pilot studies to identify a pair of reactive handles that provided sufficient working time prior to gelation for preparation of multiple samples with the mixed precursor solution, as assessed by gelation time estimated by (G’>G”), and achieve complete gelation in less than 8 hours, as assessed by time to maximum G’, with consideration of workflow cycling times. Here, BCN (exo) and CmPN_3_ were selected accordingly, enabling the preparation and formation of 3 to 4 samples per batch of precursor solution. Representative rheological measurements of PEG based hydrogels C) Time, D) frequency and E) strain sweeps representative of rheology measurements of 9 wt% PEG hydrogels and F) time, G) frequency and H) strain sweeps representative of rheology measurements of 5 wt% PEG hydrogels.

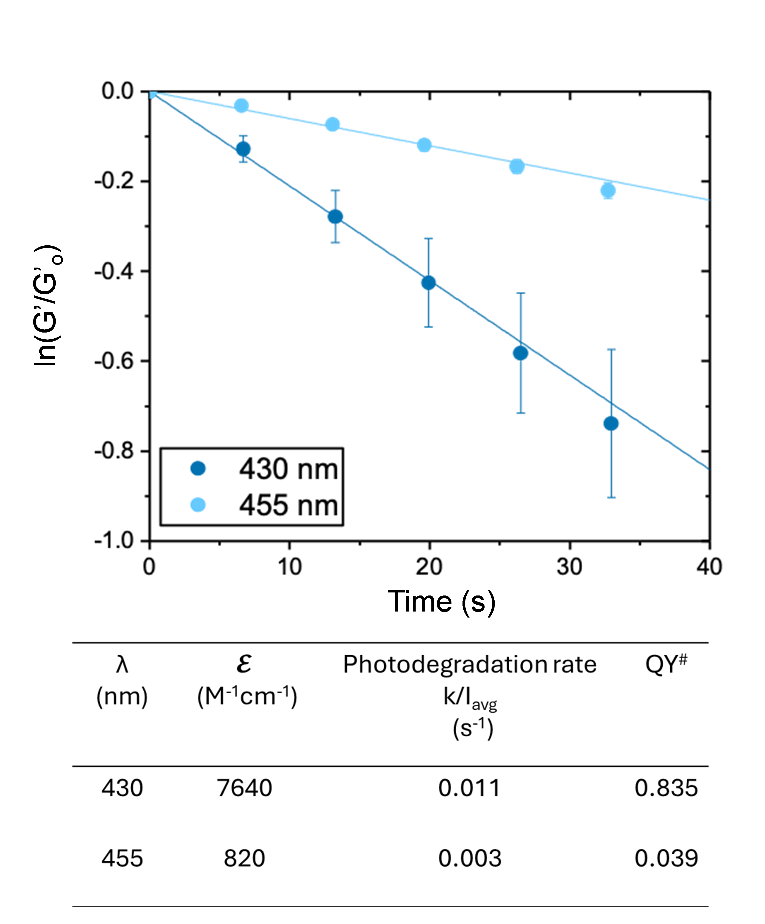

**Figure S2.** Photodegradation can be performed with different wavelengths of light *in situ*. Here, a 50 µm gap size was used to enable light penetration through the sample at different wavelengths, with greater than 50% transmittance based on the Beer-Lambert Law and molar absorptivity at each wavelength (𝓔, Table). The rate of photodegradation is dependent on both the absorbance of light (molar absorptivity) and quantum yield for the photolabile molecule at the selected light wavelength and scales with the light intensity. Here, we aimed to irradiate a 400-µm thick sample within a transwell insert during cell culture, and 430 nm light was selected for creating an ‘injury’ to the synthetic ECM owing to *i)* strong light absorbance for degradation by surface erosion for that thickness and *ii)* rapid photodegradation rate of linkers within the underlying polymer network with a low cytocompatible dose of visible light. Note, based on the Beer-Lambert Law, a shift to a less absorbed wavelength (e.g., 455 nm) would be expected to result in transmission of light through a 400-µm thick sample for degradation of the bulk. Based on the molar absorptivity and concentration of the photolabile group, the percentage of light transmitted through a 400-µm thick sample at 430 nm is ~ 0.4% and at 455 nm is ~ 55%.

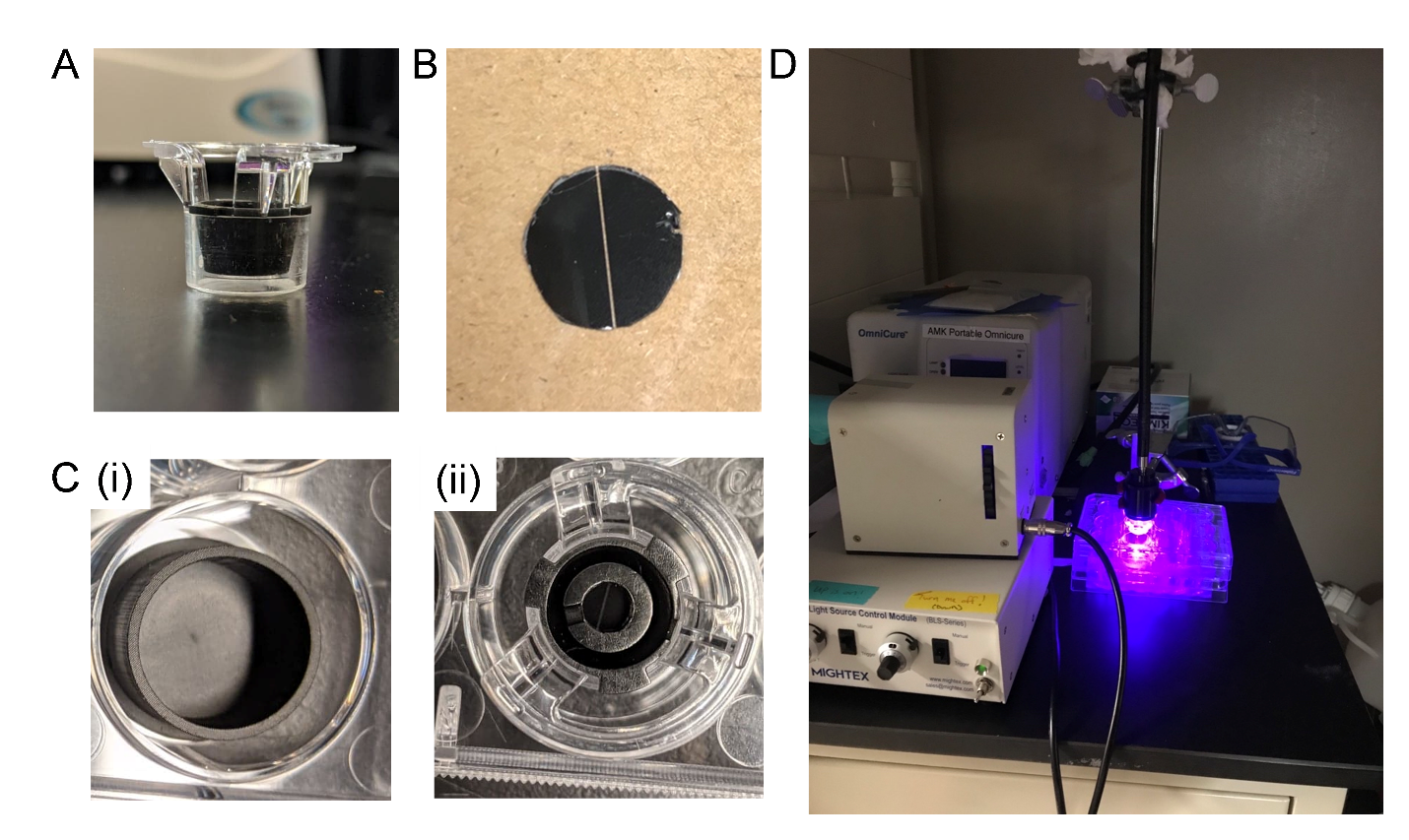

**Figure S3.** Photolithography setup for irradiation of samples in cell culture. A) A 3D printed part was designed to hold a photomask within a transwell culture insert proximate yet sufficiently above the transwell membrane surface to prevent disturbing the underlying attached cell layer. B) Example of a printed photomask with negative feature of a line (‘slit’), where width of the feature was varied in these studies to create different ‘injury’ sizes upon synthetic matrix photodegradation. C) Top-down view of *i)* 3D printed transwell black holder, *ii)* 12-well transwell insert with 3D printed parts, including ‘slit’, a small black 3D cone, and 3D small black ring. Volume of media on top of the transwell insert is controlled (~50-100 µL), and 3D printed parts then hold photomask in place and prevent light for being transmitted to neighboring samples during irradiation. D) Example irradiation setup: here, LED system (left) with connected light guide and collimating lens held at appropriate height with ring stand and clamp for irradiating a well of interest (right) with desired intensity; plate sterility is maintained outside of biosafety cabinet by irradiating through plate lid, which was secured to plate with parafilm. Here, irradiation intensity of 6.75 mW cm^-2^ was used to compensate for absorbance of light be the plate lid and achieving an intensity of ~ 5 mW cm^-2^ at the sample surface.

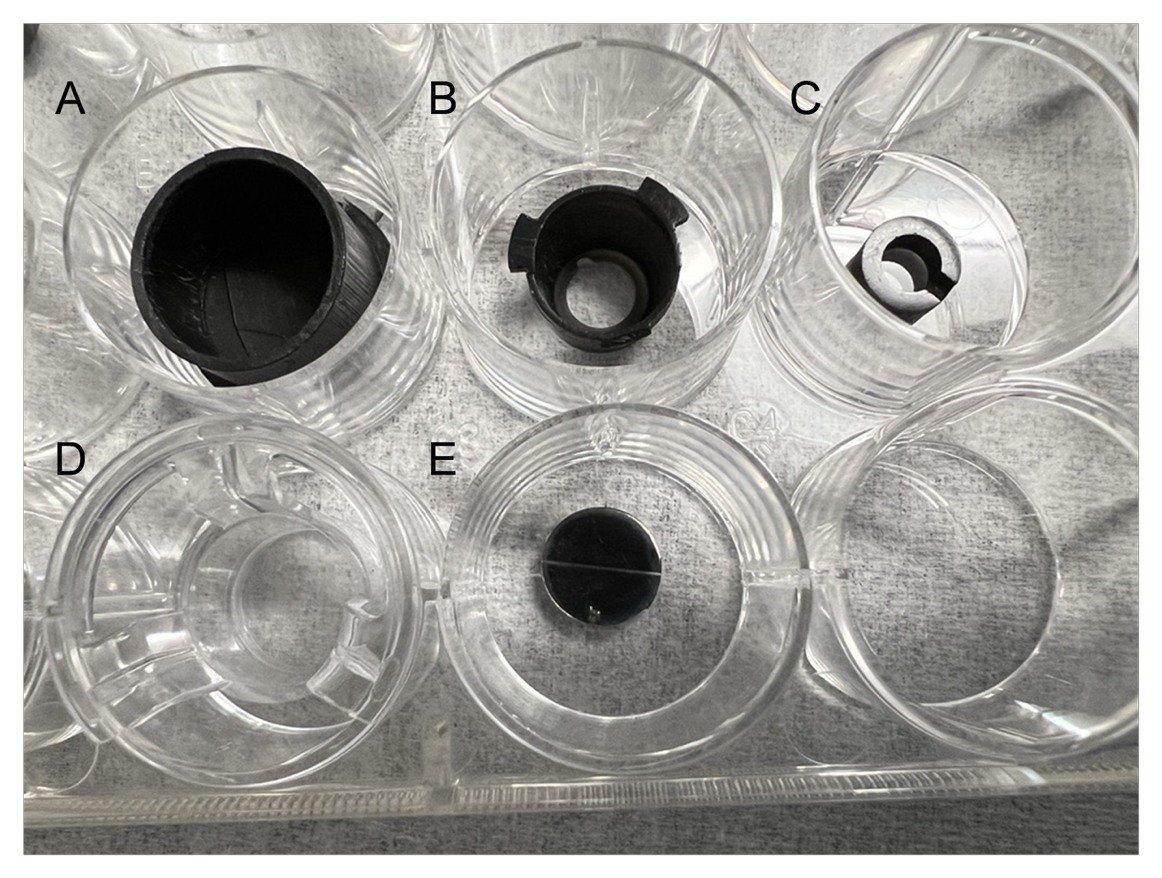

**Figure S4.** Design a 3D printed parts for irradiation of epithelial layer in transwell insert. A) Black bottom chamber transwell holder for protecting insert from bottom light. B) Black cone for blocking irradiation of undesired areas from a top view. C) Black ring for protecting against the extra irradiation from the edge of the 3D cone. D) 12-well insert. E) Photomask ‘slit’. Assembly of the parts for irradiation of samples is shown in Video S1.

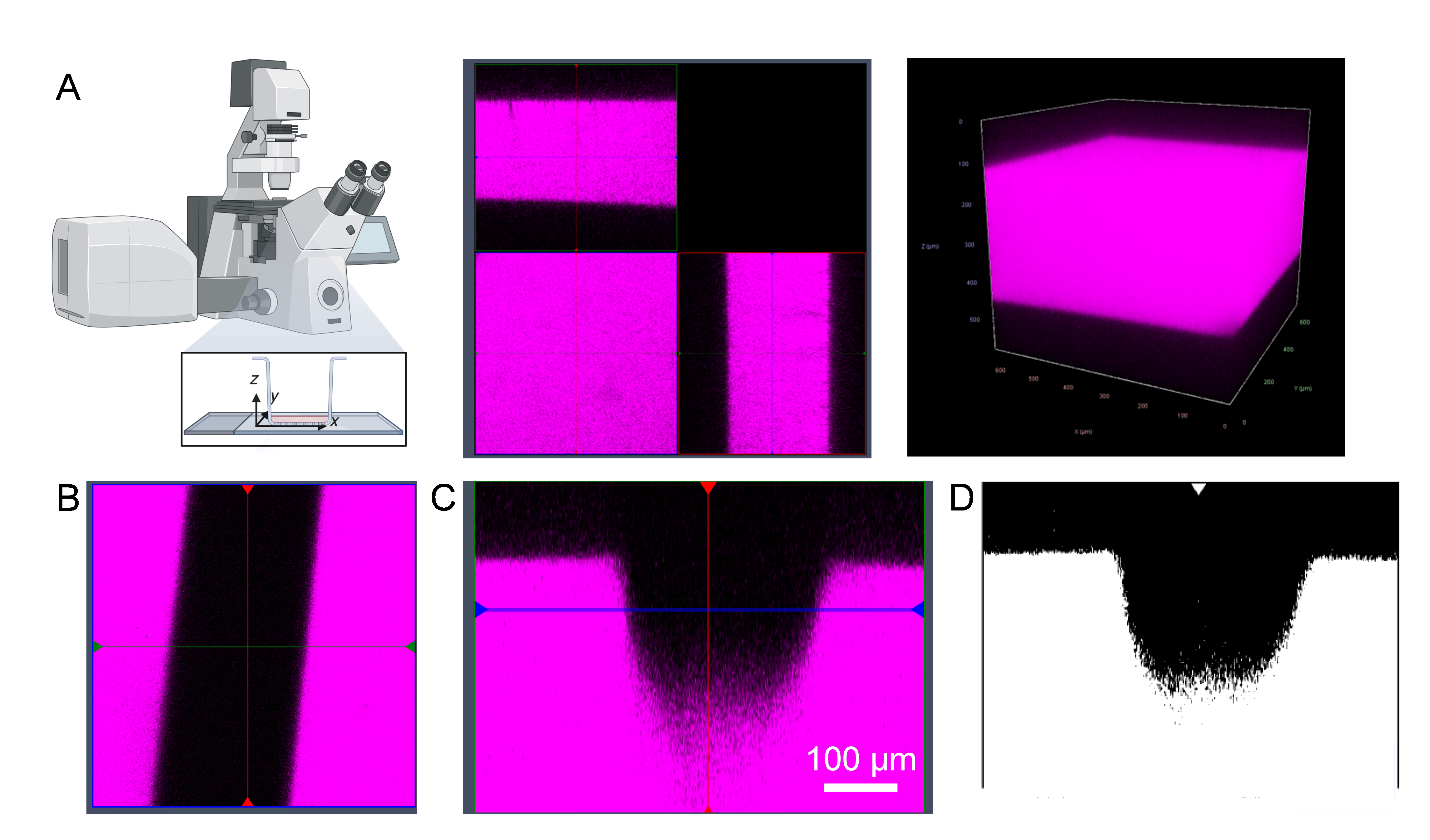

**Figure S5.** Imaging approach for characterization of injury to synthetic ECM upon irradiation. A) Representative images of synthetic ECM layer (here, labeled with AF647; top-down view (middle); 3D view (right)) created on transwell insert and imaged with confocal microscopy (left). B) Representative top-down view of confocal z-stack (LSM 800, 10X objective and image frame size of 1024 × 1024) of synthetic ECM after ‘injury’ (absence of fluorescence owing to synthetic ECM degradation). C) Side view of the same confocal z-stack shown in (B): red line notes the same x-y position between (B) and (C) running parallel with the phototriggered ‘injury’; the blue line notes the z-position of (B) within the z-stack; and the green box around (C) itself indicates the cross-section of (B) that is being viewed, that is perpendicular to the phototriggered ‘injury’. D) Example of cross-section image converted to a binary image (here, fluorescence converted to white, no fluorescence converted to black) from (C) for quantitative analysis of ‘injury’ width and depth with Image J.

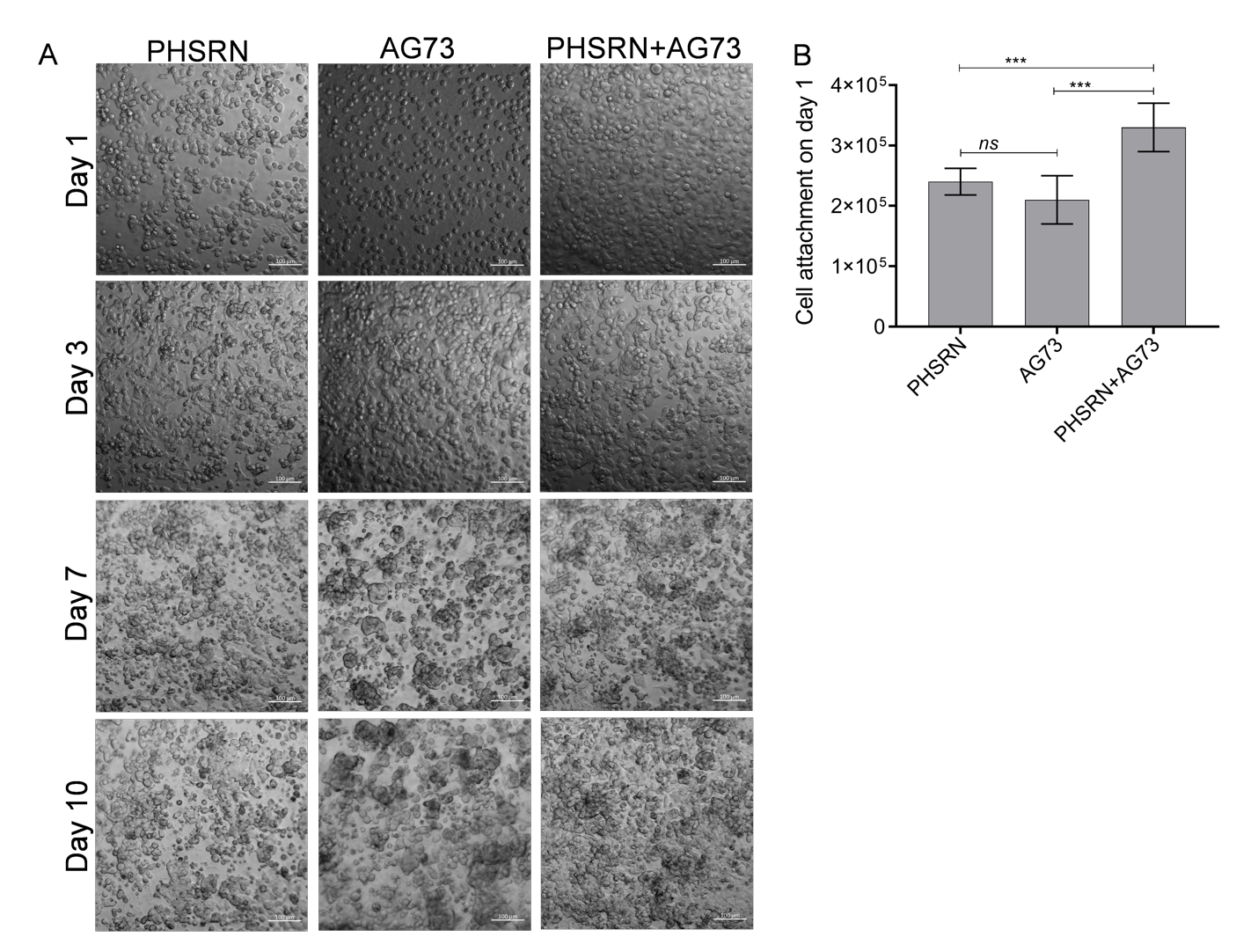

**Figure S6.** Monitoring of attachment and monolayer formation of A549 cells in brightfield. A) Representative images of A549 cells cultured over time from single cell attachment to confluent monolayer formation on different soft synthetic ECM compositions (PHSRN, AG73, or PHSRN+AG73) (confocal LSM 800, 10X objective and image frame size of 1024 × 1024). B) Quantitative analysis of initial A549 cell attachment at day 1. Significant differences assessed by Student’s two-sided t-test (*p < 0.05, **p < 0.01; ***p < 0.001). Scale bar = 100 µm. Data shown represent mean ± SD (n=3).

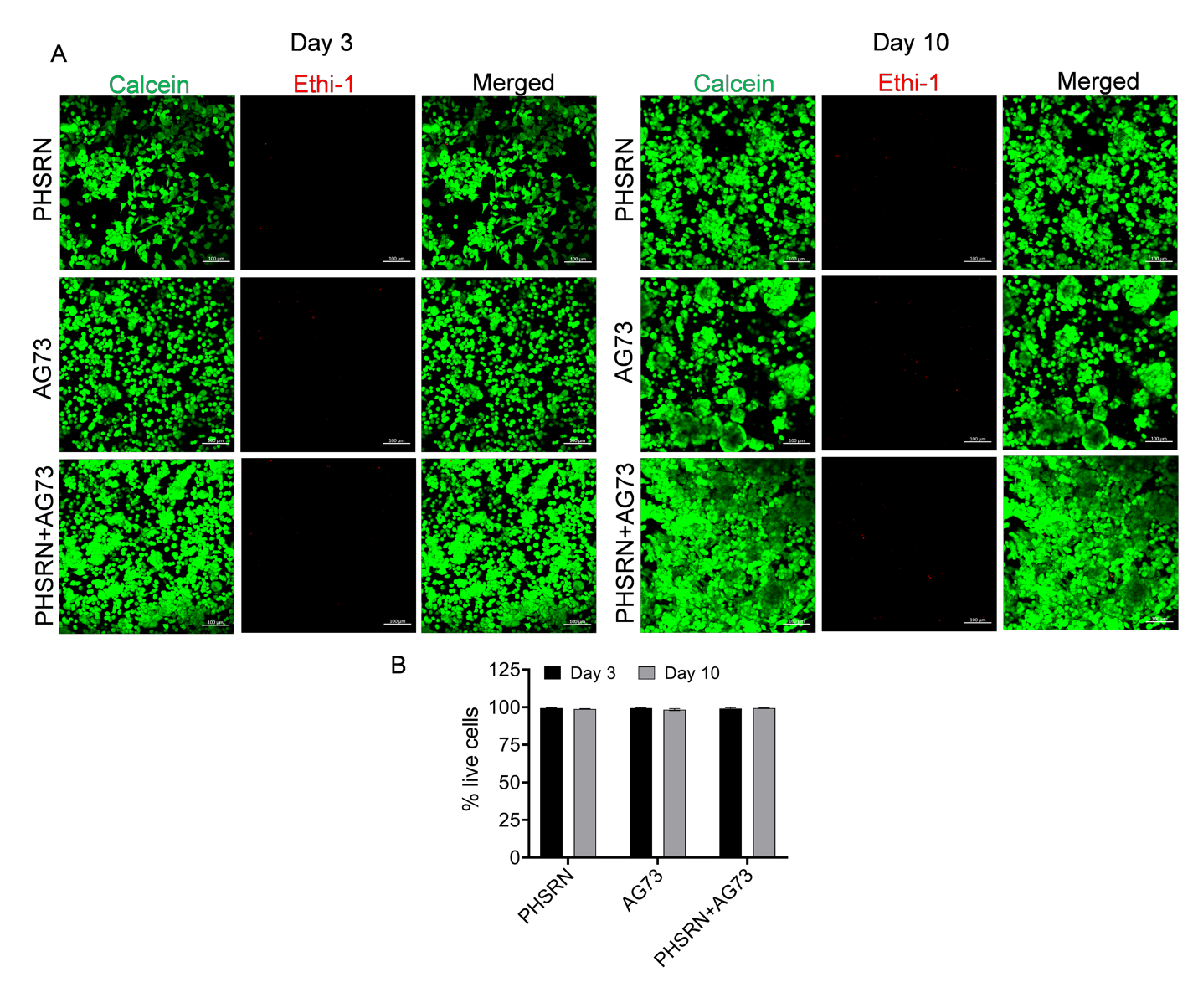

**Figure S7**. Assessment of model epithelial monolayer formation with A549. A) Representative images (individual channels and merged overlay) of A549 cells cultured on different soft synthetic ECM compositions (PHSRN, AG73, or PHSRN+AG73) and stained with live/dead dyes, where the percentage of the surface area covered by cells (Calcein positive area) was used to assess epithelial layer formation at day 3 and day 10 on PHSRN, AG73, or PHSRN+AG73 synthetic ECMs (LSM 800, 10X objective and image frame size of 1024 × 1024, z-projection of z-stack <10 µm with ~2 µm slice interval, Live cells stained with Calcein and dead cells stained with Ethidium homodimer-1 (Ethi-1)). B) Quantification was carried out using Volocity software based on live-dead confocal images. Scale bar = 100 µm. Data shown represent mean ± SD (n=3).

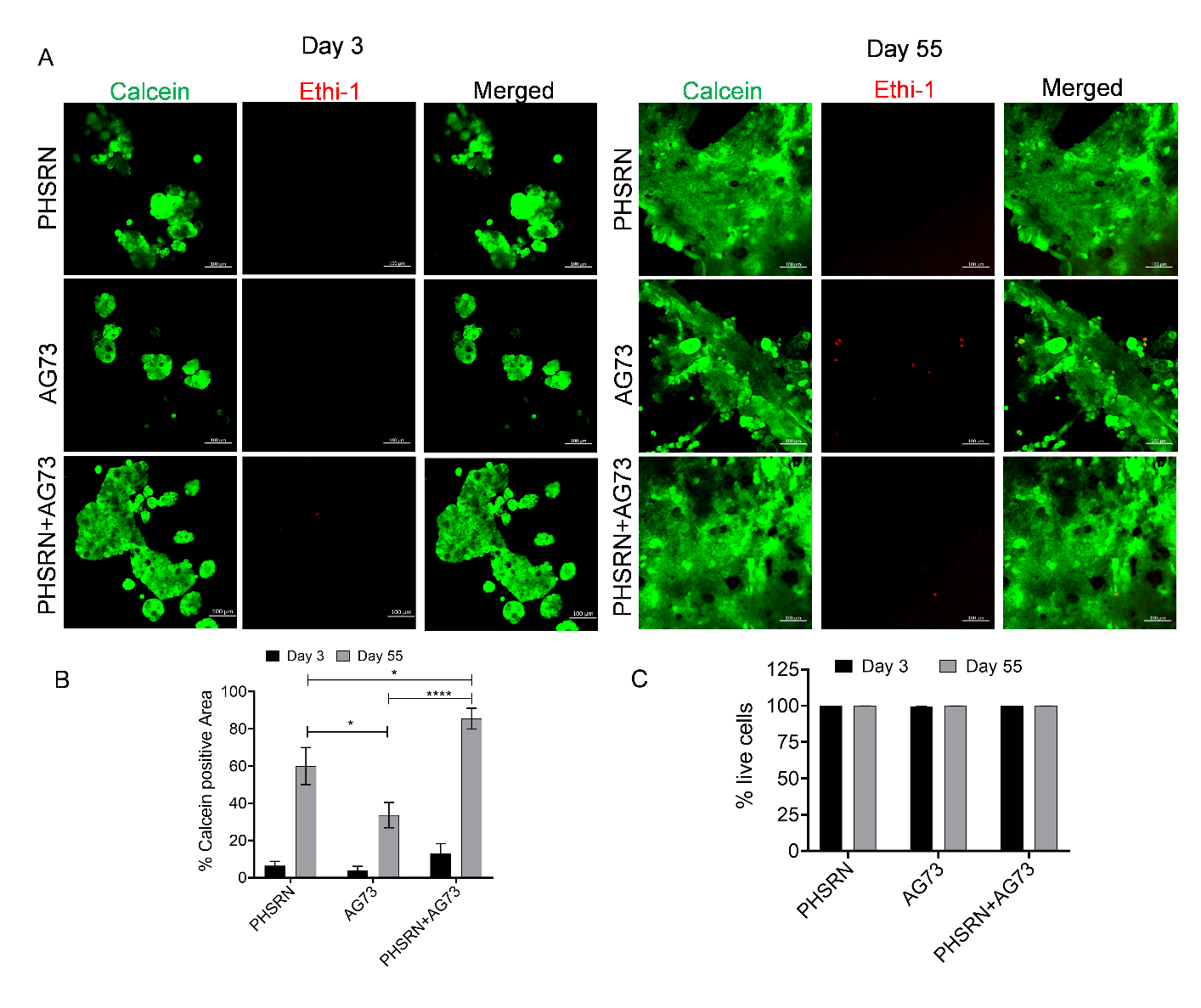

**Figure S8.** Assessment of model epithelial monolayer formation with Calu-3. A) Representative images (individual channels and merged overlay) of Calu-3 cells cultured on soft hydrogels and stained with live/dead dyes, where the percentage of the surface area covered by cells (Calcein positive area) was used to assess epithelial layer formation at day 3 and day 55 on soft PHSRN, AG73, or PHSRN+AG73 synthetic ECMs (LSM 800, 10X objective and image frame size of 1024 × 1024, z-projection of z-stack <10 µm with ~2 µm slice interval, Live cells stained with Calcein and dead cells stained with Ethidium homodimer-1 (Ethi-1)). B) Quantification was carried out using Volocity software based on live-dead confocal images. Scale bar = 100 µm. Significant differences assessed by Student’s two-sided t-test, where differences shown for comparison between conditions at specific timepoints (*p < 0.05, **p < 0.01; ***p < 0.001). Data shown represent mean ± SD (n=3).

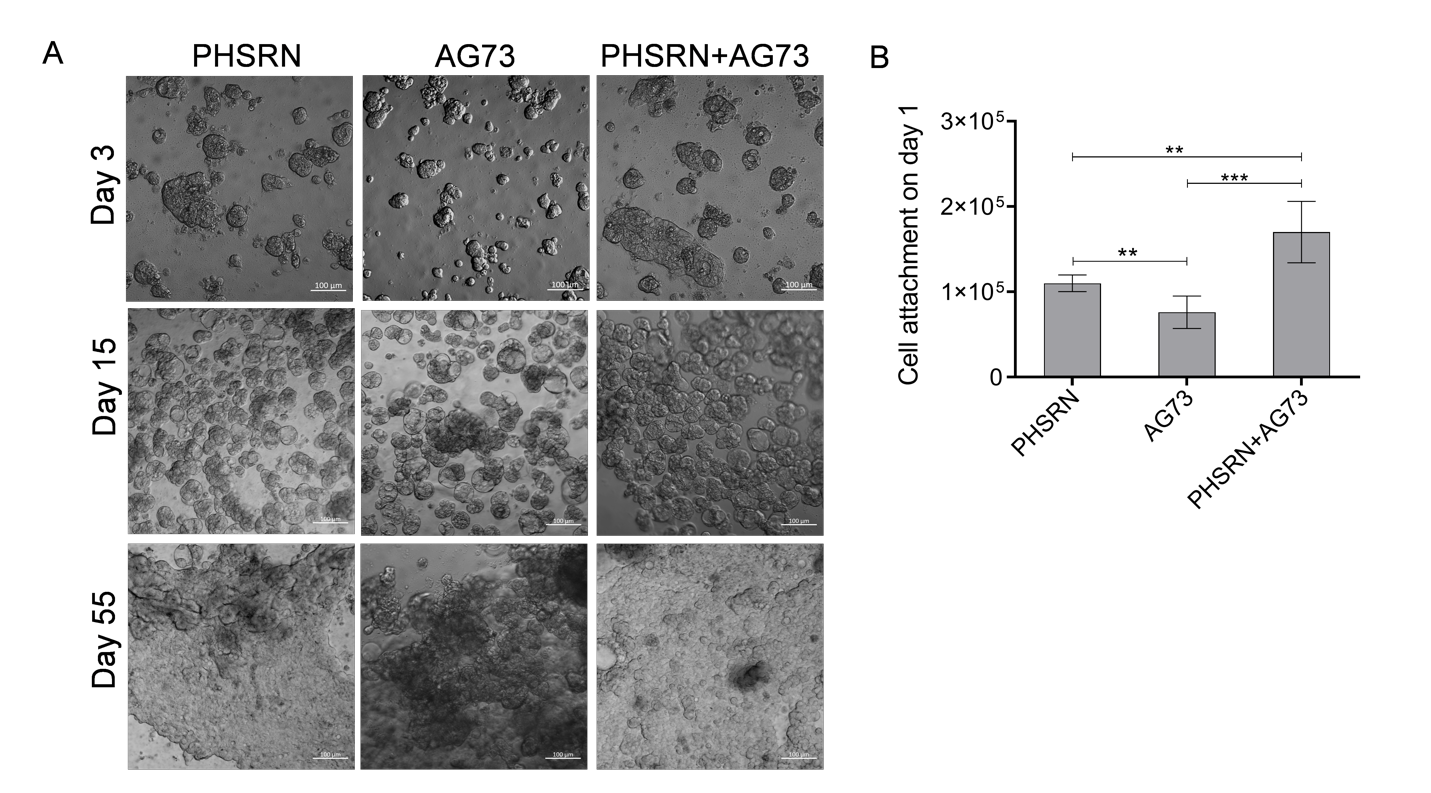

**Figure S9.** Monitoring of attachment and monolayer formation of Calu-3 cells in brightfield. A) Representative images of Calu-3 cells cultured over time from single cell attachment to confluent monolayer on different soft synthetic ECM compositions (PHSRN, AG73, or PHSRN+AG73) (confocal LSM 800, 10X objective and image frame size of 1024 × 1024). B) Quantitative analysis of initial Calu-3 cell attachment at day 1. Significant differences assessed by Student’s two-sided t-test (*p < 0.05, **p < 0.01; ***p < 0.001). Scale bar = 100 µm. Data shown represent mean ± SD (n=3).

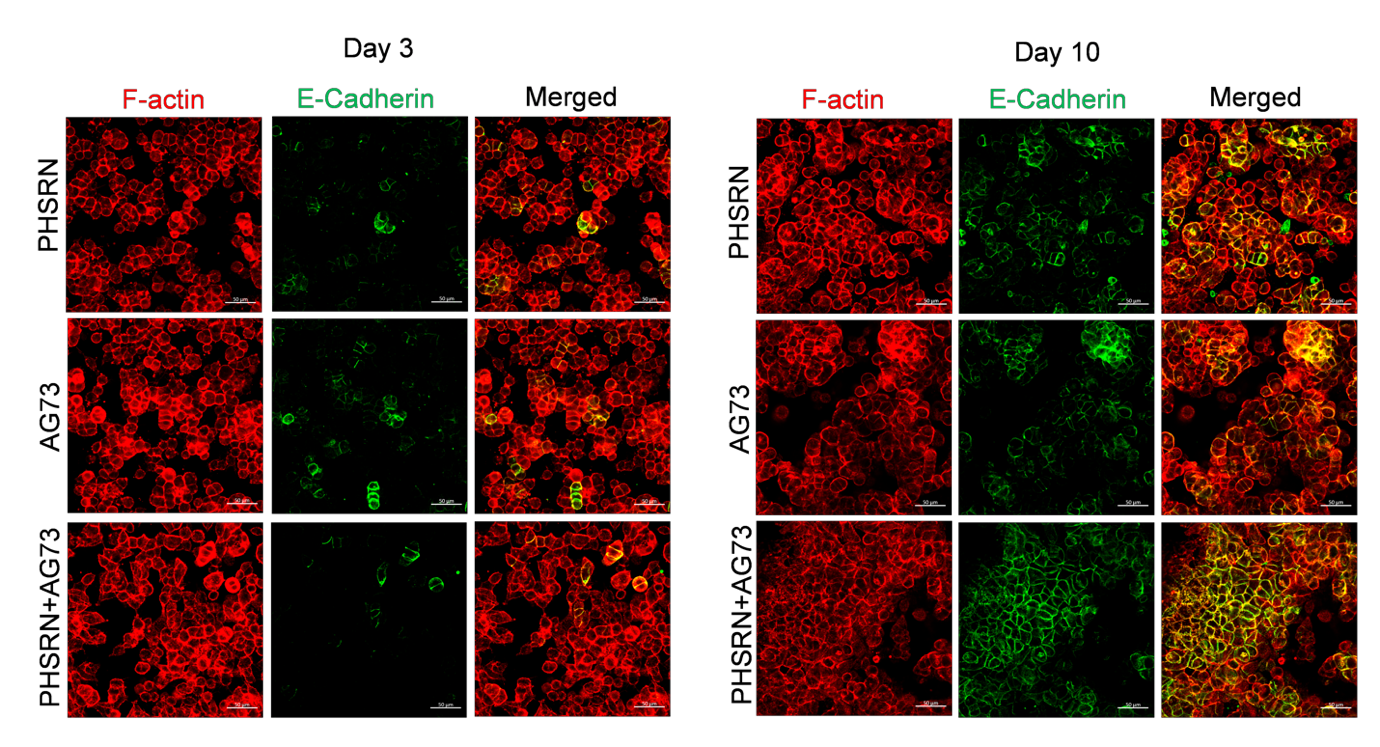

**Figure S10.** Immunofluorescence staining to assess A549 epithelial monolayer formation. Representative images (individual channels and merged overlay) for A549 cultured over time on different soft synthetic ECM compositions (PHSRN, AG73, or PHSRN+AG73), showing the formation of cell-cell adherens junctions with E-cadherin protein expression (green) and development of net-like structure that could be observed with F-actin (red) staining on the PHSRN+AG73 condition relative to PHSRN or AG73 conditions (confocal LSM 800, 10X objective and image frame size of 1024 × 1024, z-projection of z-stack <10 µm with ~2 µm slice interval), demonstrating a synergistic effect of combinations of bioinspired peptides on epithelium formation. Scale bar = 50 µm. Data shown represent mean ± SD (n=3).

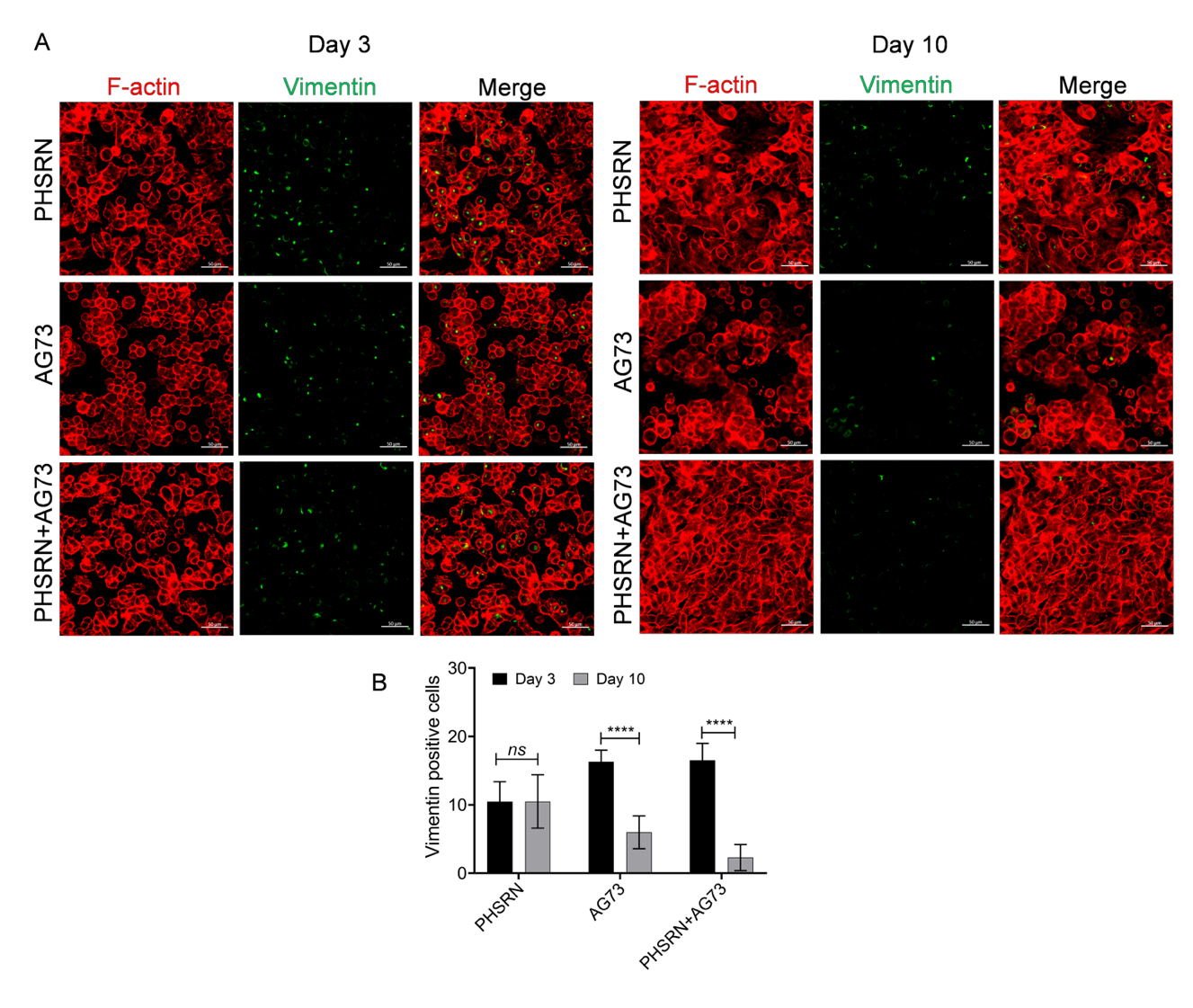

**Figure S11.** Immunofluorescence staining to assess A549 expression of mesenchymal marker. A) Representative images (individual channels and merged overlay) for A549 cultured over time on different soft synthetic ECM compositions (PHSRN, AG73, or PHSRN+AG73) staining for mesenchymal marker Vimentin. A549 cells showed higher expression of a Vimentin at early times culture. At later times, a lower number of Vimentin positive cells was observed along with the development of sheet of connected cells based on cytoskeletal protein F-actin staining on PHSRN+AG73 condition relative to PHSRN or AG73 alone (confocal LSM 800, 10X objective and image frame size of 1024 × 1024, z-projection of z-stack <10 µm with ~2 µm slice interval), demonstrating a synergistic effect of combinations of bioinspired peptides on epithelium formation. B) Quantitative analysis of Vimentin positive cells of (A). Significant differences assessed by Student’s two-sided t-test, where differences shown for comparison between time points (*p < 0.05, **p < 0.01; ***p < 0.001). Scale bar = 50 µm. Data shown represent mean ± SD (n=3).

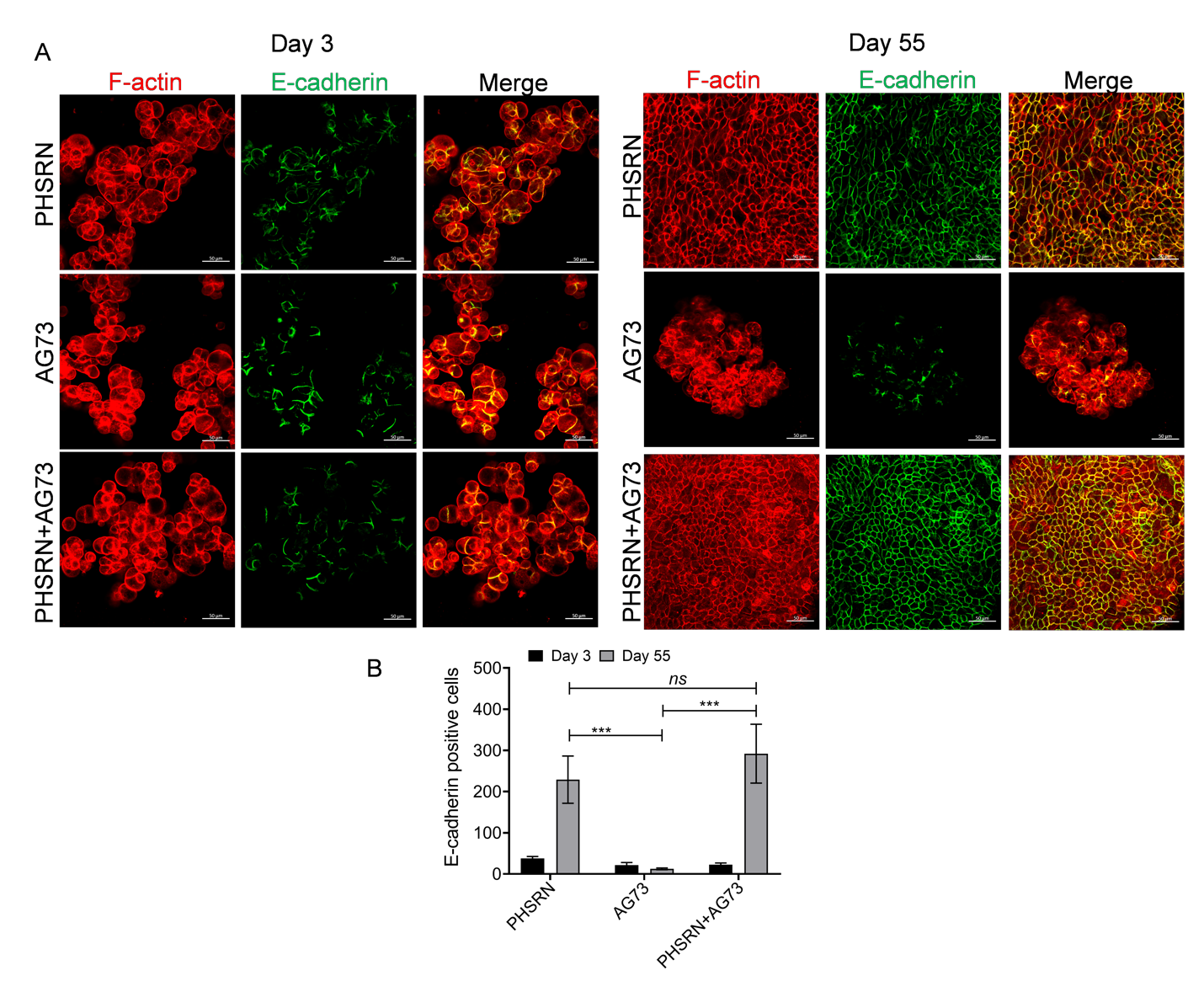

**Figure S12.** Immunofluorescence staining to assess Calu-3 epithelial monolayer formation. A) Representative images (individual channels and merged overlay) for Calu-3 cultured over time on different soft synthetic ECM compositions (PHSRN, AG73, or PHSRN+AG73), showing the formation of cell-cell adherens junctions with E-cadherin protein expression (green) and development of net-like structure that could be observed with F-actin (red) staining on the PHSRN+AG73 condition relative to PHSRN or AG73 conditions (confocal LSM 800, 10X objective and image frame size of 1024 × 1024, z-projection of z-stack <10 µm with ~2 µm slice interval), demonstrating a synergistic effect of combinations of bioinspired peptides on epithelium formation. B) Quantitative analysis of E-cadherin positive cells of (A). Significant differences assessed by Student’s two-sided t-test, where differences shown for comparison between conditions (*p < 0.05, **p < 0.01; ***p < 0.001). Scale bar = 50 µm. Data shown represent mean ± SD (n=3).

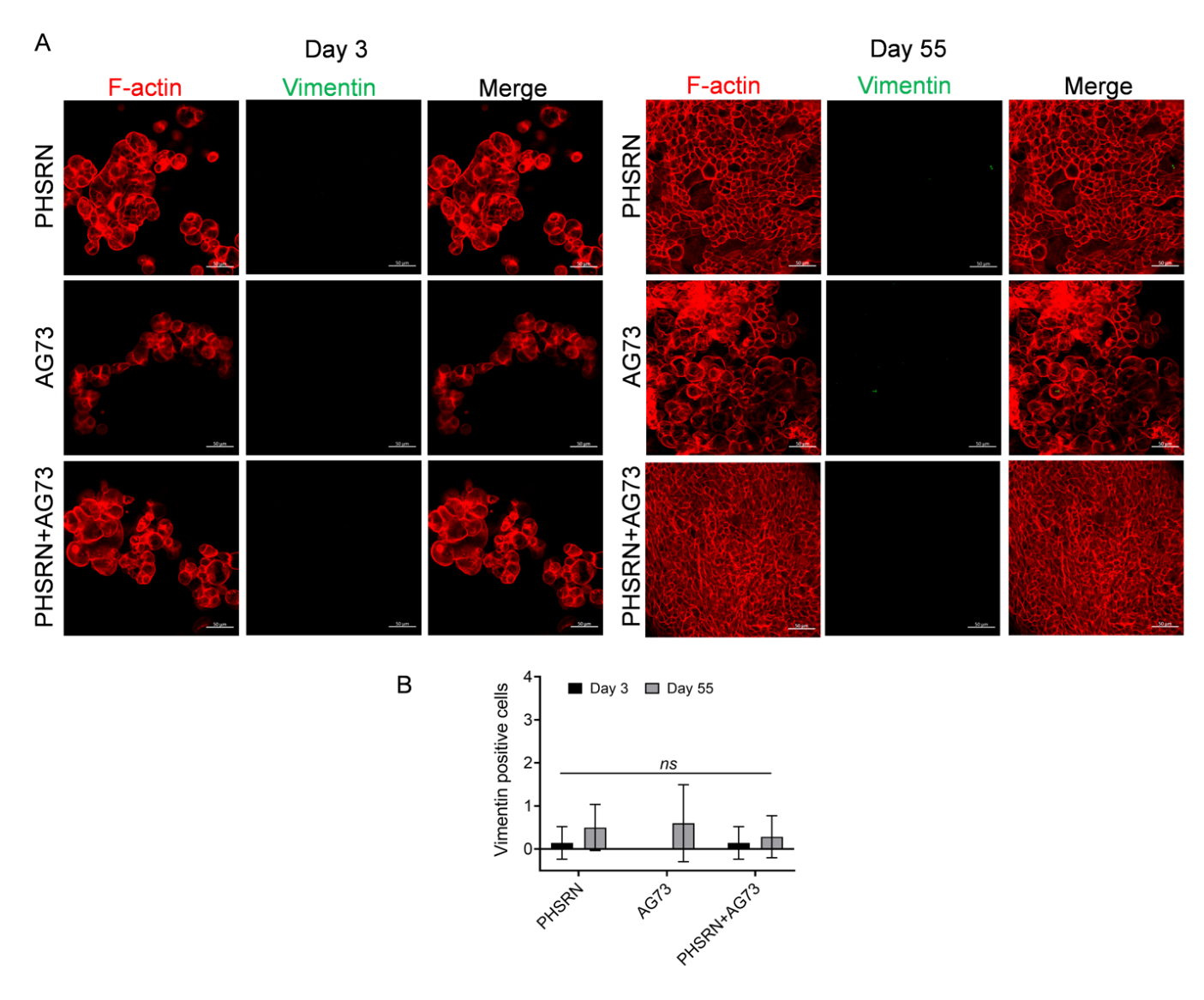

**Figure S13.** Immunofluorescence staining to assess Calu-3 expression of mesenchymal marker. A) Representative images (individual channels and merged overlay) for Calu-3 cultured over time on different soft synthetic ECM compositions (PHSRN, AG73, or PHSRN+AG73) staining for mesenchymal marker Vimentin (confocal LSM 800, 20X objective and image frame size of 1024 × 1024, z-projection of z-stack <10 µm with ~2 µm slice interval, stained for F-actin and Vimentin, Scale bar = 50 µm). B) Quantitative analysis of Vimentin positive cells of (A). Significant differences assessed by Student’s two-sided t-test, where differences shown for comparison between conditions (*p < 0.05, **p < 0.01; ***p < 0.001). Data shown represent mean ± SD (n=3).

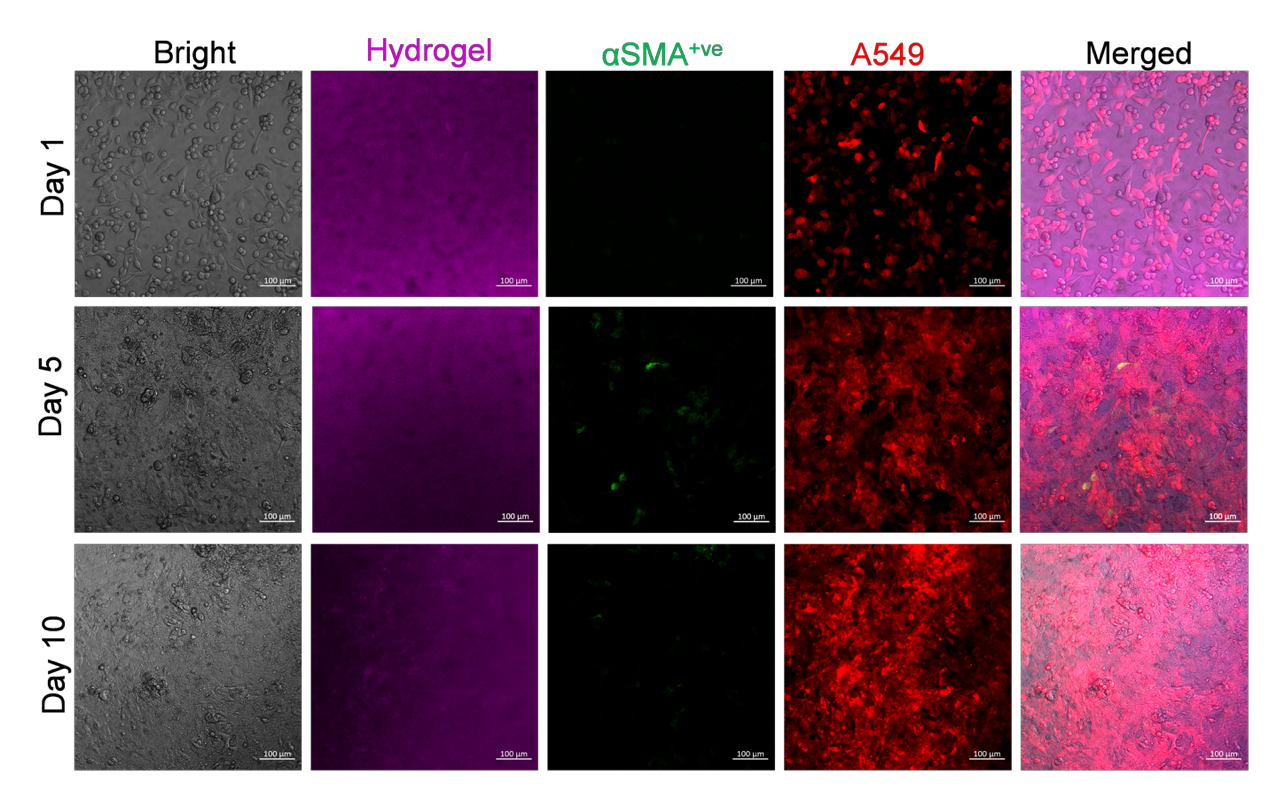

**Figure S14.** Epithelium formation *in situ* using reporter A549 cells. Representative images (individual channels and merged overlay) for stable reporter AF549 cells cultured on the soft PHSRN+AG73 condition over time (confocal LSM 800, 10X objective and image frame size of 1024 × 1024, constitutive expression of a red fluorescent protein, conditional expression of a green fluorescent protein when αSMA is expressed, hydrogel labeled with AF647, Scale bar = 100 µm, n=3).

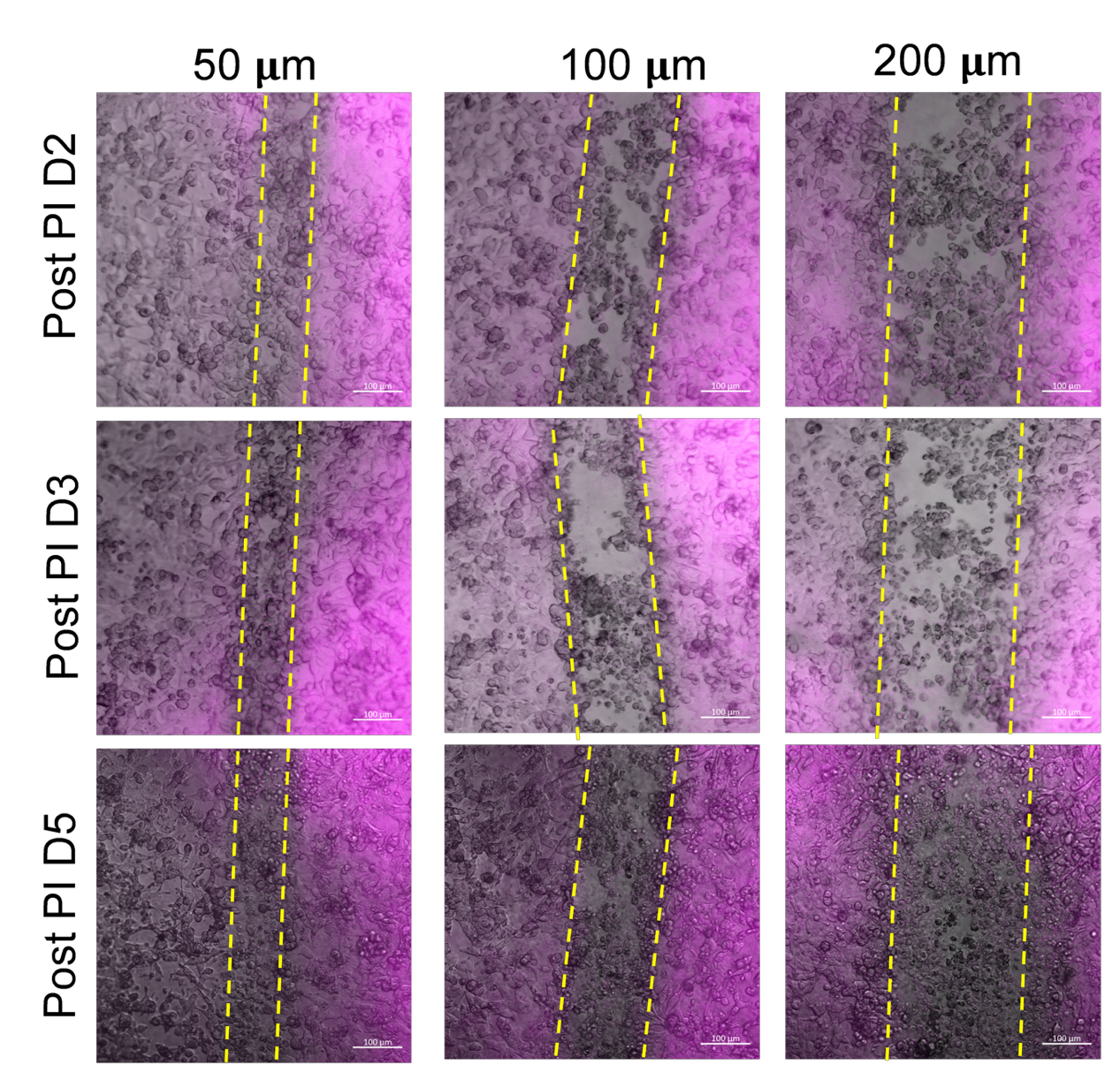

**Figure S15.** Representative images of recovery after photo-injury (PI) using A549 cells. Photomask with different ‘slit’ widths (50, 100, 200 µm; 6.75 mW cm^-2^ at 430 nm for 4 min) were used to create injury on the soft PHSRN+AG73 synthetic ECM (labeled with AF647), and cell response monitored in brightfield over time (confocal LSM 800, 10X objective and image frame size of 1024 × 1024, z-projection of z-stack <10 µm with ~2 µm slice interval, brightfield, magenta hydrogel AF647, injury edges denoted with dashed yellow lines, n=3). Scale bar = 100 µm.

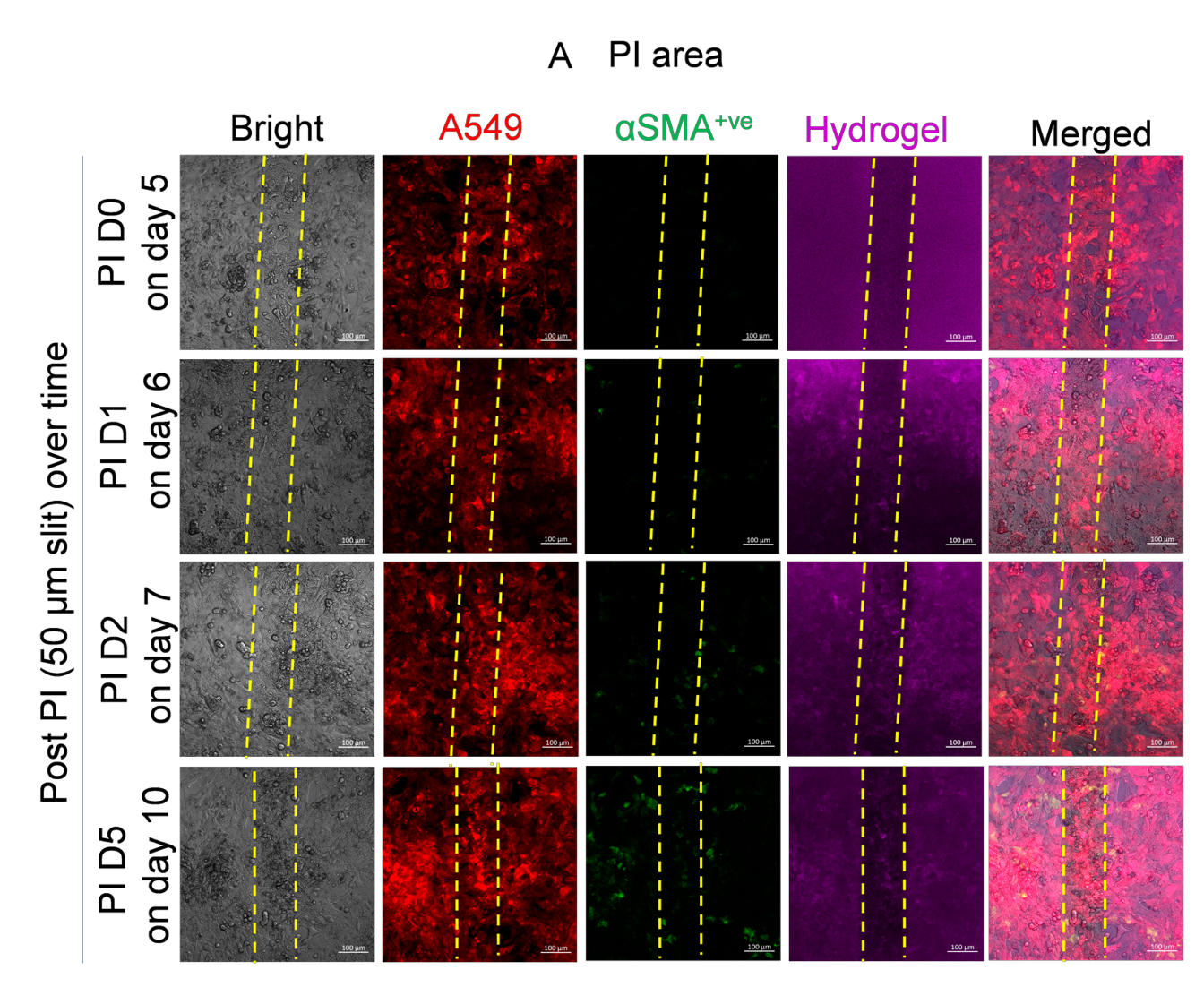

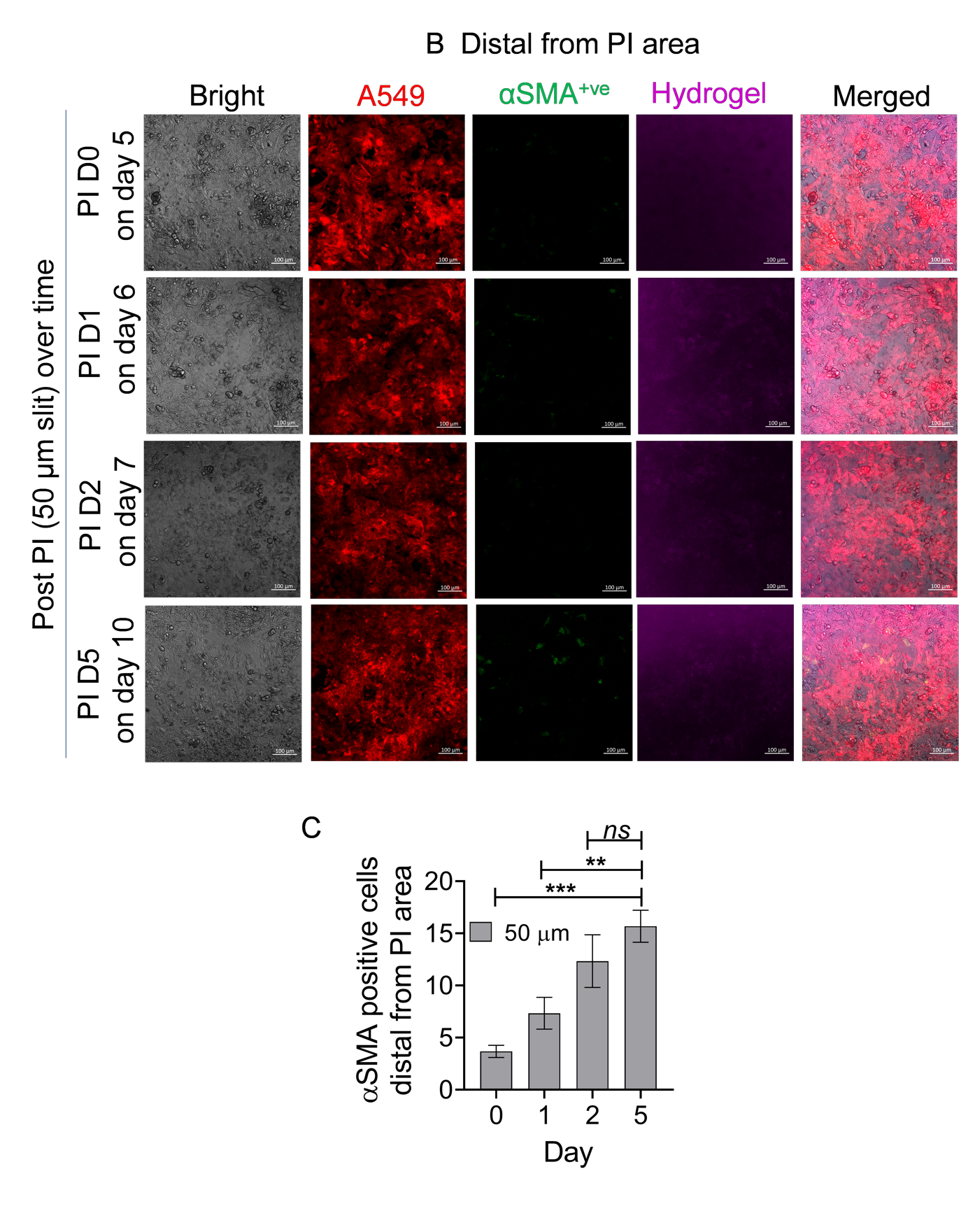

**Figure S16.** Response to injury (50 µm) using reporter A549 cells. A) Representative *in situ* images of epithelial cell recovery using reporter A549 cells either A) proximal to PI (50 µm) area or B) distal from PI area over time (confocal LSM 800, 10X objective and image frame size of 1024 × 1024, z-projection of z-stack <10 µm with ~2 µm slice interval, constitutive expression of a red fluorescent protein, conditional expression of a green fluorescent protein when is αSMA expressed, hydrogel labeled with AF647). C) Quantitative analysis of (B). Significant differences assessed by Student’s two-sided t-test, where differences shown for comparison between day timepoints in PHSRN+AG73 conditions (*p < 0.05, **p < 0.01; ***p < 0.001). Data shown represent mean ± SD (n=3). Scale bar = 100 µm.

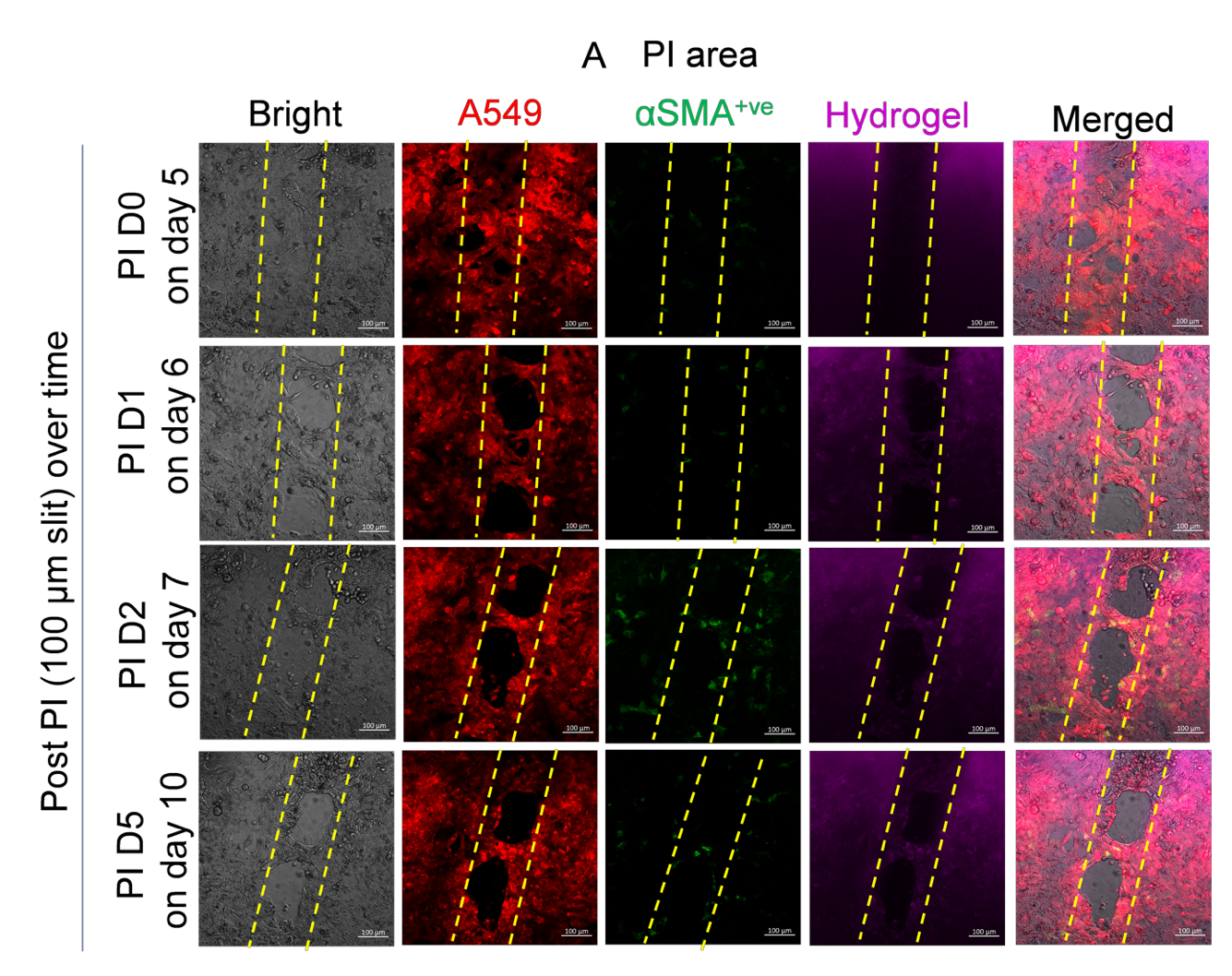

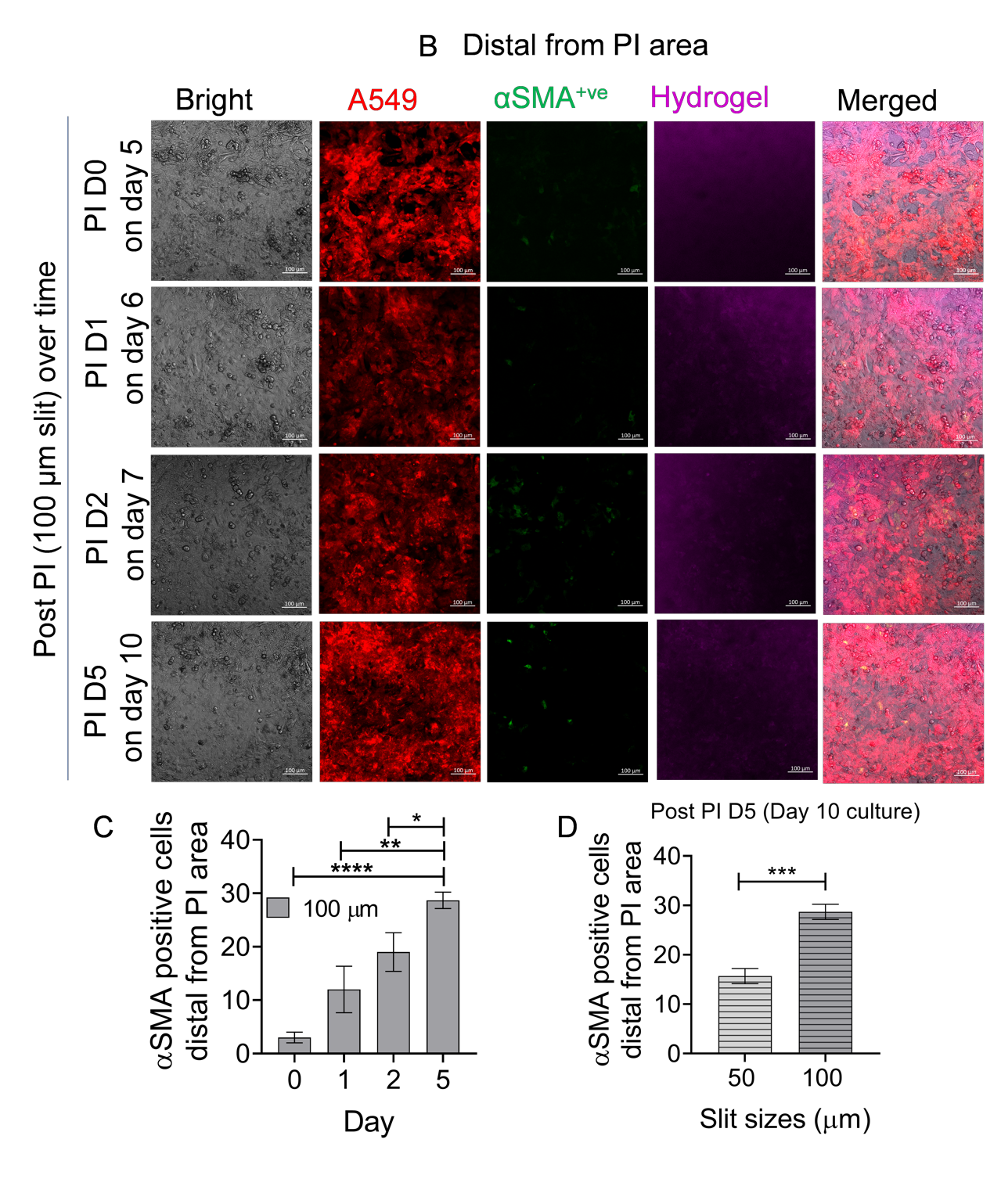

**Figure S17.** Response to injury (100 µm) using reporter A549 cells with comparison to 50 µm. A) Representative *in situ* images of epithelial cell recovery using reporter A549 cells either A) proximal to PI (100 µm) area or B) distal from PI area over time (confocal LSM 800, 10X objective and image frame size of 1024 × 1024, z-projection of z-stack <10 µm with ~2 µm slice interval, constitutive expression of a red fluorescent protein, conditional expression of a green fluorescent protein when is αSMA expressed, hydrogel labeled with AF647). C) Quantitative analysis of (B). D) Comparison of αSMA protein expression distal from PI area on day 5 post PI (Day 10 culture) between 50 and 100 µm size injury. Significant differences assessed by Student’s two-sided t-test, where differences shown for comparison between day timepoints in PHSRN+AG73 conditions (*p < 0.05, **p < 0.01; ***p < 0.001). Data shown represent mean ± SD (n=3). Scale bar = 100 µm.

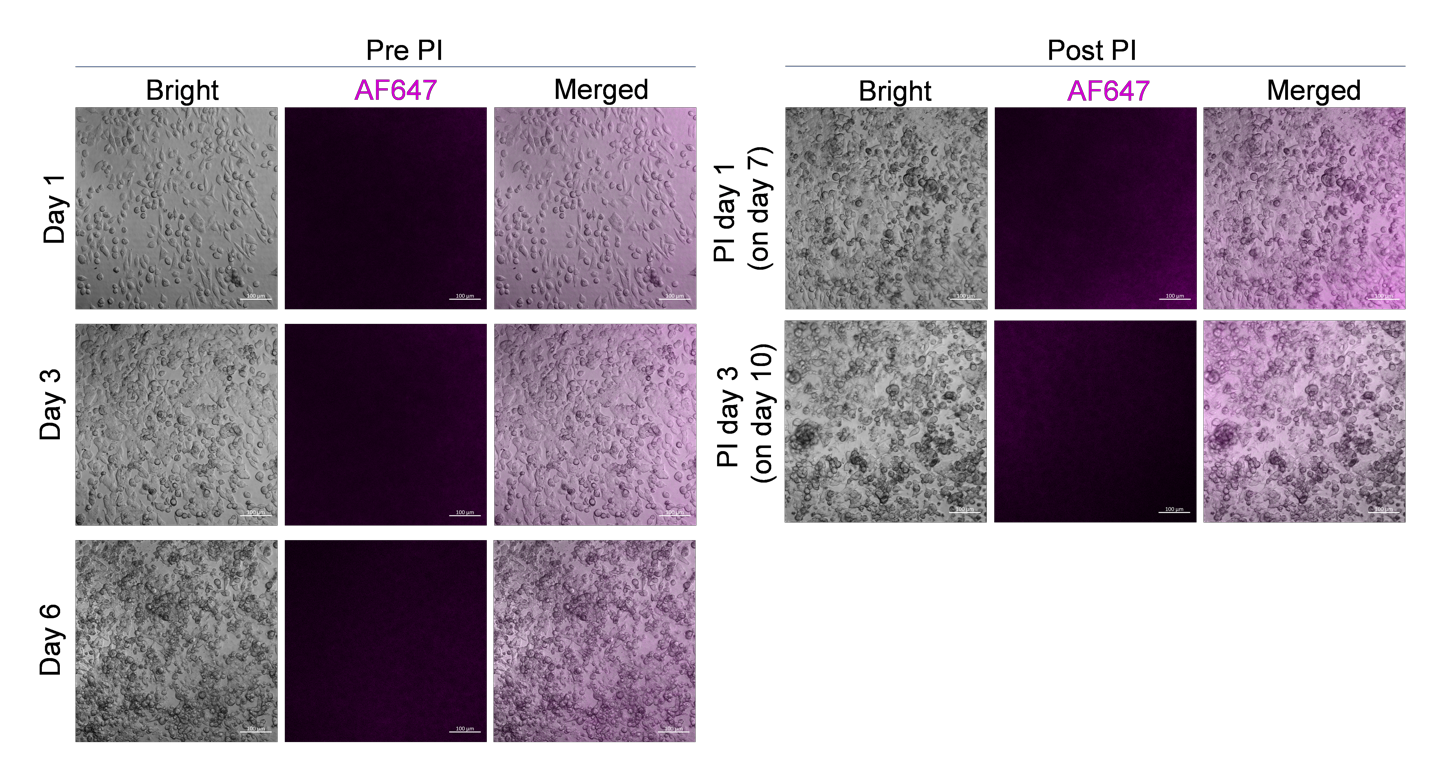

**Figure S18**. Irradiation of cells on a non-degradable synthetic ECM. Representative images (individual channels and merged overlay) of A549 cells on non-degradable hydrogel (PEG-4-BCN linked with PEG-2-azide and integrating PHSRN+AG73) before (pre) and after (post) light exposure (confocal images captured by LSM 800, 10X objective, image frame size of 1024 × 1024, z-projection of z-stack <10 µm with ~2 µm slice interval, brightfield, magenta hydrogel labeled with AF647). Scale bar = 100 µm.

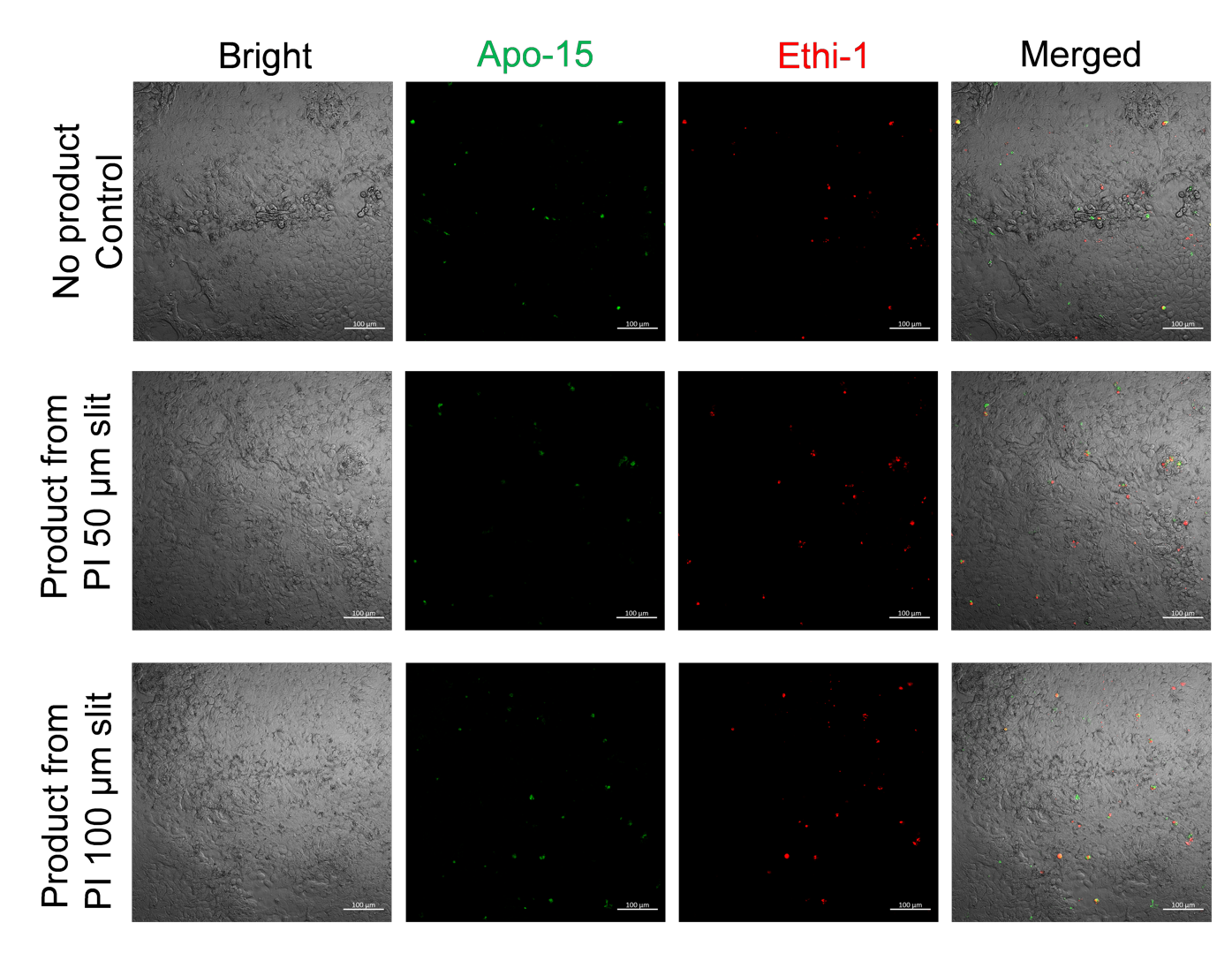

**Figure S19**. Exposure of cells to photodegradation products. Representative images (individual channels and merged overlay) of A549 cells in a 2D culture with 48 hours of exposure to the degradation products from a synthetic ECM. Apo-15 (green) was used to identify apoptotic cells, while Ethidium homodimer-1 (Ethi-1, red) was used for necrotic cells. Confocal images captured by LSM 800 using 10X objective (image frame size of 1024 × 1024, z-projection of z-stack <10 µm with ~2 µm slice interval). Scale bar = 100 μm.

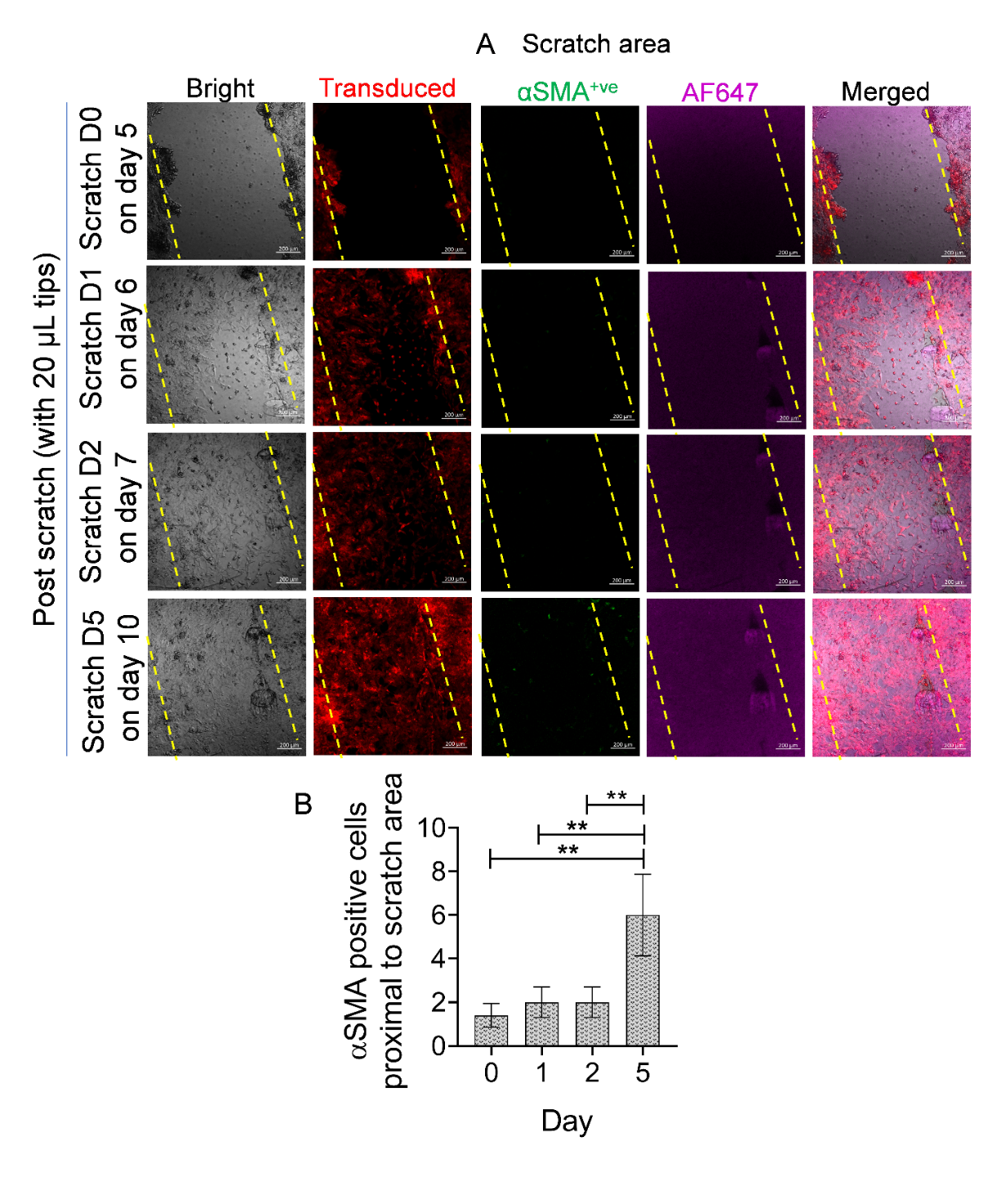

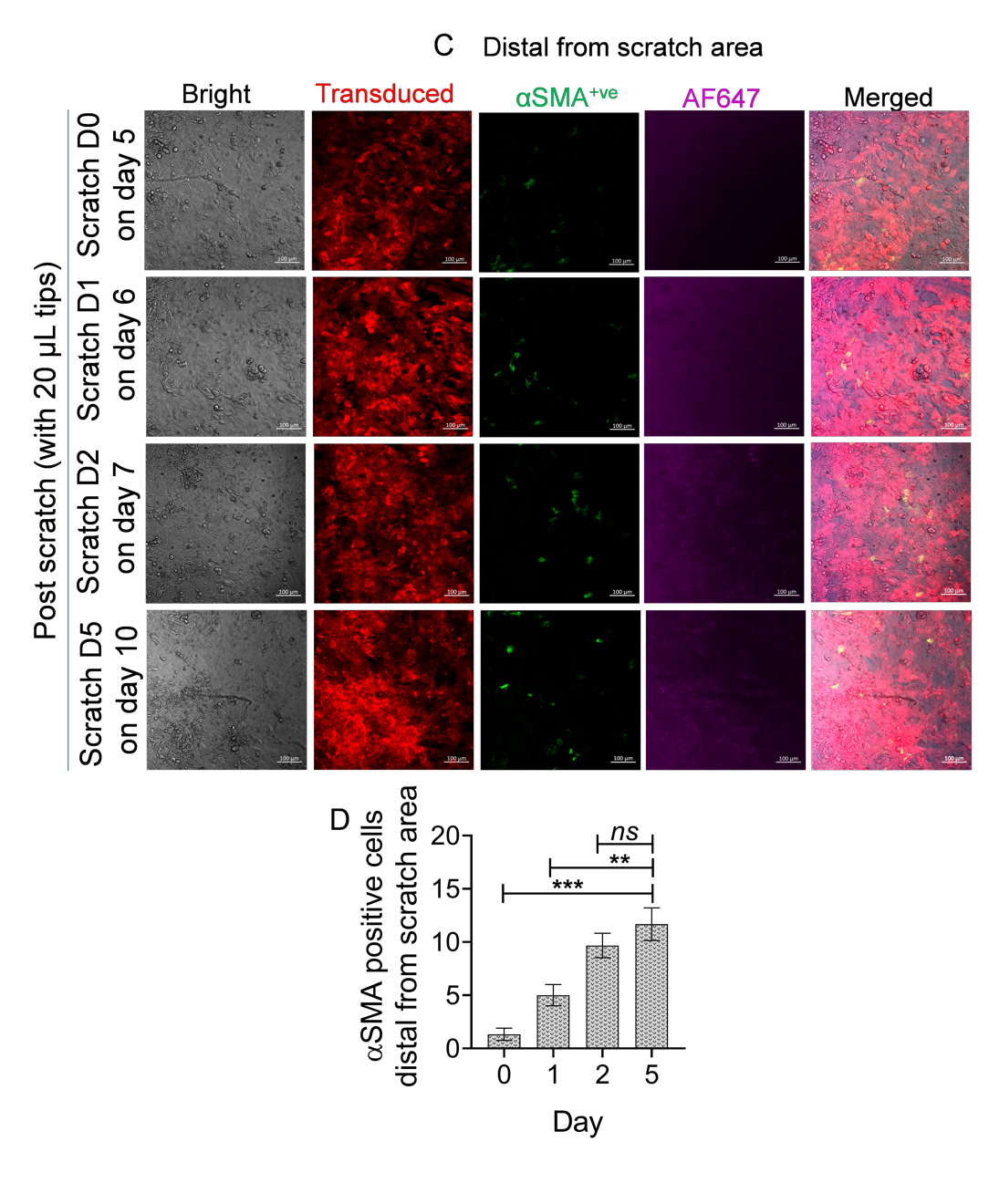

**Figure S20.** Response to mechanical scratch on synthetic ECM using reporter A549 cells. Representative *in situ* images of epithelial cell recovery using reporter A549 cells either A) proximal to a mechanical scratch (made with a pipette tip) with B) quantitative analysis or C) distal to the scratch area with D) quantitative analysis (confocal LSM 800, 10X objective and image frame size of 1024 × 1024, z-projection of z-stack <10 µm with ~2 µm slice interval, constitutive expression of a red fluorescent protein, conditional expression of a green fluorescent protein when is αSMA expressed, hydrogel labeled with AF647). Significant differences assessed by Student’s two-sided t-test, where differences shown for comparison between day timepoints with scratch on PHSRN+AG73 condition (*p < 0.05, **p < 0.01; ***p < 0.001). Data are shown represent mean ± SD (n=3). Scale bar = 100 µm.

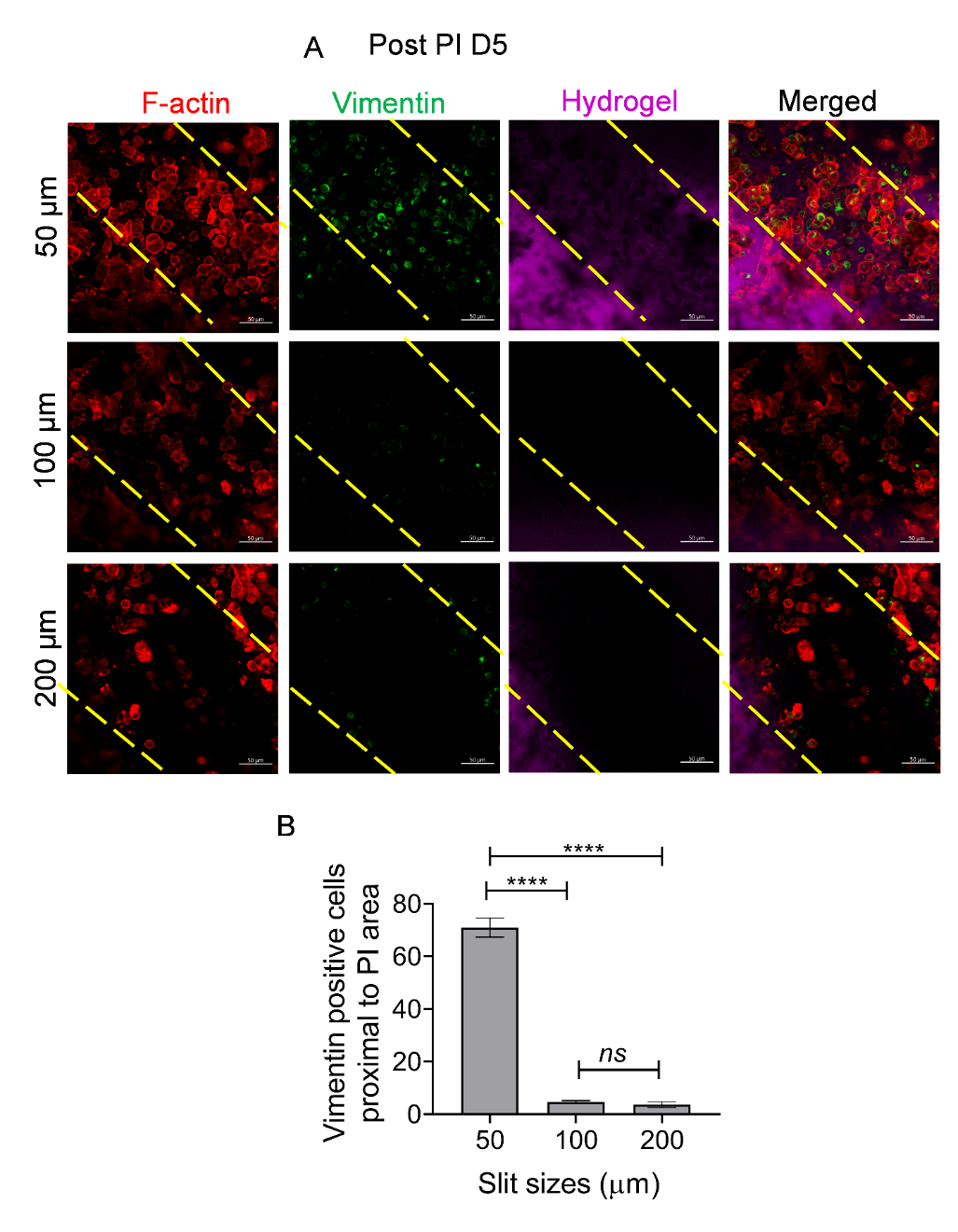

**Figure S21**. Immunostaining for mesenchymal marker proximal to photo-injuries of different widths. A) Representative images (individual channels and merged overlay) of A549 cells on the photoresponsive synthetic ECM with different sizes of photo-injury (PI) and proximal to the PI (confocal LSM 800, 10X objective and image frame size of 1024 × 1024, z-projection of z-stack <10 µm with ~2 µm slice interval, red F-actin cytoskeletal marker, green Vimentin mesenchymal marker, magenta hydrogel labeled with AF647, boundaries of ‘injury’ noted with yellow dashed lines, Scale bar = 50 µm). B) Quantitative analysis of cells positive for Vimentin in (A). Statistical differences determined Student’s two-sided t-test, where differences shown for comparison between conditions (*p < 0.05, **p < 0.01; ***p < 0.001). Data shown represent mean ± SD (n=3).

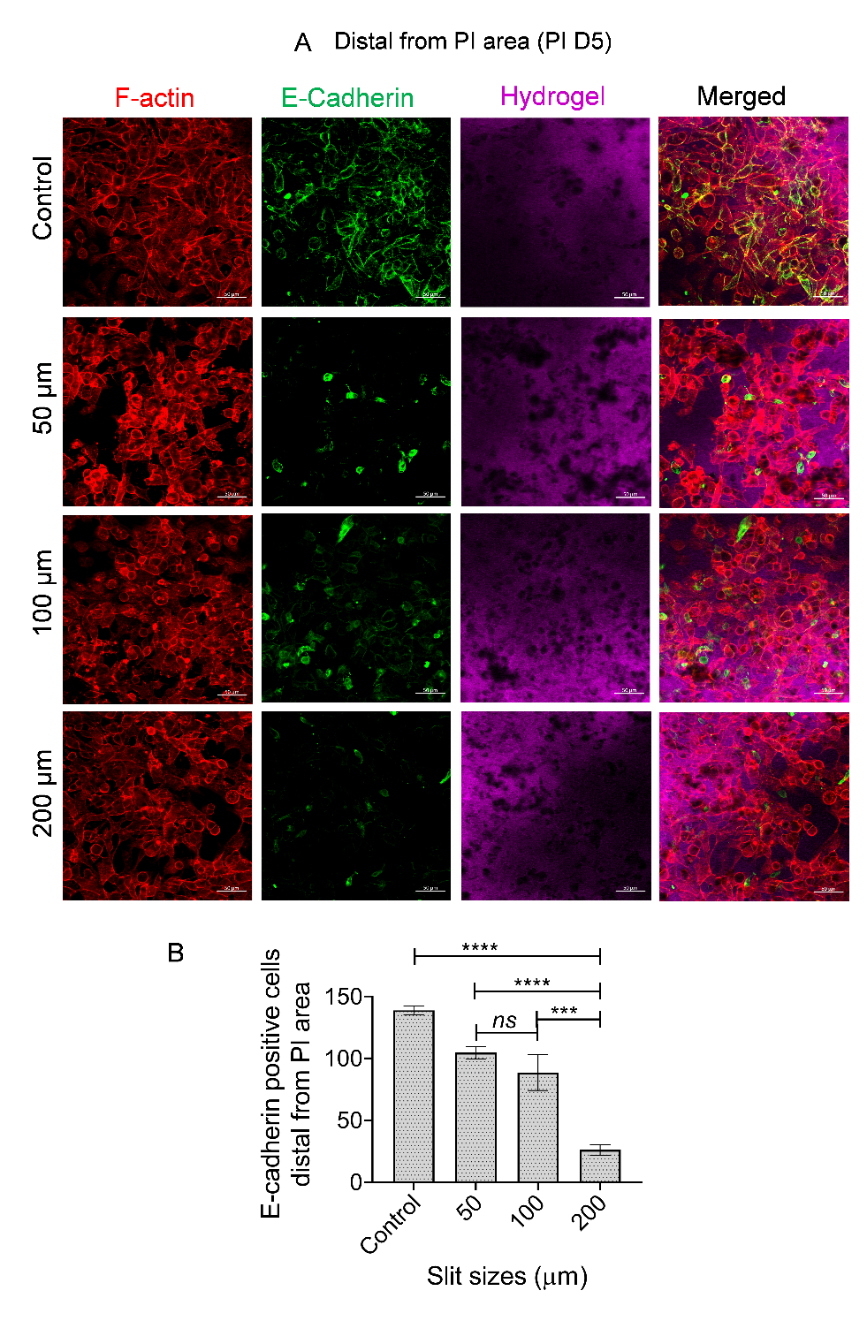

**Figure S22**. Immunostaining for epithelial marker distal to photo-injuries of different widths. A) Representative images (individual channels and merged overlay) of A549 showed overall loss of the E-cadherin expression for cells that distal from PI area ( confocal LSM 800, 10X objective and image frame size of 1024 × 1024, z-projection of z-stack <10 µm with ~2 µm slice interval, epithelial marker E-cadherin stained green, cytoskeletal marker F-actin stained red, hydrogel labeled with AF647, Scale bar = 50 µm). B) Quantitative analysis of cells positive for E-cadherin protein in (A). Statistical differences determined Student’s two-sided t-test, where differences shown for comparison between conditions (*p < 0.05, **p < 0.01; ***p < 0.001). Data shown represent mean ± SD (n=3).

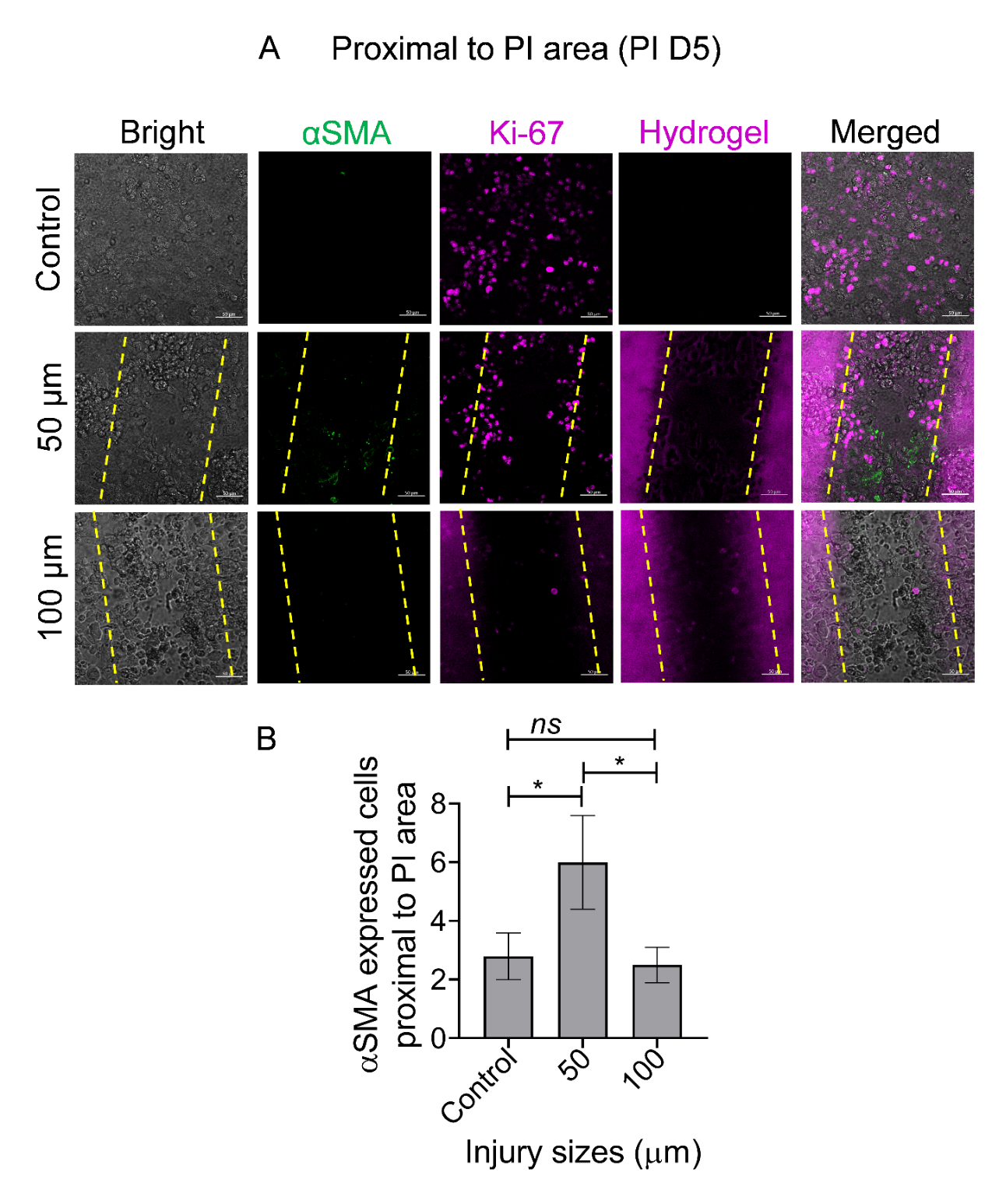

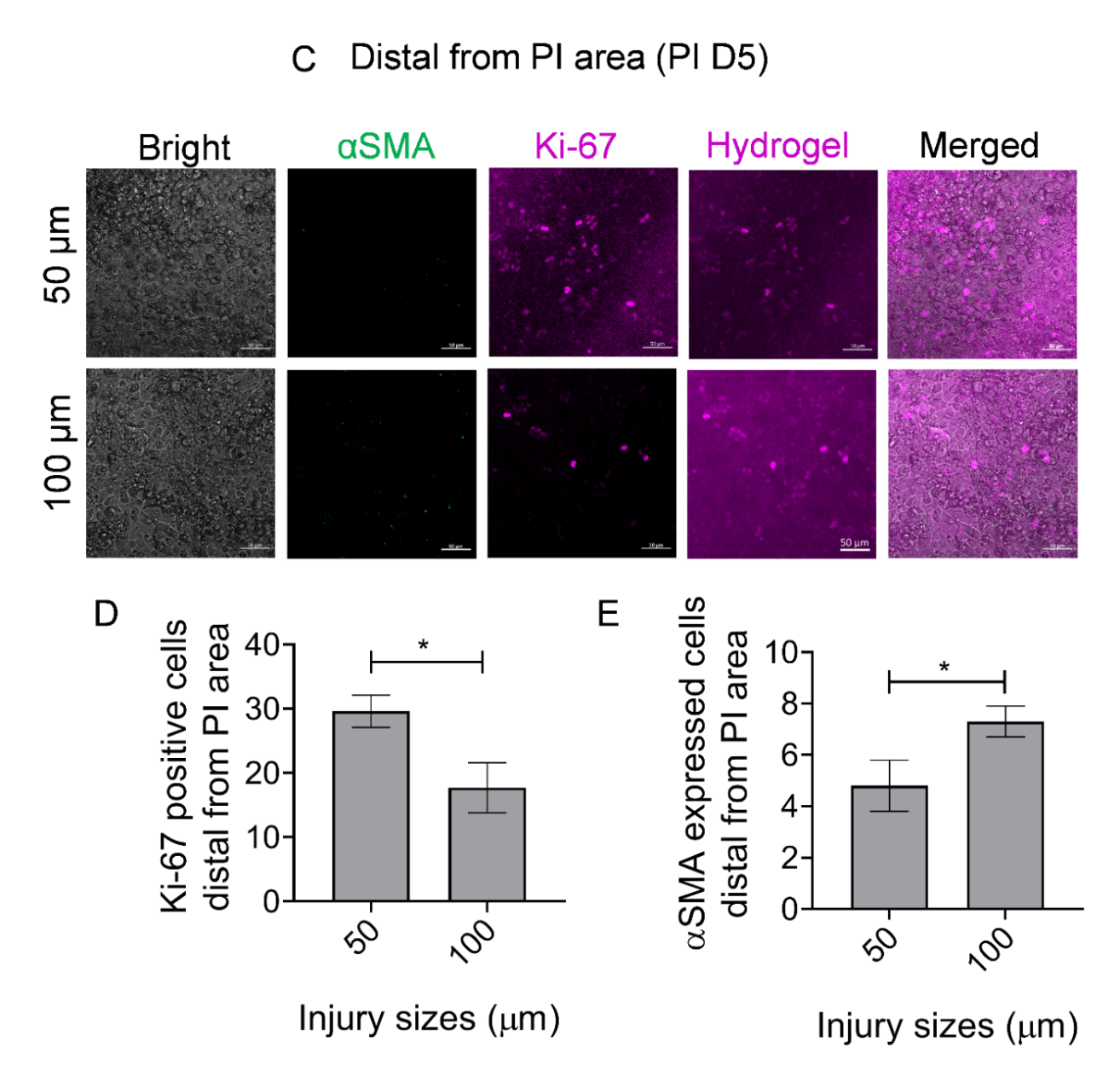

**Figure S23**. Immunostaining for phenotypic and proliferation markers proximal and distal to photo-injuries of different injury widths. A) Representative confocal images of A549 responses on the photoresponsive synthetic hydrogel proximal to with different sizes of photo-injury (PI) in comparison to controls (confocal LSM 800, 10X objective and image frame size of 1024 × 1024, z-projection of z-stack <10 µm with ~2 µm slice interval, Ki-67-positive magenta, αSMA stained green, F-actin stained red, AF647 stained hydrogel, Scale bar = 50 µm). Here, all individual channels are shown for the composite images that are displayed in Figure 4 for full visualization. B) Quantitative analysis of cells positive for αSMA marker expression of (A, proximal to PI). C) Representative images for distal from PI area (50, 100 µm slits). D) Quantitative analysis of cells positive for Ki-67 (magenta) marker expression of (C). E) Quantitative analysis of cells positive for αSMA marker expression of (C). Statistical differences determined Student’s two-sided t-test, where differences shown for comparison between conditions (*p < 0.05, **p < 0.01; ***p < 0.001). Data are shown represent mean ± SD (n=3).

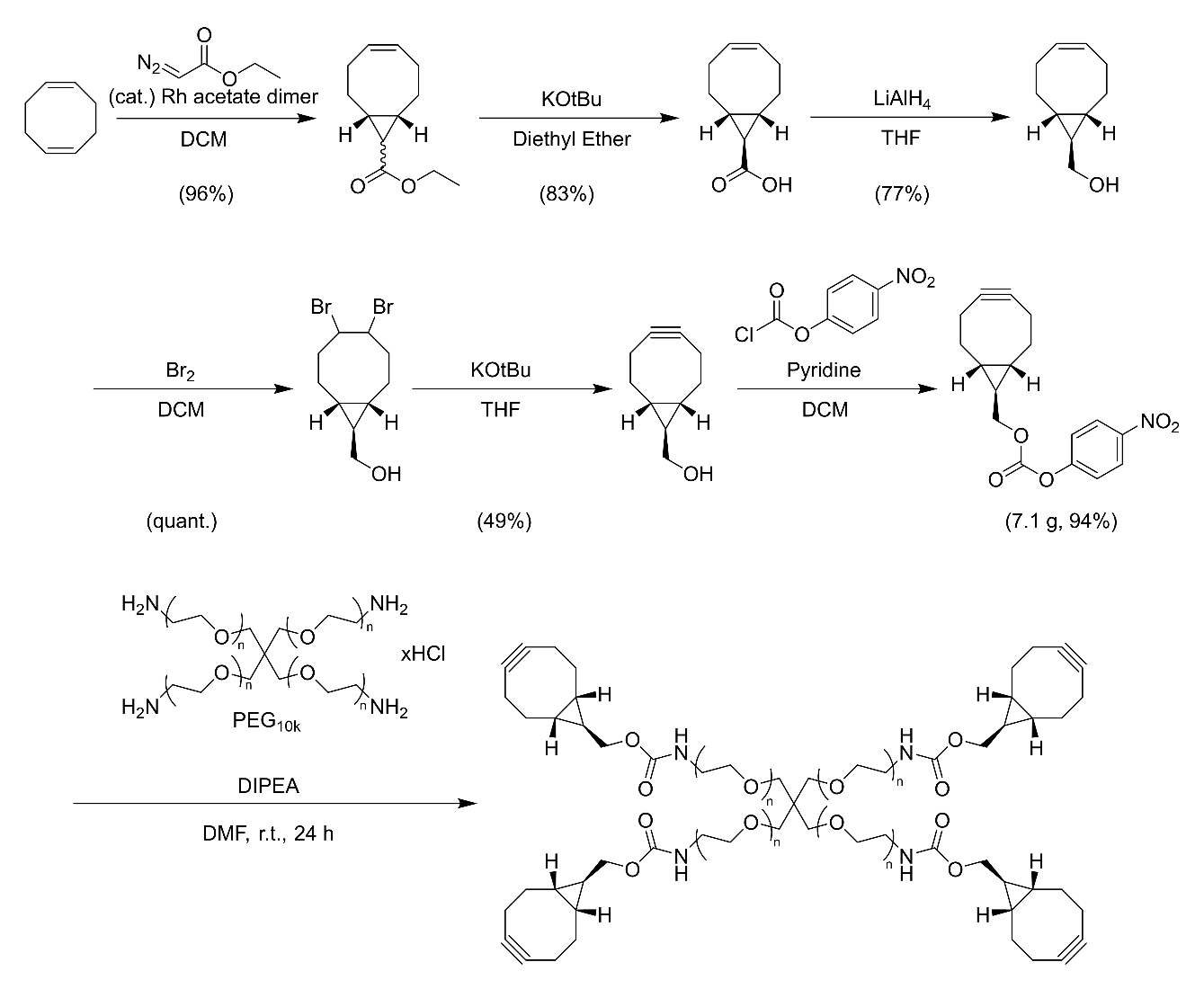

**Figure S24.** Synthetic scheme for PEG-4-BCN (exo).

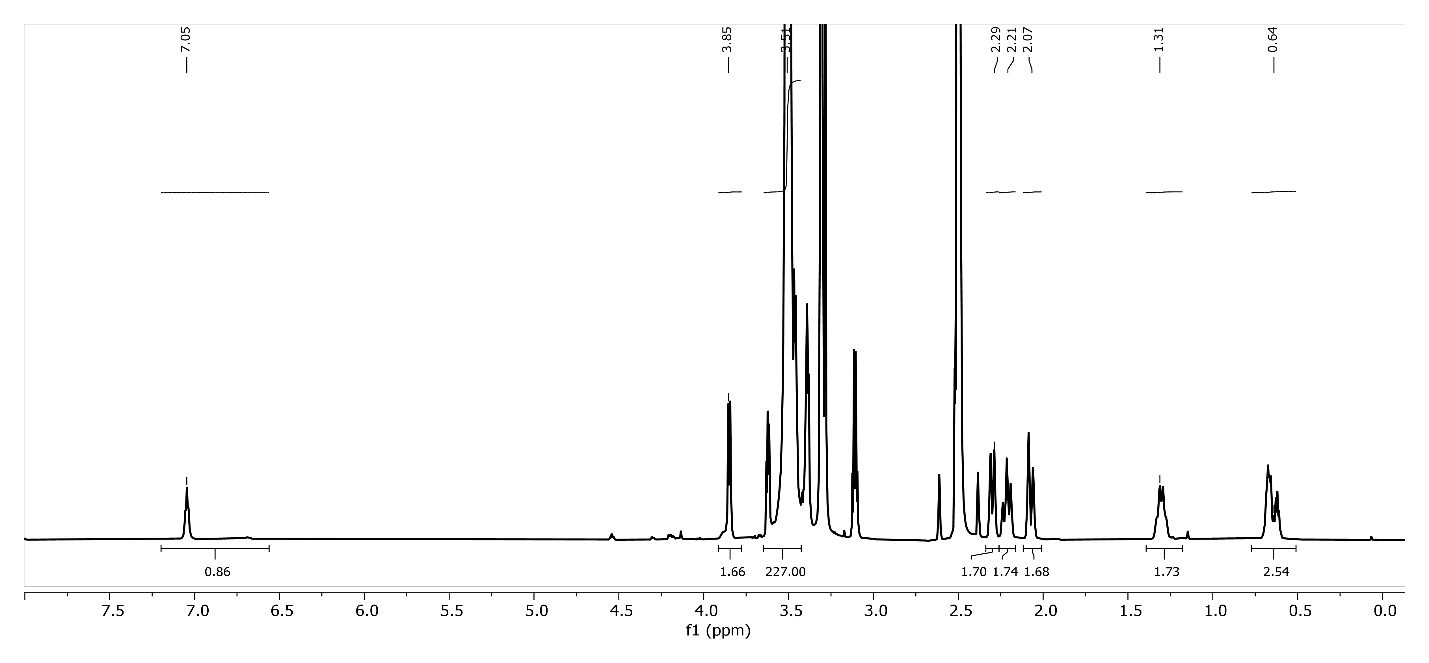

**Figure S25.** NMR of PEG-4-BCN (exo). Representative ^1^H NMR spectra (Bruker AVIII600; Bruker Corporation, Billerica, MA) for PEG-*tetra*-bicyclononyne (600 MHz, DMSO). Functionality was confirmed via integration normalization to one PEG arm and comparing integral values to characteristic BCN ring peaks (δ 2.29, 2.21, 2.07) and the PEG backbone (δ 3.5).

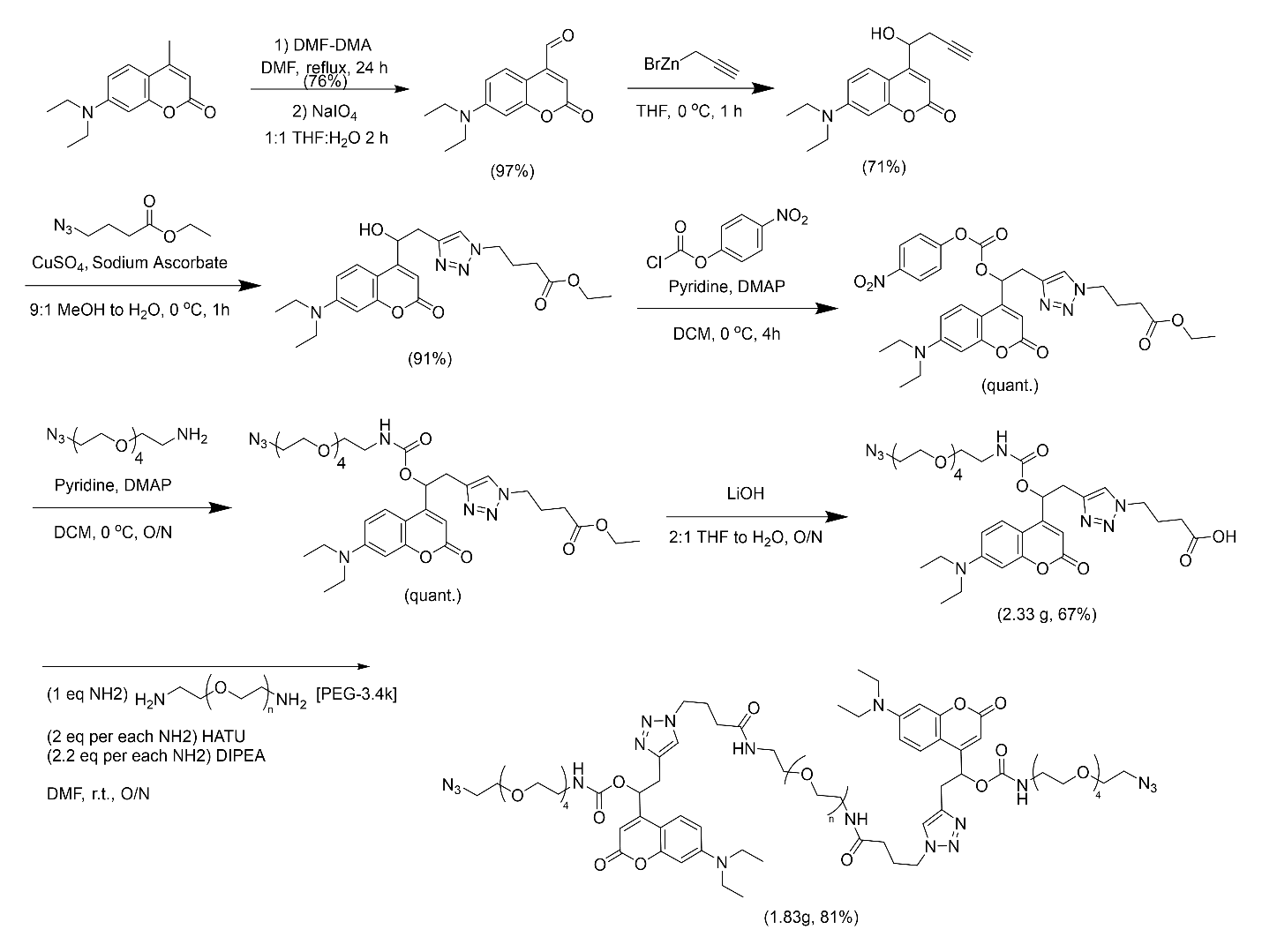

**Figure S26.** Synthetic scheme for PEG-2-CmPN_3_.

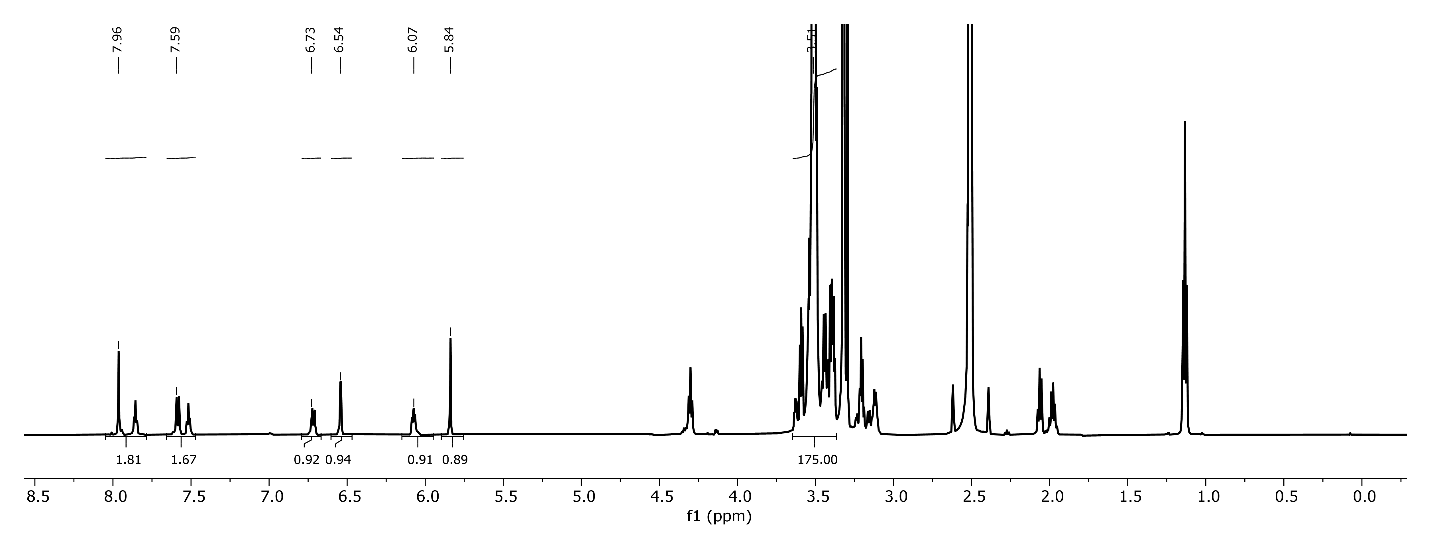

**Figure S27.** NMR of PEG-2-CmPN_3_. Representative ^1^H NMR (Bruker AVIII600; Bruker Corporation, Billerica, MA) spectra for (PEG)-*bis-*coumarin-azide (600 MHz, DMSO). Functionality was confirmed via integration normalization to one PEG arm and comparing integral values to characteristic coumarin peaks (δ 7.96, 7.59, 6.73, 6.54, 6.07, 5.84) and the PEG backbone (δ 3.5).

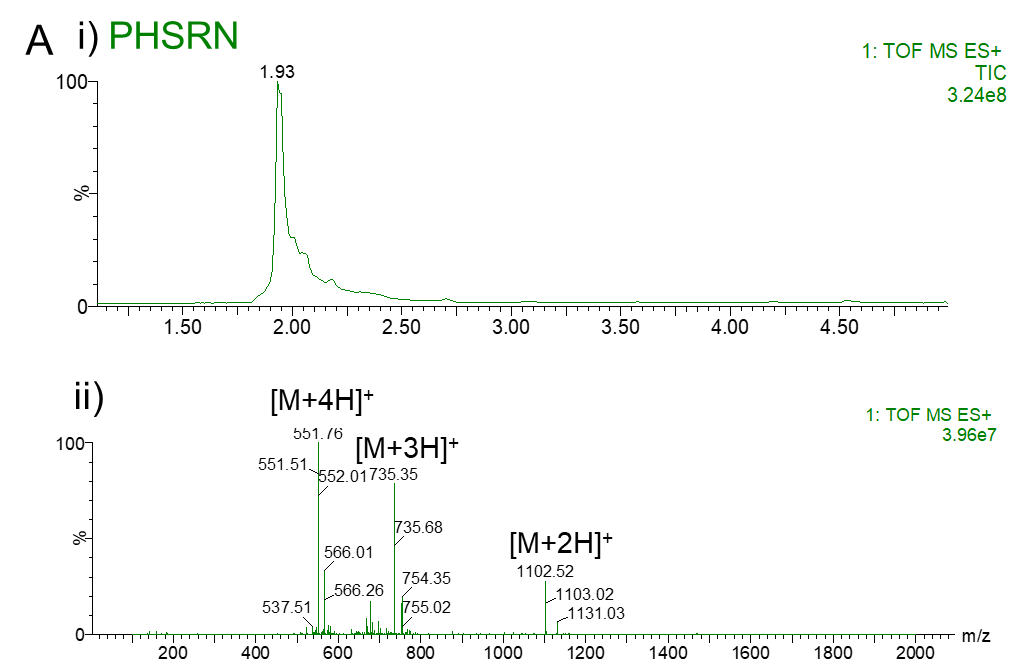

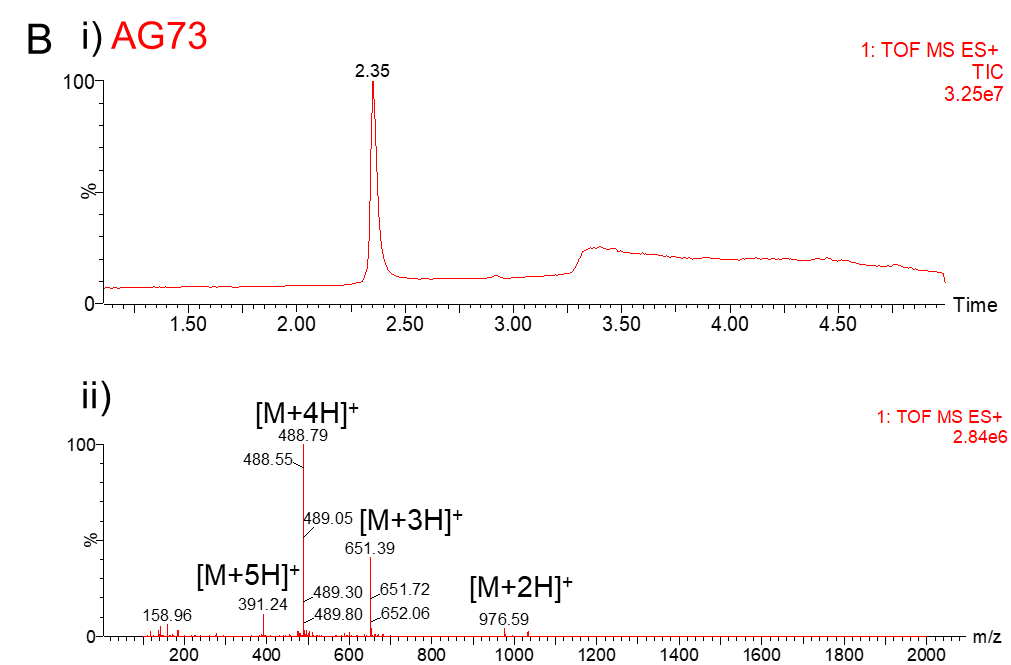

**Figure S28.** Representative mass spectrometry of bioactive peptides. A) PHSRN-RGDSP *i)* UPLC and *ii)* MS spectrum. B) AG73 *i)* UPLC and *ii)* MS spectrum.
